# Mitochondrial Energy Transformation Capacity Influences Brain Activation During Sensory, Affective, and Cognitive Tasks

**DOI:** 10.1101/2025.06.25.661482

**Authors:** Ke Bo, Catherine Kelly, Caroline Trumpff, Michio Hirano, Michel Thiebaut de Schotten, Martin Picard, Tor D. Wager

## Abstract

Mitochondria supply the energy required for brain function, but how energetic capacity constrains human brain activity remains unclear. We used genetically defined mitochondrial disease as a human model of impaired bioenergetics to test whether energetic limitations produce global disruption of brain function or selective vulnerability across certain functional domains. Participants completed resting-state fMRI and tasks probing working memory, sensory perception, and affective responses to cold pain, together with behavioral, clinical, and circulating biomarker assessments. We report impairment in working memory performance and reduced activation of associated brain networks in a subset of patients with the largest clinical and cognitive impairments. A resilient subgroup showed higher working-memory–related BOLD responses, which were associated with preserved task performance. Neural and behavioral effects scaled with the circulating marker of energetic stress growth differentiation factor 15 (GDF15). Brain activity related to sensory and affective responses and resting-state connectivity were comparable for patients and controls. These findings are consistent with an energy-hierarchy model and link clinical disease severity to working-memory brain responses and performance, while suggesting relative preservation of sensory and pain-related brain processes under energetic stress.

## Introduction

The human brain accounts for 20-24% of the body’s energy consumption while representing only about 2% of total body mass ^1–3^. This large and constant energy requirement underpins critical neural processes such as synaptic transmission, plasticity, and large-scale network coordination, which together support brain functions ranging from basic sensory perception to complex cognitive and affective regulation ^4^. While cellular and animal studies have long established the critical role of energy metabolism in neural function ^5,6^, recent human research increasingly highlights metabolic health as essential for human mental functioning ^7,8^.

Bioenergetic abnormalities are now recognized across diverse psychiatric and neurological disorders including major depressive disorder ^9^, psychotic disorders ^10^, autism spectrum disorders ^11^, bipolar disorder ^12^, as well as neurodegenerative diseases including Alzheimer disease ^13^ and Parkinson disease ^14^. Despite this growing evidence and the urgent need to elucidate specific neural mechanisms linking metabolism and brain function ^15^, directly translating these insights into human neuroscience remains challenging due to the limited availability of non-invasive methodologies for assessing or manipulating brain bioenergetics in vivo.

Mitochondria are central to cellular energy metabolism. They transform oxygen and food substrates into usable energy, and also serve as intracellular processors through dozens of interrelated functions ^16,17^. Mitochondria generate adenosine triphosphate (ATP) via oxidative phosphorylation, fueling the high demands of neurons and glial cells, powering intracellular processes such as ion transport, endocytosis, and neurotransmitter biosynthesis and reuptake, among others ^18^. Impaired mitochondrial energy transformation can disrupt synaptic transmission and ultimately compromise neuronal survival ^19^, alter gene expression ^20^ and neurogenesis ^21,22^, and therefore compromise brain functions. In terms of behavior, in animal models mitochondrial biology regulates anxiety ^23^, memory ^24^, dominance and social behaviors ^25,26^. Moreover, in humans, the brain contains distinct populations of mitochondria subserving a spectrum of energy demands and bioenergetic requirements, including more mitochondria specialized for energy transformation in metabolically-demanding areas serving more recently evolved brain areas involved in cognitive tasks ^27,28^. Thus, mitochondrial perturbations provide a unique window into understanding how brain functions and associated behaviors are shaped by cellular energy metabolism.

Mitochondrial diseases (MitoD) represent extreme cases of mitochondrial defects caused by mutations or deletions in the mitochondrial DNA (mtDNA). Among the most common mtDNA defects are the point mutation m.3243A>G in a transfer RNA gene, or single, large-scale deletions that commonly affect multiple mtDNA genes ^29,30^ (mutations in autosomal genes in the cell nucleus also cause MitoD ^31^). mtDNA defects cause functional abnormalities in mitochondria that decrease energy flux through the mitochondrial circuitry ^32^, impairing multiple mitochondrial functions. Consequently, affected individuals exhibit a spectrum of neurological symptoms, including cognitive impairment, psychiatric disturbances, and sensory deficits ^31,33–35^. Because mitochondrial defects directly compromise cellular energy metabolism, MitoD populations offer a natural opportunity to clarify the mind-mitochondria relationships. Comparing MitoD and individuals with healthy mitochondria along the axes of brain biology, cognition, and behavior, we can get at metabolism-function relationships, with the caveat that mitochondrial deficits drive systemic effects, likely associated with chronic developmental or acute compensatory adaptations. Despite the immense research potential of this population, prior brain imaging studies of MitoD have predominantly focused on structural and pathological abnormalities ^36–39^, leaving functional brain responses comparatively understudied. Functional neuroimaging studies involving multiple brain functions together with deep multi-modal phenotyping in this rare population remain scarce. The Mitochondrial Stress, Brain Imaging, and Epigenetics (MiSBIE) study ^40^ addresses this gap.

One key concept examined in this paper is the notion of energy constraints (EC). This refers to the notion that energy depletion or diversion can prevent normal or optimal functions, even in the absence of structural or functional alterations, and that energy can be depleted by disease-related processes that over-consume energy ^41^. This principle is based on fundamental bioenergetic constraints that operate across all levels of life ^42^. If a biological unit, a brain region, an activated cell type, or an organ, costs more energy than it should under optimal health conditions, it becomes “hypermetabolic” and steals energy away from other functions, a process called energy trade-offs ^43^. Consistent with this logic, mitochondrial defects like those causing MitoD produce cellular hypermetabolism in cells ^44^ and systemic hypermetabolism in affected patients ^45^. We hypothesize that this results in energy trade-offs relevant to health and cognition, although this question has not been directly addressed.

The current study provides, to our knowledge, the first systematic functional MRI (fMRI) investigation examining the relationship between bioenergetic deficits and brain function across multiple cognitive and functional domains in the MitoD population. This investigation draws on a rare, deeply phenotyped cohort of individuals with genetically defined mitochondrial disease, assembled through a 10-year effort at Columbia University. Our approach spans resting-state, sensory, affective, and cognitive domains using: (1) a multisensory checkerboard+tone task, (2) a cold pain task, and (3) a working memory (WM) task. We chose the WM task because it is both highly ATP-demanding^46^, making it sensitive to mitochondrial constraints, and central to executive control and fluid intelligence ^47,48^. Mitochondrial disease severity in MitoD was assessed using the Newcastle Mitochondrial Disease Adult Scale (NMDAS), the most widely validated, comprehensive, and disease-specific clinical severity scale in mitochondrial medicine ^49^, which captures overall mitochondrial disease burden rather than over-weighting any particular symptom domain. We used multiple models to extract spontaneous and task-related brain network activation and compare them between control participants and those with MitoD, as well as across patients with varying levels of disease severity.

We hypothesize that mitochondrial defects may manifest in two potential ways. First, the MitoD-related deficits in ATP production could lead to impaired synaptic activity ^50^, which might cause global impairments in brain functions, including sensory, cognitive, and affective domains, and spontaneous resting-state activity. If this hypothesis holds, we would expect MitoD patients to show reduced BOLD responses across all task conditions—multisensory, cold pain, and working memory—as well as reduced resting-state functional connectivity relative to controls.

Alternatively, due to energy-based competition driving energy constraints, some fundamental brain functions for sustaining life-critical processes might be indispensable, such as maintaining spontaneous brain activity and affective processing. Thus, the brain may preferentially allocate its limited energy resources to the essential operations^51,52^, leaving less energy available for functions that, while energy-intensive, are less fundamental, such as working memory ^52^. This second “competitive energy allocation” hypothesis predicts a selective pattern: preserved or enhanced brain responses in lower-demand sensory and affective tasks, with impairment restricted to the working memory domain and scaling with individual differences in disease severity. Here, we tested these competing hypotheses by comparing resting-state and task-related fMRI activity and connectivity between MitoD patients and healthy controls.

## Results

### Multidimensional characterization of the mitochondrial disease phenome

To provide an initial characterization of the MitoD phenome and identify affected and relatively preserved domains, we compared 70 healthy controls with 40 MitoD patients (91 of the participants have available MRI data; see *Methods* for details) across assessments of cognitive function, psychosocial variables including personality, mental health, mental and physical fatigability, molecular biomarkers derived from blood, daily behavior, disease related symptom severity, and clinical measures of neurological, physical, and cognitive function ^40^. Group differences were concentrated in selected cognitive, functional, and physiological measures, many with at least medium effect sizes (absolute Hedges’ g > 0.5; Figure 1A). Figure 1A summarizes selected measurements. The full battery, including measures without significant group differences and Benjamini–Hochberg false discovery rate-adjusted q values, is reported in Table S1. Molecular signaling factors in blood included large increases in Growth Differentiation Factor 15 (GDF15; Hedge’s g = 1.7) and Fibroblast Growth Factor 21 (FGF21; g = 1.34) in MitoD compared to control, both critical biomarkers indicative of impaired cellular metabolism ^53–55^. Additionally, MitoD patients showed elevated lactate (g = 1.05), glycosylated hemoglobin (g = 0.84), blood urea nitrogen (BUN, g = 0.69), elevated total lymphocyte count (g = 0.51), and decreased chloride levels (g = 0.6), consistent with mitochondrial metabolic stress and associated physiological alterations ^56^.

**Figure 1.**
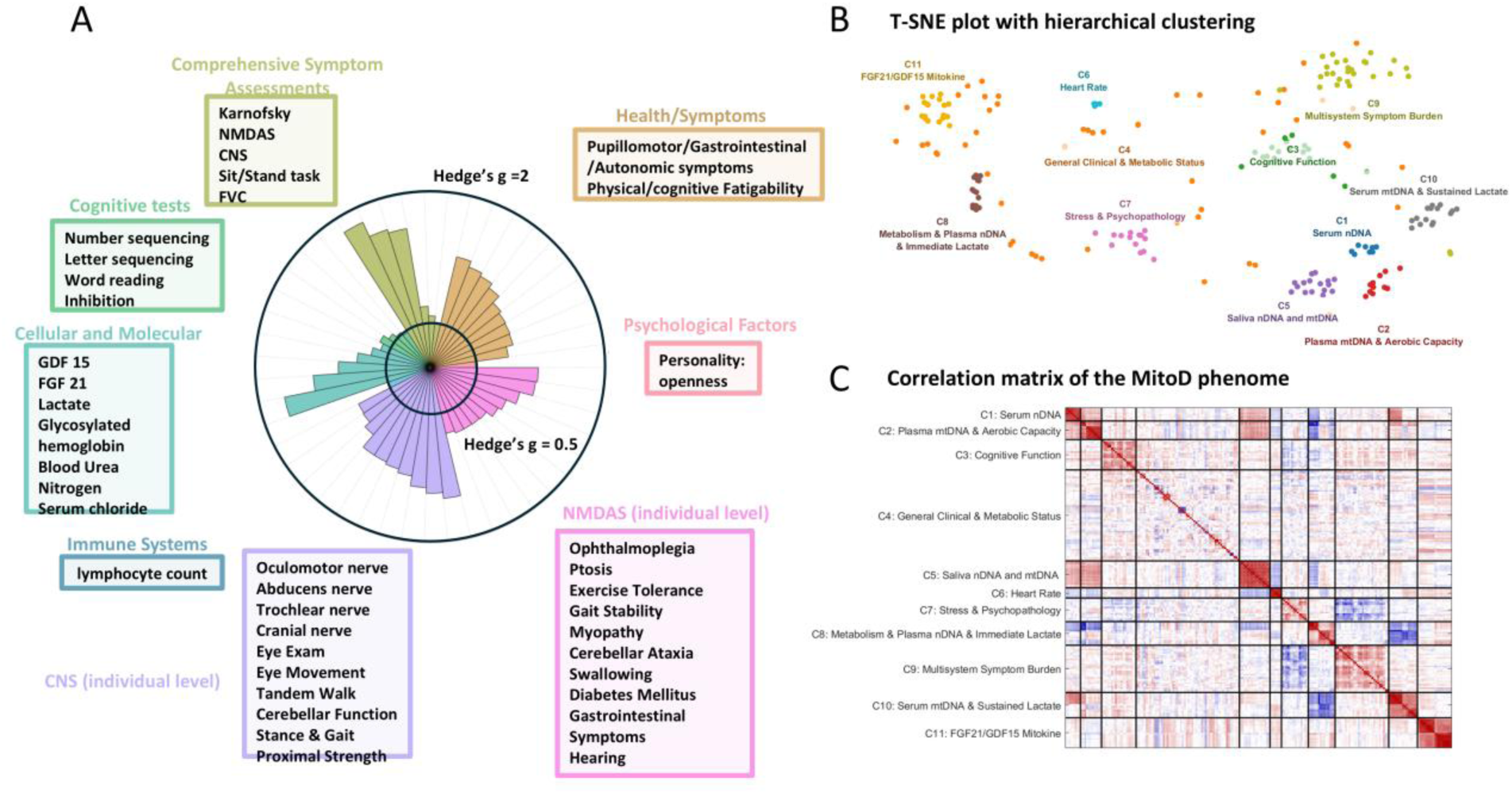
Overview of phenotypic domains and their interrelationships in the current study. **A)** Effect sizes (Hedges’ g) comparing mitochondrial disease (MitoD) patients to healthy controls across five key domains: cognitive tests, physical activity, autonomic symptoms, composite symptom assessments, and cellular/molecular markers. The wedge plot displays selected measurements rather than the full assessment battery. Complete results, including measures without significant group differences and FDR-adjusted q values, are provided in Table S1. Notably, two of the most representative comprehensive symptom assessments—the Newcastle Mitochondrial Disease Adult Scale (NMDAS), which evaluates mitochondrial disease–specific symptoms, and the Columbia Neurological Score (CNS), which evaluates general neurological health—are decomposed into single-item levels and displayed individually. The top ten individual items from each of these scales that survive FDR correction at q < 0.05 are also presented. **B)** t-SNE visualization of all measurements, with clusters defined by hierarchical clustering on partial Spearman correlations and color-coded accordingly. These clusters reflect eleven coherent biological and psychological domains, including mitokine stress responses, cell-free DNA dynamics across multiple biofluids, lactate kinetics, cognitive function, symptom burden, and general clinical measures. **C)** Partial Spearman correlation matrix of all measurements, with group membership (MitoD vs. control) statistically controlled to remove variance driven by group differences. Hierarchical clustering was used to sort the variables into eleven clusters. Red and blue indicate positive and negative partial correlations, respectively, highlighting distinct phenotypic clusters and cross-domain associations.

Compared to controls, MitoD patients also demonstrated differences on specific measures of processing speed, executive control, physical activity, and fatigability. On neuropsychological tests, patients performed worse on numeric sequencing, letter sequencing, word reading, and inhibition tasks (g = 0.64, 0.52, 0.58, and 0.48, respectively; trail-making and color-word interference tasks), and scored lower on openness in the personality test (g = 0.52, NEO Five-Factor Inventory). Patients further reported heightened physical (g = 1.23), cognitive (g = 1.00), and psychological fatigue (g = 0.99) assessed by the Modified Fatigue Impact Scale, which measures the impact of fatigue on physical, cognitive, and psychosocial functioning. Patients also reported pupillomotor symptoms (g = 1.30, COMPASS-31). These findings identify specific cognitive and functional vulnerabilities alongside the metabolic phenotype of MitoD.

Clinical assessments also identified greater disease burden and functional impairment in MitoD, including the Newcastle Mitochondrial Disease Adult Scale (NMDAS; g = 1.65; an index of adult mitochondrial disease–specific symptoms), the Columbia Neurological Score (CNS; g = 1.61; an index of neurological health across multiple domains), the Karnofsky Performance Scale (KPS; g = 1.83; an index of functional impairment), the Composite Autonomic Symptom Score (COMPASS; g = 1.01; an index of multi-system autonomic function), the sit–to–stand test (test of lower body power, balance, and endurance) (g = 0.70), and Forced Vital Capacity (FVC; g = 0.52; an index of lung function) (see *Methods* for detailed descriptions of these tests). Some of the largest single-item effects within these indices included impaired exercise tolerance (g = 1.07), ocular-function measures such as chronic progressive external ophthalmoplegia (g = 1.24) and ptosis (g = 1.22) from the NMDAS, and dysfunction in cranial nerves III, VI, and IV from the CNS (g = 1.51, 1.44, and 1.37, respectively), which regulate eye movement and pupil response, highlighting prominent ocular-motor involvement.

These differences did not extend across the full assessment battery. Measures of general cognitive ability, learning, verbal fluency, several personality dimensions, and anxiety showed little or no group difference (Table S1). Thus, Figure 1 identifies domains of vulnerability within a heterogeneous phenotype that also includes relatively preserved functions.

A summary of associations among these items across participants is shown in Figure 1B–C. To explore the functional grouping of these features, we first used a permutation test to determine the optimal number of clusters (^57^, see Methods for details). The permutation test identified two as the statistically optimal number of clusters (p < 0.05); however, to capture finer-grained phenotypic structure, we selected k = 11, which corresponded to a local maximum in the permutation-based pseudo-Z statistic. We then applied hierarchical clustering analysis using partial Spearman correlations, controlling for group membership to ensure that the observed co-variation among variables was not driven by group differences, which identified eleven distinct clusters (Figure 1B), each assigned a different color; the correlation matrix was sorted accordingly (Figure 1C). The eleven clusters captured separable biological and psychological domains: stress-test-evoked cell-free DNA release in serum, plasma, and saliva (Clusters 1, 2, and 5, respectively); immediate and sustained post-stress lactate responses co-occurring with resting metabolism and plasma DNA (Clusters 8 and 10); mitokine signaling reflecting mitochondrial stress, including FGF21 and GDF15 (Cluster 11); heart rate dynamics (Cluster 6); cognitive function (Cluster 3); personality and trauma history (Cluster 7); multisystem symptom burden including fatigue, depression, anxiety, and autonomic symptoms (Cluster 9); and a broad clinical and metabolic baseline (Cluster 4). In general, items from Cluster 9, encompassing negative psychosocial experience, fatigability, and autonomic symptoms, and

Cluster 11, encompassing mitokines GDF15 and FGF21, were negatively correlated with items in Cluster 3, including cognitive ability and neurological and physical health, reflecting an overall antagonistic relationship between physiological and psychological markers of metabolic deficits and resilience-promoting psychosocial and behavioral factors.

### Task-evoked brain activity

To probe brain function across cognitive, sensory, and affective domains, we modeled task-evoked hemodynamic responses relative to each task’s control condition. Each task induced robust, expected patterns of significant brain activation across MitoD and control participants (Benjamini–Hochberg FDR q < 0.05 across brain voxels; Figure 2A). The N-back contrast (2-back versus 0-back) recruited frontoparietal regions, striatum, cerebellum, midbrain, and visual cortices. Multisensory stimulation recruited early visual and auditory cortices and lateral prefrontal regions, whereas cold pain recruited the posterior and anterior insula, operculum, anterior midcingulate, striatum, cerebellum, and thalamus. The corresponding Neurosynth topic associations were consistent with these task domains.

**Figure 2.**
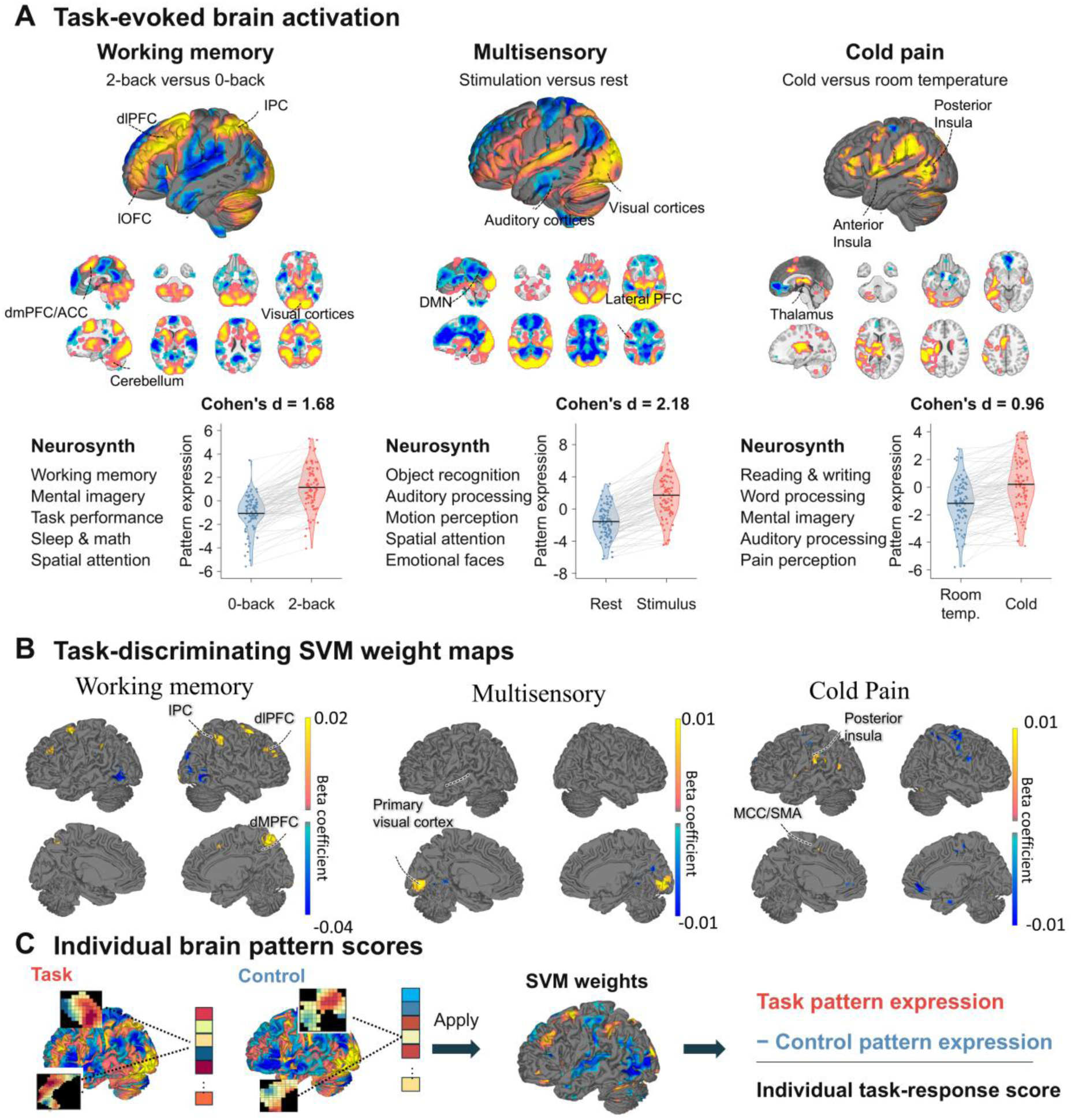
Task-evoked activation and individual brain pattern responses. **A)** Whole-brain activation maps for working memory (2-back versus 0-back), multisensory stimulation (visual checkerboard and audio versus rest), and cold pain (cold versus room-temperature arm wraps). Maps show FDR-corrected task effects (q < 0.05, corrected separately within each task); functional associations list the five most correlated Neurosynth forward-inference topic maps from 54 psychological topics ^58,59^. Paired violin plots show held-out brain pattern expression for both conditions in the final cross-validation repetition; the Results report mean effect sizes across repetitions. Dots represent participants (working memory, n = 88; multisensory, n = 90; cold pain, n = 91 before the display-only omission described below), connecting lines join conditions, and black bars indicate means. Cohen’s d denotes the paired effect d_z, calculated as the mean task-minus-control difference divided by its standard deviation: 1.68, 2.18, and 0.96. For display only, three cold-pain pairs with either condition exceeding three scaled median absolute deviations from the condition median are omitted; all pairs remain in the effect-size calculation and statistical analyses. **B)** Task-specific SVM weight maps evaluated using 5,000 bootstrap samples.

Thresholds are FDR q < 0.05 for working memory and multisensory processing and uncorrected p < 0.001 for cold pain; full, unthresholded maps were used for score calculation. **C)** Projection of task and control maps onto SVM weights yields condition-specific pattern-expression scores; their difference defines the task-related brain pattern expression. PFC, prefrontal cortex; IPC, inferior parietal cortex; ACC, anterior cingulate cortex; MCC, midcingulate cortex; SMA, supplementary motor area; DMN, default mode network. See also Figures S1 and S2.

We trained paired support vector machine (SVM) classifiers to distinguish task from control states, using 5-fold cross-validation. As shown in Figure 2A, classification accuracies were 97.0% for N-back (2-back versus 0-back, d = 1.67), 98.24% for multisensory stimulation versus rest (d = 2.18), and 87.12% for cold versus room temperature (d = 0.92). These large effects in held-out participants demonstrate robust task-related brain activation across all three paradigms. The paired condition responses in Figure 2A illustrate these effects at the individual-participant level, with most participants showing greater brain pattern expression in the task condition than in the corresponding control condition. Figure 2B shows the task-specific weight maps (see Figure S1 for uncorrected maps) and Figure 2C illustrates how the task-minus-control difference in pattern expression yields task-related brain activation at the individual level. These brain pattern expression scores provided the measures used in the subsequent brain–behavior and severity analyses ^60–62^.

### Working memory brain activation and disease severity

Alongside the affected and relatively preserved functions identified in Figure 1, disease severity varied substantially within the MitoD population. Some patients showed mild or no typical MitoD symptoms, whereas others had severe symptoms affecting daily life. We assessed this variation using NMDAS, the most widely validated, clinician-administered composite scale in mitochondrial medicine, designed to capture overall disease burden across all organ systems rather than any single symptom domain ^49^. The NMDAS distribution in the N-back patient sample appeared bimodal, with a mild (resilient) subgroup and a severe subgroup (Figure 3A). We used K-means clustering to identify these subgroups in a data-driven way. Similar distributions were observed in the multisensory and cold-pain patient samples (Figure S4). This heterogeneity motivated examining how disease severity relates to brain activation and behavioral performance within the MitoD population. We used Spearman’s rank correlations for NMDAS because of its non-normal distribution, and bootstrap inference for mediation analyses.

**Figure 3.**
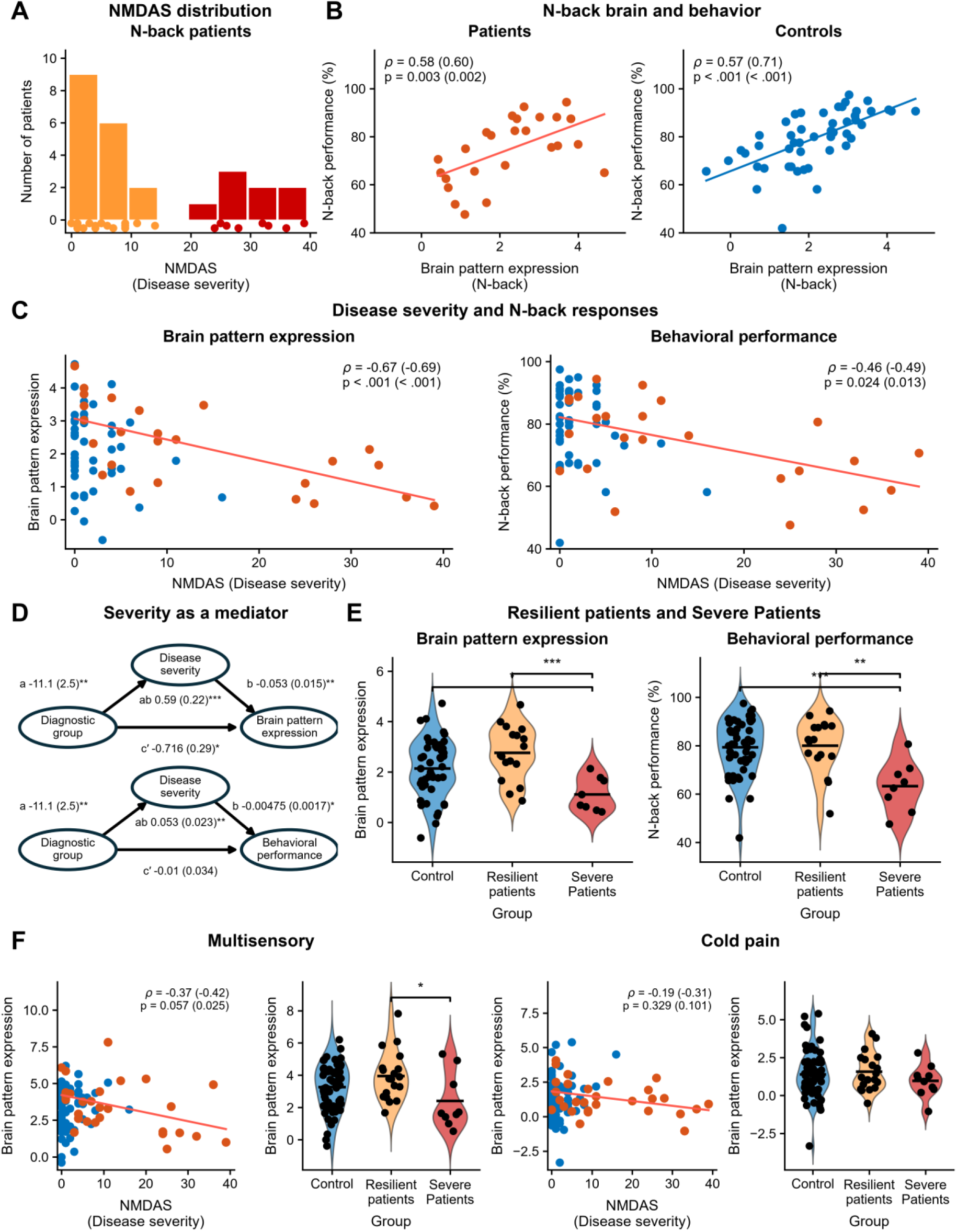
The influence of mitochondrial disease severity on brain activation and behavioral performance. A) NMDAS distribution in N-back patients. Amber and red denote the resilient and severe patient subgroups identified using K-means clustering. B) Correlations between N-back brain pattern expression and task performance in patients (n = 25) and controls (n = 49). C) Correlations between NMDAS and N-back brain pattern expression (left) and task performance (right) within patients. In B, C, and F, age-adjusted partial Spearman correlations and p values are shown first, with unadjusted values in parentheses. Points show individual observations; lines show linear fits. Controls on the NMDAS axes are included for visual context only. D) Mediation analyses testing whether disease severity mediates the relationship between MitoD status and N-back brain pattern expression (top) or task performance (bottom). Models include 72 complete cases and control for age (control = 1, MitoD = 0); coefficients are unstandardized, and accuracy is expressed as a proportion. Parentheses indicate bootstrap standard errors. E) N-back brain pattern expression and task performance in controls, resilient patients, and severe patients. F) Corresponding multisensory (28 patients, 62 controls) and cold-pain (29 patients, 62 controls) analyses. Dots represent participants and horizontal bars indicate means. Blue, controls; orange, MitoD; amber, resilient patients; red, severe patients. *p < 0.05, **p < 0.01, ***p < 0.001 (two-sided t tests for group comparisons; bootstrap inference for mediation). See also Figures S3–S8.

The N-back task provided both behavioral performance and fMRI responses. We first tested whether N-back brain pattern expression tracked behavioral performance. Higher brain pattern expression was associated with greater accuracy in both patients (age-adjusted partial Spearman ρ = 0.58, p = 0.003) and controls (ρ = 0.57, p < 0.001; Figure 3B), supporting the behavioral relevance of this brain measurement.

Within patients, higher NMDAS severity was associated with lower N-back brain pattern expression (age-adjusted partial Spearman ρ = −0.67, p < 0.001, 95% CI [−0.845, −0.366], partial R² = 0.449) and poorer behavioral performance (ρ = −0.46, p = 0.024, 95% CI [−0.728, −0.069], partial R² = 0.211; Figure 3C). Both associations survived Benjamini–Hochberg correction across the four primary severity outcomes (q = 0.0014 and 0.048, respectively). We used rank correlations because of the non-normal NMDAS distribution and treated adjustment for age as the primary analysis; unadjusted results are shown in parentheses in Figure 3. A split-half robustness analysis (1,000 iterations) further supported the reported associations across subsets of the data (brain: median r = −0.646, 95% CI [−0.762, −0.474]; behavior: median r = −0.453, 95% CI [−0.589, −0.277]; 100% directional consistency for both; see Methods). Together, these findings indicate that greater disease severity is associated with reduced working-memory network activation and poorer task performance.

To further characterize the NMDAS–brain and behavioral associations, we compared the resilient and severe patient subgroups with controls (Figure 3E). A data-driven K-means analysis identified the boundary between mild (resilient) and severe patients. Bootstrap resampling confirmed stable assignments (>99.9% across 10,000 resamples), and the subgroups did not differ significantly in age (t(23) = 0.97, p = 0.34), sex (Fisher’s exact test, p = 0.36), or genotype distribution (χ²(2) = 4.61, p = 0.10; Figure S8). Resilient patients showed comparable behavioral performance to controls (t(64) = 0.23, p = 0.815) and numerically higher working memory-related brain activation (t(64) = 1.97, p = 0.053). In contrast, severe patients showed reduced brain activation (t(55) = −2.47, p = 0.017) and poorer performance (t(55) = −3.71, p < 0.001). This pattern is consistent with a potential compensatory mechanism whereby resilient MitoD patients recruit greater neural activity to maintain performance comparable to controls, while severe patients show reduced activation and impaired performance.

Sensitivity analyses further evaluated the disease severity–brain and behavioral associations after additionally controlling for sex, genotype/variant class, fatigue, and vision-related neurological measures (see Supplementary Results and Tables S4 and S5). No participants were receiving disease-specific pharmacological treatments. Psychiatric comorbidities were assessed using the Beck Depression Inventory, State-Trait Anxiety Inventory, and DSM-5 personality scales; phenome-wide associations for these measures are reported in Supplementary Table S2. The primary NMDAS–brain activation association remained significant after controlling for psychiatric symptoms (Supplementary Table S4). To assess potential structural brain damage in MitoD ^63,64^ as a confound, supplementary analyses confirmed that the brain–NMDAS association remained significant after controlling for structural measures including cortical thickness, white matter integrity, and NODDI microstructural indices (Supplementary Results).

Disease duration was not included as a covariate because pathogenic mtDNA variants are present from birth in genetic mitochondrial disorders, while clinical onset, diagnosis, and symptom progression occur at highly variable ages, making a single disease-duration metric ill-defined and potentially misleading.

### Multisensory and cold-pain responses

We next examined whether the disease severity–brain association extended to the multisensory and cold pain tasks (Figure 3F). In the multisensory task, a similar but weaker association was observed after controlling for age (partial Spearman ρ = −0.37, p = 0.057, 95% CI [−0.658, 0.011]). In contrast, the cold pain task showed no significant association with disease severity (partial Spearman ρ = −0.19, p = 0.329, 95% CI [−0.527, 0.196]) or differences between patient subgroups (Figure 3F, right). Behavioral performance was not collected for these tasks. We also did not find strong or widespread alterations in spontaneous brain activity or network-level connectivity at rest (Figures S5–S6).

In the corresponding multisensory subgroup analysis, resilient patients showed higher brain pattern expression than severe patients (t(26) = 2.42, p = 0.023, 95% CI [0.235, 2.861]). Relative to controls, resilient patients showed a trend toward higher brain pattern expression (t(79) = 1.82, p = 0.072, 95% CI [−0.062, 1.430]), whereas severe patients showed numerically lower expression (t(69) = −1.67, p = 0.099, 95% CI [−1.895, 0.167]; Figure 3F, left).

Finally, Bayes factors for the age-adjusted severity–brain associations supported a non-zero association for working memory (BF₁₀ = 89.54), were inconclusive for multisensory processing (BF₁₀ = 0.88), and provided moderate evidence for a null association for cold pain (BF₁₀ = 0.23).

Overall, these findings point to selective effects of disease severity on working memory, and are consistent with the notion of a hierarchical prioritization of function under energetic constraints ^41–43^: Affective processing, as indexed by the cold pain task, showed no significant BOLD differences with different levels of mitochondrial disease severity, consistent with the notion that survival-critical functions are prioritized under energetic constraints. In contrast, working memory, a high-energy-demand but less survival-critical function, showed the strongest severity-related reductions in neural engagement. Broader phenome-wide associations with working-memory brain activation and behavioral performance are reported in Supplementary Tables S2 and S3.

## Spatial correspondence with cortical mitochondrial maps

To relate the NMDAS findings to the brain’s energetic architecture, we compared task-evoked cortical activation with normative MitoBrainMap measures^27,65^. Complex I, II, and IV maps index the activity of respiratory-chain enzymes supporting oxidative phosphorylation, whereas mitochondrial density indexes mitochondrial abundance. Tissue respiratory capacity (TRC) combines respiratory-chain enzyme activities at the tissue level; mitochondrial respiratory capacity (MRC) expresses TRC relative to mitochondrial density, indexing capacity per unit of mitochondria. Working-memory activation correlated positively with complex I (r = 0.40, p = 0.004, q = 0.023), complex II (r = 0.31, p = 0.019, q = 0.038), and mitochondrial density (r = 0.33, p = 0.012, q = 0.036; Figure 4). These associations survived cortical spin testing with FDR correction across six mitochondrial maps within each task. The remaining working-memory associations and all multisensory and cold-pain associations did not reach FDR-corrected significance. This spatial correspondence provides converging anatomical support for the NMDAS findings by linking working-memory recruitment to regional mitochondrial abundance and respiratory-chain activity.

**Figure 4.**
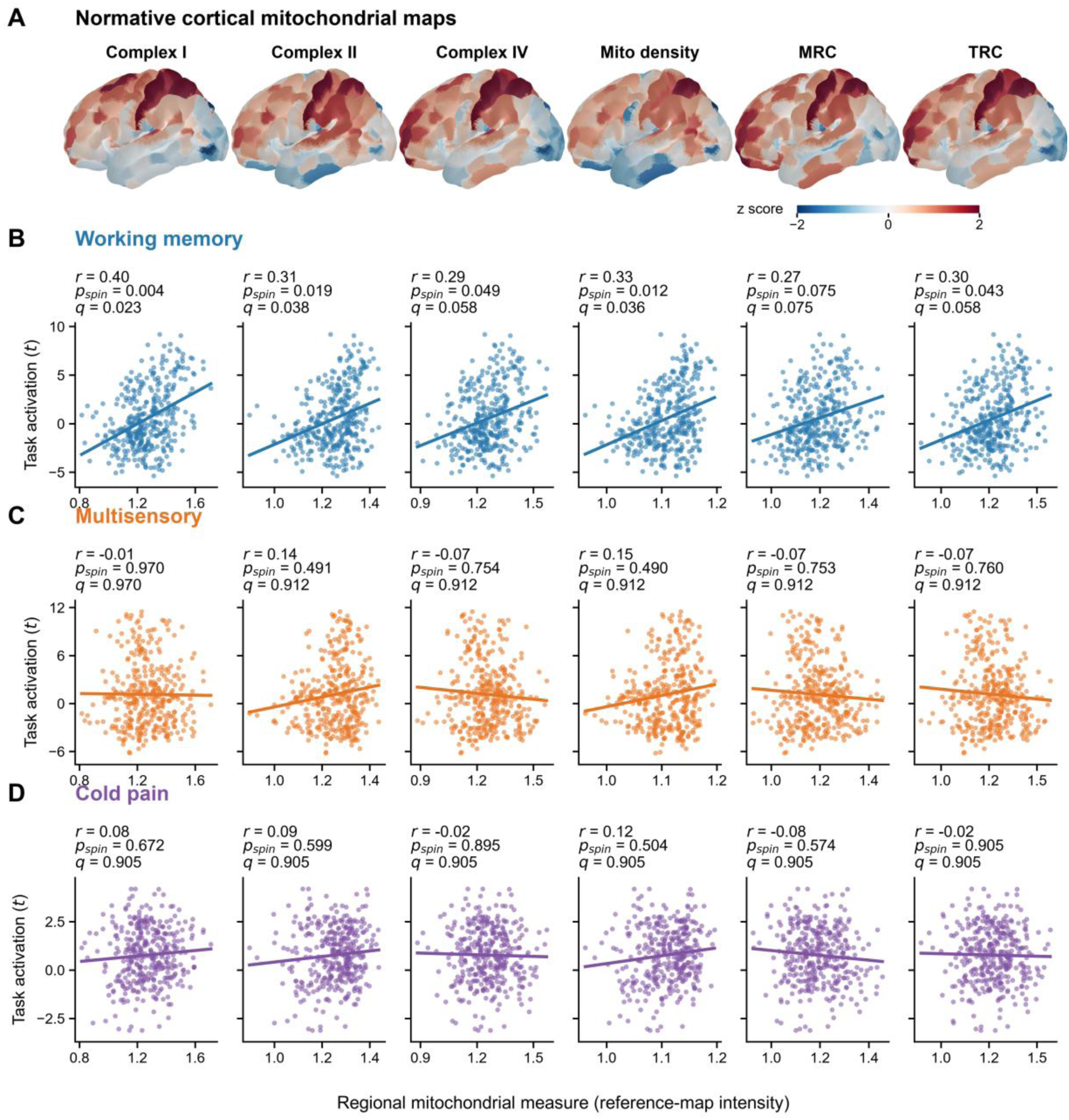
Spatial correspondence between task-evoked activation and normative cortical mitochondrial maps. A) Mitochondrial maps rendered from the published regional values^27,65^ on the left lateral cortical surface. Colors show z scores across all 360 parcels for display only, with the color scale limited to ±2. Columns show complexes I, II, and IV, mitochondrial density, mitochondrial respiratory capacity (MRC), and tissue respiratory capacity (TRC). B–D) Spatial correlations for working memory (blue), multisensory stimulation (orange), and cold pain (purple). Each point represents one of 360 HCP-MMP1 parcels across both hemispheres. Task values are parcel-averaged, unthresholded voxelwise t statistics (88, 90, and 91 participants, respectively); mitochondrial values retain the original reference-map intensity. Lines show linear fits. Labels report Pearson’s r, two-sided cortical-spin p, and FDR-adjusted q (20,000 rotations; six comparisons per task). Mitochondrial maps are normative references obtained independently of the MiSBIE participants.

### Overall group comparisons and possible compensation

We then compared task-related brain activation between MitoD patients and controls. At the available group sizes, sensitivity analyses indicated 80% power to detect between-group differences of Cohen’s d = 0.70 for working memory and d = 0.64 for both multisensory stimulation and cold pain (two-sided α = 0.05; see Methods). None of the three tasks showed a significant overall group difference: working memory, t(72) = 0.33, p = 0.741; multisensory, t(88) = 0.54, p = 0.589; and cold pain, t(89) = −0.12, p = 0.907. N-back behavioral performance also did not differ significantly, t(72) = −1.55, p = 0.126 (Figure S7). The absence of a categorical group difference may reflect the wide clinical heterogeneity within MitoD, with greater neural engagement in resilient patients and reduced activation in severe patients averaging to similar overall responses.

Age-adjusted mediation analyses further examined the role of disease severity (Figure 3D). The indirect effects through NMDAS were positive for N-back brain expression (ab = 0.59, bootstrap SE = 0.22) and accuracy (ab = 0.053, SE = 0.023; both bootstrap p < 0.01). The direct group coefficient remained significant for brain expression (c′ = −0.716, SE = 0.29, p < 0.05) but not for accuracy (c′ = −0.010, SE = 0.034). These opposing direct and indirect associations help describe how severity heterogeneity can coexist with a small overall group difference, consistent with the proposed compensatory mechanism in resilient MitoD patients.

To ensure that group differences in head motion did not account for the reported effects, we compared mean framewise displacement (FD) and DVARS between MitoD patients and controls for each task. No significant group differences were observed (Figure S2).

### The energetic stress marker GDF15 and brain function

Functional measures of MitoD disease severity may be paralleled by systemic processes related to mitochondrial function and cellular metabolism detectable in blood and other tissues. If so, blood or salivary biomarkers related to these processes could provide indices of systemic mitochondrial health in MitoD and even beyond, as well as targets for prevention and treatment efforts. Among potential circulating biomarkers, GDF15 is the most robust marker of MitoD ^56,66^. GDF15 also is a stress-responsive cytokine increasingly recognized as a marker of cellular and mitochondrial energetic stress ^67^. Blood GDF15 meets several key criteria for a marker of systemic energetic stress: (1) it is the most dramatically elevated circulating protein with aging; (2) it is induced by mitochondrial-targeting drugs including metformin; and (3) its levels increase in many chronic conditions—including cardiovascular disease, cancer, mood disorders, Alzheimer’s disease, and autoimmune disorders ^67,68^.

In the current study, plasma GDF15 was substantially higher in MitoD patients than controls (t = 6.1, p < 0.001; 95% CI [1.03, 2.24] see Figure 5A) and strongly correlated with NMDAS disease severity in MitoD (r = 0.75, p < 0.001;R² = 0.562, 95% CI [0.47, 0.89]), replicating previous findings using the MiSBIE dataset^69^ (reproduced analysis). The effect sizes were comparable to those of composite clinical batteries like the NMDAS and CNS and large enough to separate patients from controls, achieving an area under the ROC curve of >0.85 ^69^. These findings and others reviewed elsewhere ^32^ suggest GDF15 has the potential to serve as a broader index of mitochondrial energetic stress that can be applied in both control and MitoD populations.

**Figure 5.**
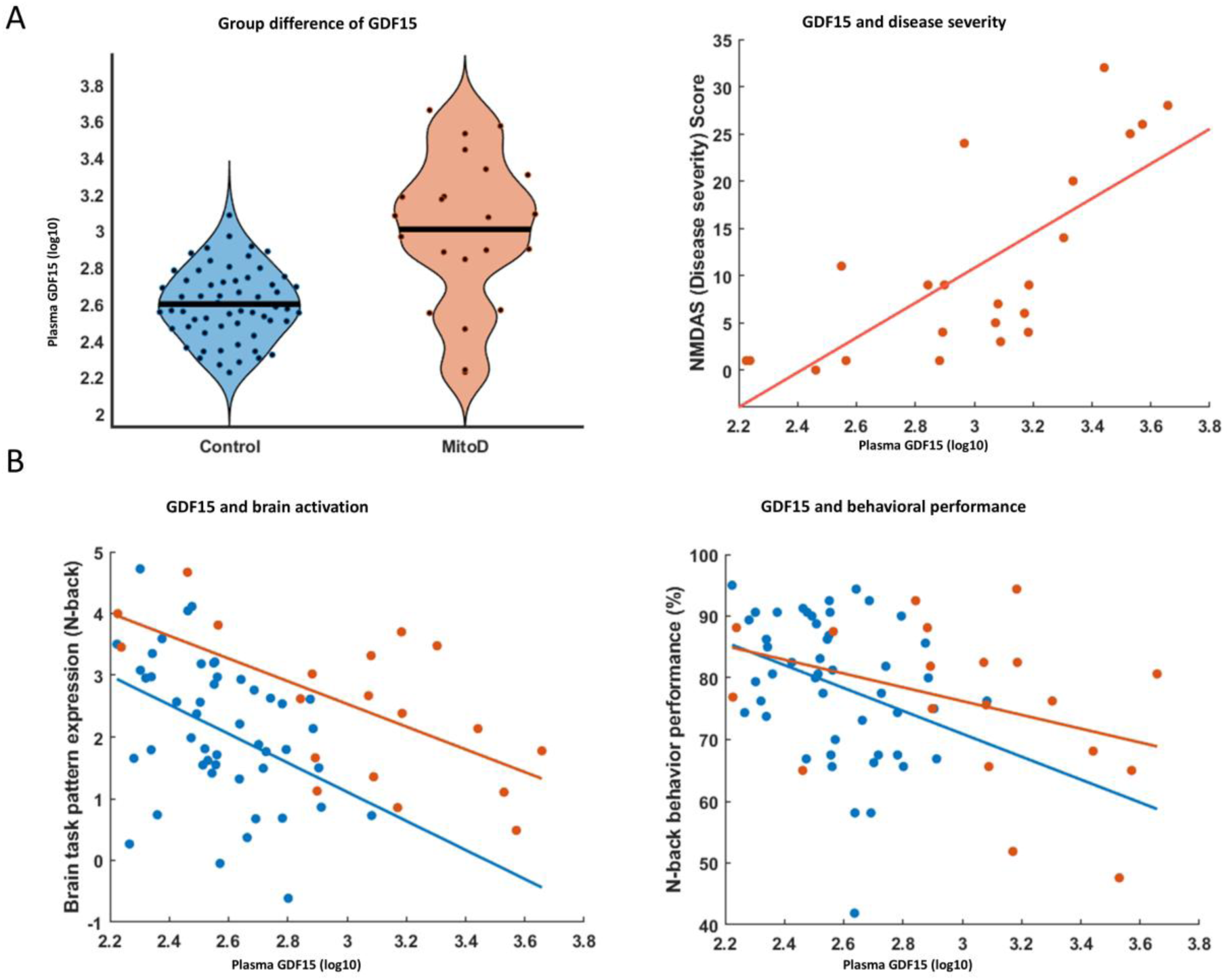
GDF15 as a potential biomarker linking mitochondrial bioenergetic dysfunction to cognitive deficits tested by brain and behavioral measurements. **A)** Validation and reproduced analyses demonstrate that plasma GDF15 (log-transformed) robustly differentiates the MitoD population from normal controls and is significantly associated with NMDAS disease severity within the MitoD group, replicating findings published in a previous study using the same dataset^69^ **B)** Relationships between plasma GDF15 and working-memory outcomes are shown in two panels. Left panel: Correlation (Pearson) with brain-pattern expression during the N-back task—whole group: r = -0.48, p < 0.001; MitoD: r = -0.64, p = 0.003; Control: r = -0.41, p = 0.004. Right panel: Correlation with behavioral performance— whole group: r = -0.33, p = 0.007; MitoD: r = -0.36, p = 0.12; Control: r = -0.33, p = 0.02. Data points are colored blue for controls and orange for MitoD patients.

Unlike disease severity—which is only relevant to MitoD patients—GDF15 is non-zero in controls and reflects their relative mitochondrial energetic stress level. Therefore, we tested whether GDF15 levels were associated with working memory–related brain activity and task performance across the entire sample. As shown in Figure 5B, higher GDF15 levels predicted reduced N-back brain pattern expression across the entire sample (r = –0.48, p < 0.001, partial R^2^ = 0.235, 95% CI [-0.65, -0.28]), controlling for Group (MitoD vs. control). Within each group, this relationship remained significant (patients: r = –0.64, p = 0.003, partial R2 = 0.408, 95% CI [-0.85, -0.26]; controls: r = –0.41, p = 0.004, partial R^2^ = 0.166, 95% CI [-0.62, -0.14]).

Similarly, higher GDF15 levels predicted poorer behavioral performance on the N-back task across the whole cohort (r = –0.33, p = 0.007, partial R^2^ = 0.108, 95% CI [-0.53, -0.10]), controlling for Group. GDF15 is produced by non-brain tissues and signals, possibly exclusively, through the brainstem to alter behavior ^70,71^, consistent with the notion that peripheral signals of energetic stress may suppress energetically expensive brain functions ^72^. The effect sizes were comparable in controls (r = –0.33, p = 0.02, partial R^2^ = 0.106, 95% CI [-0.56, -0.05]) and in

MitoD patients (r = –0.36, p = 0.12, partial R^2^ = 0.132, 95% CI [-0.70, 0.11]), although the latter did not reach significance, likely due to its smaller sample size. Together, these results highlight GDF15’s potential as an effective biomarker of metabolic health relevant for tracking brain activity and cognitive performance during an energy-demanding task.

GDF15 is highly related to age in humans where it increases exponentially with advancing age ^73–75^. In the current data, Age was not correlated with Group (r = -0.016, p = 0.89), but was correlated with higher GDF15 (r = 0.38, p = 0.001). Therefore, we further tested whether the GDF15 is able to predict working memory brain/behavior performance independently from the age effect. We tested two multiple regression models. Model 1 considered the main effects of Group (Control = 1, MitoD = -1), GDF15 (Log 10), and Age, and Model 2 tested a full model with their interactions.

In models predicting working memory–related brain activation, we found significant main effects of GDF15, age, and group in Model 1 (t = –2.82, p = 0.006; t = –2.75, p = 0.008; t = –2.95, p = 0.004, respectively; df = 63). No interaction effects in Model 2 reached significance. These results indicate that, while aging is associated with reduced brain activation, higher levels of GDF15 account for additional variance in working memory–related activity, independent of age. Thus, metabolic health, as indexed by GDF15, may influence brain function independently from chronological aging.

In models predicting working memory–related behavioral performance, we found that only Age showed a significant main effect in Model 1 (t = – 3.22, p = 0.002, df = 63). In Model 2, two interaction terms were significant: Age × Group (t = –3.01, p = 0.004, df = 59) and GDF15 × Group (t = 2.02, p = 0.048, df = 59). These results suggest two key points: 1) GDF15 does not predict behavioral performance independently of age across the full sample. 2) Both GDF15 and Age showed negative associations with behavioral performance overall, but their effects differ by group. The significant interactions indicate that in MitoD patients, compared to controls, GDF15 is stronger in predicting performance, whereas age is less influential. This suggests that factors beyond age, such as genetic influences, may play an additional role in shaping metabolic health and cognitive behavior in the patient group.

In sum, GDF15 emerges as a robust marker of systemic energetic stress that predicts brain activation and behavior during cognitively demanding tasks across the whole sample. Its association with brain activation appears only partly driven by age-related metabolic decline, while its relationship with behavioral performance is more strongly influenced by the aging process.

## Discussion

In this study, genetically defined mitochondrial disease was associated with a selective profile of clinical, cognitive, autonomic, and circulating biomarker differences, alongside relatively preserved measures across the phenome. The imaging findings similarly showed no widespread differences in task-evoked or resting-state fMRI measures. Instead, clinical severity was most strongly associated with working-memory activation and performance. A resilient subgroup maintained performance and showed numerically higher working-memory-related BOLD responses than controls. Multisensory activation showed a weaker association, whereas no significant cold-pain association or widespread resting-state differences were detected. The spatial alignment of working-memory activation with normative mitochondrial density and complex I and II activity provides complementary anatomical evidence, supporting an energetic interpretation of the working-memory findings. GDF15, a circulating marker of mitochondrial stress, was associated with working-memory activation in both patients and controls and explained variance beyond age.

Together, the phenome-wide associations with genetic mitochondrial disease and task-based neurocognitive findings are consistent with domain-specific functional vulnerability in working memory rather than uniform disruption across the brain systems examined. The full phenome assessment showed that many measures of general cognitive ability, learning, fluency, recall, personality, and negative affect differed little between groups, whereas larger deficits were concentrated in cognitive functions dependent on working memory, including sequencing, inhibition, and executive function (Supplementary Table S1). This pattern is consistent with prior evidence that adult MitoD produces focal rather than global cognitive deficits^35^.

Two nonexclusive mechanisms may explain why complex cognition is preferentially affected. First, working memory requires recurrent, distributed network activity and may impose greater energetic demands than simpler functions^46,76–78^. Intrinsic, spontaneous activity already consumes the majority of the brain’s energy budget (∼60–80%)^79^, while task engagement typically requires only a modest “top-up” of oxidative metabolism, on the order of ∼5% above resting levels^80^. In a healthy brain, this additional cost can be covered by available metabolic reserve. In MitoD, however, hypermetabolic stress responses may reduce that reserve^43–45^, so even a modest incremental demand can become limiting, causing high-demand functions such as working memory to falter first. Second, chronic bioenergetic constraints may promote adaptive neural reallocation toward essential or lower-cost functions. In healthy individuals, increasing attentional demand can shift metabolic resources toward attended visual information and away from unattended information^52^, demonstrating that energy allocation can be dynamically redistributed. By extension, under chronic MitoD the brain may preserve pain, basic sensorimotor processing, and spontaneous activity while leaving fewer resources available for complex cognitive operations. Together, these mechanisms provide a plausible basis for the hypothesized energy hierarchy: selective vulnerability may arise from both reduced energetic reserve and how limited resources are allocated across functions.

Several factors warrant consideration when interpreting the neuroimaging findings. First, long-term mitochondrial respiratory chain deficits in MitoD may drive developmental or experience-dependent adaptations in neural circuitry^81,82^, such that the observed BOLD responses may partly reflect the accumulated effects of compensatory plasticity rather than the immediate consequences of energetic limitation alone^83^. Second, mitochondrial disease may also influence neurovascular coupling through endothelial and vascular processes that shape the relationship between neural activity and the resulting hemodynamic response ^84,85^. Future studies combining fMRI with more direct measures of cerebral metabolism, such as MR spectroscopy, PET, or cerebrovascular reactivity mapping, could help distinguish among these potential contributions.

GDF15 provided a complementary blood-based link between systemic energetic stress and cognitive brain function. It was markedly elevated in MitoD, tracked disease severity, and was associated with lower working-memory brain pattern expression in both patients and controls, extending its relevance beyond rare genetic disease^56,66^. GDF15 is also elevated across psychiatric conditions^86–88^ and has been associated with cognitive decline^89,90^, raising the possibility that systemic energetic stress contributes to neural function during effortful cognition and resource allocation^51,91^. In the current study, GDF15 increased with age^75^ but explained additional variance in working-memory brain activity beyond chronological age. This residual variation may reflect modifiable influences on metabolic and cognitive health, including physical activity, education, and diet^92–94^, as well as age-independent genetic and environmental influences in MitoD^95–98^. These findings open new avenues for understanding the effects of mitochondrial disease on the brain and cognition and understanding energetic constraints on cognition more broadly. In addition, our findings suggest that working memory—and executive functions that rely on it—is energetically constrained, and that blood biomarkers such as GDF15 may provide important insights into cognitive capacity limitations across clinical and nonclinical populations.

### Limitations

Several limitations should be considered. First, the imaging sample was modest, particularly for within-patient and subgroup analyses, and smaller effects may therefore have gone undetected. Second, while the task battery sampled cognitive, sensory, and affective domains, it does not encompass the full range of energetically demanding brain functions. Studies incorporating additional processes, such as sustained attention, language, and emotion regulation, will be important for testing the generality of the energy-hierarchy framework. Finally, although we statistically controlled for age, the strong correlation between GDF15 levels and age in healthy controls underscores the challenge of disentangling aging effects from mitochondrial health when interpreting metabolic biomarkers. Longitudinal designs and complementary measures of mitochondrial biology will be needed to clarify the independent contributions of chronological aging and energetic deficits.

### Conclusions

In summary, genetic mitochondrial disease was associated with elevated blood levels of GDF15 and other metabolic markers, specific cognitive impairments in sequencing and inhibition, and increased autonomic symptoms and fatigue. These differences coexisted with relatively preserved measures across the phenome, indicating a selective rather than global pattern of impairment. In line with this pattern, overall brain BOLD activity and connectivity were largely intact across multiple tasks and connectivity metrics, but individuals with greater cognitive impairment and higher GDF15 levels exhibited reduced working memory-related BOLD activity, which in turn was associated with poorer task performance. Conversely, high-functioning MitoD individuals showed higher BOLD responses in working memory regions that supported relatively intact performance. This represents a resilient phenotype that could be further studied to identify resilience-related factors. Disease severity was associated with working-memory brain responses and performance; the age-adjusted multisensory and cold-pain associations did not reach statistical significance. This pattern points to a potential energy hierarchy in the brain that prioritizes essential functions under energy constraints ^41^. Together, these results advance our understanding of how intrinsic mitochondrial energy deficits may reshape neural activity and cognitive outcomes. They also underscore the potential of metabolic biomarkers like GDF15 to inform future interventions targeting brain energy metabolism across both rare mitochondrial diseases and more common neurodegenerative conditions.

## Methods

### Participants

This study was conducted at Columbia University from 2018 through 2023 under institutional review board approval (IRB protocol #7424). All participants provided written informed consent. English-speaking adults aged 18 to 60 years with genetically confirmed primary mitochondrial disease were recruited through clinical and research networks. Controls without mitochondrial disease were frequency matched on age and sex (Supplementary Figure S9). Exclusion criteria included recent acute illness, pregnancy, current cancer, steroid treatment, participation in another clinical trial, and substantial cognitive impairment^40^. The study was registered with ClinicalTrials.gov (NCT04831424).

The full cohort included 40 patients with MitoD and 70 controls. Ninety-one participants completed MRI. Task-specific imaging data were available for 88 participants for working memory, 90 for multisensory processing, and 91 for cold pain. Fourteen participants were further excluded from the primary N-back working-memory analyses based on a prespecified behavioral data-quality/task-engagement criterion: unavailable or severely incomplete behavioral response data (n = 12) or usable response logs with accuracy below 40% (n = 2). Of these 14 exclusions, 11 were healthy controls and 3 were MitoD patients. This criterion was used to exclude uninterpretable task engagement rather than low-performing participants, was applied identically across groups, and had no systematic relationship to disease severity. Analyses retaining these participants yielded similar conclusions and are reported in Supplementary Figure S3.

### Materials and procedures

Participants completed a broad battery of neuropsychological, clinical, psychosocial, autonomic, physical, physiological, and blood-based assessments^40^. The primary clinical severity measure was the NMDAS, a clinician-administered composite assessing multisystem mitochondrial disease burden. Plasma GDF15 was selected as the primary circulating marker because it is strongly elevated in mitochondrial disease and reflects cellular stress^56,66,67^. GDF15 values were log10 transformed because of right skew. Working-memory performance was defined as the percentage of correct responses during the N-back task.

### MRI experimental paradigm and preprocessing

Imaging was performed on a 3-T Siemens Prisma scanner and included resting-state fMRI and 3 tasks probing multisensory processing^99^, cold pain^100^, and working memory^101^. The task contrasts compared combined visual-auditory stimulation with rest, cold with room-temperature arm wraps, and 2-back with 0-back blocks, respectively. Data were organized according to the Brain Imaging Data Structure and preprocessed with fMRIPrep^102^. Task effects were estimated using robust general linear models^103^, and resting-state analyses assessed regional homogeneity, amplitude of low-frequency fluctuations^104^, canonical-network connectivity, and atlas-wide connectivity.

### Task data analysis

For each task, we used linear support vector machines (SVMs) to identify task-state-related activity patterns and assessed task-state classification accuracy for out-of-sample participants using 5-fold cross-validation leaving out all images from a participant together^60–62^. The difference in pattern expression between the task and control conditions served as an individual-level measure of task-related brain activation. Group differences were assessed using 2-sided t tests, NMDAS associations using age-adjusted partial Spearman correlations, and GDF15 associations using Pearson correlations and regression models including group and age. False discovery rate correction was applied where multiple outcomes were evaluated.

Partial-correlation confidence intervals used the Fisher-z approximation; bootstrap intervals were used for mediation and robustness analyses. Bayes factors quantified evidence for non-zero versus null age-adjusted severity–brain associations^105^. Detailed acquisition, preprocessing, SVM estimation, resting-state, mediation, subgroup, robustness, and sensitivity-testing procedures are provided in the Supplementary Methods.

### Cortical mitochondrial-map analysis

Unthresholded one-sample t maps were computed from the saved participant task contrasts and averaged within the 360 HCP-MMP1 cortical parcels^106^. Atlas labels were resampled to the task-image grid using nearest-neighbor interpolation. Regional values were correlated with published MitoBrainMap measures of complexes I, II, and IV, mitochondrial density, mitochondrial respiratory capacity, and tissue respiratory capacity^27,65^. Pearson correlations were tested using 20,000 spherical rotations of the task maps, with mirrored rotations across hemispheres, surface-based parcel centroids on fsaverage10k, and nearest-neighbor reassignment^107,108^. Two-sided p values were calculated as (1 + the number of absolute null correlations at least as large as the observed absolute correlation)/(20,000 + 1). Benjamini–Hochberg FDR correction was applied across all six mitochondrial maps separately within each task.

### Between-group power analysis

Power for independent MitoD–control comparisons was calculated using two-sided, equal-variance two-sample t tests (α = 0.05) and the noncentral t distribution. Using the actual task-specific group sizes (25 patients and 49 controls for working memory, 28 and 62 for multisensory stimulation, and 29 and 62 for cold pain), we evaluated hypothetical between-group Cohen’s d values numerically equal to 50% of the reported paired task effect sizes and calculated the effect size detectable with 80% power. Paired task d_z and between-group d use different standard deviations; the fractional task effects were used as sensitivity benchmarks.

### Age-adjusted Bayes factors

Bayes factors compared non-zero versus null NMDAS–brain associations after controlling for age. We residualized ranked NMDAS and ranked brain responses against ranked age and used a Jeffreys–Zellner–Siow prior (scale r = 1)^105^; computational details are provided in the Supplementary Methods.

### Methodological details

Detailed descriptions of participant recruitment and eligibility, clinical and phenotypic assessments, MRI experimental paradigms and acquisition, imaging preprocessing, resting-state and task-fMRI analyses, and all additional statistical, machine-learning, robustness, and sensitivity analyses are provided in the Supplementary Methods.

OpenAI ChatGPT/Codex assisted with preparation and revision of scripts for figure generation, between-group power calculations, age-adjusted Bayes factors, cortical spin tests, and supplementary table verification. The analyses used the statistical procedures described above and in the Supplementary Methods.

## Acknowledgments

This work was supported by the National Institutes of Health under grants RF1AG076821, R01MH137190, R37MH076136,R21MH113011, and R01MH122706; the Seed Grant Program for MR Studies of the Zuckerman Mind Brain Behavior Institute at Columbia University; the Robert N. Butler Columbia Aging Center Fellowship Program at the Mailman School of Public Health; the National Center for Advancing Translational Sciences (UL1TR001873) and the National Cancer Institute (P30CA013696); the Columbia Irving Institute Scholars Program, the Wharton Fund, and the Baszucki Brain Research Fund (to M.P.).

## Author Contributions

MP, and TDW conceived and designed the study. CK, CT, MH, and MP acquired the clinical, phenotypic, and biomarker data. KB, MP, and TDW conceptualized the questions and analytic framework for the present study. KB performed the neuroimaging and statistical analyses with methodological input from MTdS, MP, and TDW. KB prepared the figures and wrote the first draft. All authors interpreted the data, critically revised the manuscript, and approved the final version.

## Data, Materials, and Software Availability

Requests for access to deidentified participant data and study materials should be directed to Martin Picard. Requests will be considered after publication and will be subject to participant consent, institutional review board approval, applicable institutional policies, and completion of a data-use agreement. Study-specific code will be published in the MiSBIE replication repository. General MATLAB code for analyses is available at https://github.com/canlab, with documentation and examples at https://canlab.github.io. Additional information about the MiSBIE study is available at www.picardlab.org/MiSBIE.

## Declaration of generative AI and AI-assisted technologies in the manuscript preparation process

During preparation of this manuscript, the authors used OpenAI ChatGPT/Codex to assist with manuscript organization and language editing. The authors reviewed and edited the content and take responsibility for the final manuscript.

## Supplemental information

Document S1. Supplementary Methods and Results, Figures S1–S9, Tables S1–S5, and Supplemental References.

## References

1. Attwell, D. & Laughlin, S. B. An energy budget for signaling in the grey matter of the brain. J Cereb Blood Flow Metab 21, 1133–1145 (2001).

2. Raichle, M. E. & Gusnard, D. A. Appraising the brain’s energy budget. Proc Natl Acad Sci U S A 99, 10237–10239 (2002).

3. Padamsey, Z. & Rochefort, N. L. Paying the brain’s energy bill. Curr. Opin. Neurobiol. 78, 102668 (2023).

4. Raichle, M. E. Two views of brain function. Trends Cogn. Sci. 14, 180–190 (2010).

5. Sibson, N. R. et al. Stoichiometric coupling of brain glucose metabolism and glutamatergic neuronal activity. Proc Natl Acad Sci U S A 95, 316–321 (1998).

6. Dienel, G. A. Brain glucose metabolism: Integration of energetics with function. Physiol. Rev. 99, 949–1045 (2019).

7. Mujica-Parodi, L. R. et al. Diet modulates brain network stability, a biomarker for brain aging, in young adults. Proc. Natl. Acad. Sci. U. S. A. 117, 6170–6177 (2020).

8. Imaizumi, T. et al. Identifying high-risk population of depression: association between metabolic syndrome and depression using a health checkup and claims database. Sci. Rep. 12, 18577 (2022).

9. Harper, D. G. et al. Tissue type-specific bioenergetic abnormalities in adults with major depression. Neuropsychopharmacology 42, 876–885 (2017).

10. Song, X. et al. Bioenergetics and abnormal functional connectivity in psychotic disorders. Mol. Psychiatry 26, 2483–2492 (2021).

11. Khaliulin, I., Hamoudi, W. & Amal, H. The multifaceted role of mitochondria in autism spectrum disorder. Mol. Psychiatry 30, 629–650 (2025).

12. Du, F. et al. Abnormalities in high-energy phosphate metabolism in first-episode bipolar disorder measured using 31P-magnetic resonance spectroscopy. Biol. Psychiatry 84, 797–802 (2018).

13. Procaccini, C. et al. Role of metabolism in neurodegenerative disorders. Metabolism 65, 1376–1390 (2016).

14. Anandhan, A. et al. Metabolic dysfunction in Parkinson’s disease: Bioenergetics, redox homeostasis and central carbon metabolism. Brain Res. Bull. 133, 12–30 (2017).

15. Freyberg, Z., Ford, J. M. & Phillips, M. L. Metabolism matters in mental health. Biol. Psychiatry Cogn. Neurosci. Neuroimaging 10, 239–240 (2025).

16. Monzel, A. S., Enríquez, J. A. & Picard, M. Multifaceted mitochondria: moving mitochondrial science beyond function and dysfunction. Nat. Metab. 5, 546–562 (2023).

17. Picard, M. & Shirihai, O. S. Mitochondrial signal transduction. Cell Metab. 34, 1620–1653 (2022).

18. Pekkurnaz, G. & Wang, X. Mitochondrial heterogeneity and homeostasis through the lens of a neuron. Nat. Metab. 4, 802–812 (2022).

19. Mattson, M. P., Gleichmann, M. & Cheng, A. Mitochondria in neuroplasticity and neurological disorders. Neuron 60, 748–766 (2008).

20. Cline, S. D. Mitochondrial DNA damage and its consequences for mitochondrial gene expression. Biochim. Biophys. Acta 1819, 979–991 (2012).

21. Arrázola, M. S. et al. Mitochondria in developmental and adult neurogenesis. Neurotox. Res. 36, 257–267 (2019).

22. Brunetti, D., Dykstra, W., Le, S., Zink, A. & Prigione, A. Mitochondria in neurogenesis: Implications for mitochondrial diseases. Stem Cells 39, 1289–1297 (2021).

23. Rosenberg, A. M. et al. Brain mitochondrial diversity and network organization predict anxiety-like behavior in male mice. Nat. Commun. 14, 4726 (2023).

24. Sharpley, M. S. et al. Heteroplasmy of mouse mtDNA is genetically unstable and results in altered behavior and cognition. Cell 151, 333–343 (2012).

25. Hollis, F. et al. Mitochondrial function in the brain links anxiety with social subordination. Proc. Natl. Acad. Sci. U. S. A. 112, 15486–15491 (2015).

26. Gebara, E. et al. Mitofusin-2 in the nucleus accumbens regulates anxiety and depression-like behaviors through mitochondrial and neuronal actions. Biol. Psychiatry 89, 1033–1044 (2021).

27. Mosharov, E. V. et al. A human brain map of mitochondrial respiratory capacity and diversity. Nature 641, 749–758 (2025).

28. Castrillon, G., et al. An energy costly architecture of neuromodulators for human brain evolution and cognition. bioRxiv (2023) doi:10.1101/2023.04.25.538209.

29. Chen, T., He, J., Huang, Y. & Zhao, W. The generation of mitochondrial DNA large-scale deletions in human cells. J. Hum. Genet. 56, 689–694 (2011).

30. Gorman, G. S. et al. Prevalence of nuclear and mitochondrial DNA mutations related to adult mitochondrial disease. Ann. Neurol. 77, 753–759 (2015).

31. Gorman, G. S. et al. Mitochondrial diseases. Nat. Rev. Dis. Primers 2, 16080 (2016).

32. Picard, M. & Murugan, N. J. The Energy Resistance Principle. (2025) doi:10.31219/osf.io/hgnmj_v2.

33. Set, K. K., Sen, K., Huq, A. H. M. & Agarwal, R. Mitochondrial disorders of the nervous system: A review. Clin. Pediatr. (Phila.) 58, 381–394 (2019).

34. Alston, C. L., Rocha, M. C., Lax, N. Z., Turnbull, D. M. & Taylor, R. W. The genetics and pathology of mitochondrial disease. J. Pathol. 241, 236–250 (2017).

35. Moore, H. L., Blain, A. P., Turnbull, D. M. & Gorman, G. S. Systematic review of cognitive deficits in adult mitochondrial disease. European Journal of Neurology 27, 3–17 (2020).

36. Saneto, R. P., Friedman, S. D. & Shaw, D. W. W. Neuroimaging of mitochondrial disease. Mitochondrion 8, 396–413 (2008).

37. Friedman, S. D., Shaw, D. W. W., Ishak, G., Gropman, A. L. & Saneto, R. P. The use of neuroimaging in the diagnosis of mitochondrial disease. Dev. Disabil. Res. Rev. 16, 129–135 (2010).

38. Gropman, A. L. Neuroimaging in mitochondrial disorders. Neurotherapeutics 10, 273–285 (2013).

39. Huisman, T. A. G. M., Kralik, S. F., Desai, N. K., Serrallach, B. L. & Orman, G. Neuroimaging of primary mitochondrial disorders in children: A review. J. Neuroimaging 32, 191–200 (2022).

40. Kelly, C. et al. A platform to map the mind-mitochondria connection and the hallmarks of psychobiology: the MiSBIE study. Trends Endocrinol. Metab. 35, 884–901 (2024).

41. Picard, M., Kempes, C., Pontzer, H., Behnke, A. & Shaulson, E. D. Energy constraints on human health. (2025) doi:10.31219/osf.io/ne3qg_v1.

42. Yang, X. et al. Physical bioenergetics: Energy fluxes, budgets, and constraints in cells. Proc. Natl. Acad. Sci. U. S. A. 118, e2026786118 (2021).

43. Bobba-Alves, N., Juster, R.-P. & Picard, M. The energetic cost of allostasis and allostatic load. Psychoneuroendocrinology 146, 105951 (2022).

44. Sturm, G. et al. OxPhos defects cause hypermetabolism and reduce lifespan in cells and in patients with mitochondrial diseases. *Commun*. Biol. 6, 22 (2023).

45. Sercel, A. J. et al. Hypermetabolism and energetic constraints in mitochondrial disorders. Nat. Metab. 6, 192–195 (2024).

46. Jamadar, S. D., Behler, A., Deery, H. & Breakspear, M. The metabolic costs of cognition. Trends Cogn. Sci. 29, 541–555 (2025).

47. Miyake, A. et al. The unity and diversity of executive functions and their contributions to complex ‘Frontal Lobe’ tasks: a latent variable analysis. Cogn. Psychol. 41, 49–100 (2000).

48. Conway, A. R. A., Kane, M. J. & Engle, R. W. Working memory capacity and its relation to general intelligence. Trends Cogn. Sci. 7, 547–552 (2003).

49. Schaefer, A. M. et al. Mitochondrial disease in adults: a scale to monitor progression and treatment. Neurology 66, 1932–1934 (2006).

50. Hara, Y., Yuk, F., Puri, R., Janssen, W. G. M., Rapp, P. R. & Morrison, J. H. Presynaptic mitochondrial morphology in monkey prefrontal cortex correlates with working memory and is improved with estrogen treatment. Proc Natl Acad Sci 111, 486–491 (2014).

51. Kelley, D. P. et al. The allostatic triage model of psychopathology (ATP Model): How reallocation of brain energetic resources under stress elicits psychiatric symptoms. Neurosci. Biobehav. Rev. 179, 106419 (2025).

52. Bruckmaier, M., Tachtsidis, I., Phan, P. & Lavie, N. Attention and capacity limits in perception: A cellular metabolism account. J. Neurosci. 40, 6801–6811 (2020).

53. Flippo, K. H. & Potthoff, M. J. Metabolic messengers: FGF21. Nat. Metab. 3, 309–317 (2021).

54. Patel, S. et al. GDF15 provides an endocrine signal of nutritional stress in mice and humans. Cell Metab. 29, 707–718.e8 (2019).

55. Chang, J. Y. et al. The role of growth differentiation factor 15 in energy metabolism. Diabetes Metab. J. 44, 363–371 (2020).

56. Sharma, R. et al. Circulating markers of NADH-reductive stress correlate with mitochondrial disease severity. J. Clin. Invest. 131, e136055 (2021).

57. Wager, T. D., Davidson, M. L., Hughes, B. L., Lindquist, M. A. & Ochsner, K. N. Prefrontal-subcortical pathways mediating successful emotion regulation. Neuron 59, 1037–1050 (2008).

58. Yarkoni, T., Poldrack, R. A., Nichols, T. E., Van Essen, D. C. & Wager, T. D. Large-scale automated synthesis of human functional neuroimaging data. Nat Methods 8, 665–670 (2011).

59. Bo, K. et al. A systems identification approach using Bayes factors to deconstruct the brain bases of emotion regulation. Nat. Neurosci. 27, 975–987 (2024).

60. Kohoutová, L. et al. Individual variability in brain representations of pain. Nat. Neurosci. 25, 749–759 (2022).

61. Kragel, P. A., Koban, L., Barrett, L. F. & Wager, T. D. Representation, pattern information, and brain signatures: From neurons to neuroimaging. Neuron 99, 257–273 (2018).

62. Baranger, D. A. A. et al. Enhancing task fMRI individual difference research with neural signatures. medRxiv (2025) doi:10.1101/2025.01.30.25321355.

63. Haast, R. A. M. et al. Anatomic & metabolic brain markers of the m.3243A>G mutation: A multi-parametric 7T MRI study. NeuroImage Clin. 18, 231–244 (2018).

64. Evangelisti, S. et al. Molecular biomarkers correlate with brain grey and white matter changes in patients with mitochondrial m.3243A > G mutation. Mol. Genet. Metab. 135, 72–81 (2022).

65. Guo, F. et al. Assessing molecular, cellular and transcriptomic bases of laminar perfusion and cytoarchitecture coupling in the human cortex. Nat. Commun. 17, 9748 (2026). doi:10.1038/s41467-026-76812-w.

66. Fujita, Y. et al. GDF15 is a novel biomarker to evaluate efficacy of pyruvate therapy for mitochondrial diseases. Mitochondrion 20, 34–42 (2015).

67. Huang, Q., et al. The energetic stress marker GDF15 is induced by acute psychosocial stress. bioRxiv (2024) doi:10.1101/2024.04.19.590241.

68. Deng, Y.-T. et al. Atlas of the plasma proteome in health and disease in 53,026 adults. Cell 188, 253–271.e7 (2025).

69. Huang, Q. et al. The mitochondrial disease biomarker GDF15 is dynamic, quantifiable in saliva, and correlates with disease severity. Mol. Genet. Metab. 145, 109179 (2025).

70. Lockhart, S. M., Saudek, V. & O’Rahilly, S. GDF15: A hormone conveying somatic distress to the brain. Endocr. Rev. 41, 610–642 (2020).

71. Engström Ruud, L., et al. Activation of GFRAL+ neurons induces hypothermia and glucoregulatory responses associated with nausea and torpor. Cell Rep. 43, 113960 (2024).

72. Shaulson, E. D., Cohen, A. A. & Picard, M. The brain-body energy conservation model of aging. *Nat*. Aging 4, 1354–1371 (2024).

73. Tanaka, T. et al. Plasma proteomic biomarker signature of age predicts health and life span. Elife 9, e61073 (2020).

74. Lehallier, B. et al. Undulating changes in human plasma proteome profiles across the lifespan. Nat. Med. 25, 1843–1850 (2019).

75. Trumpff, C. et al. Blood mitochondrial health markers cf-mtDNA and GDF15 in human aging. bioRxiv (2025). doi:10.1101/2025.01.28.635306.

76. Ripp, I. et al. Working memory task induced neural activation: A simultaneous PET/fMRI study. Neuroimage 237, 118131 (2021).

77. DiNuzzo, M. et al. Perception is associated with the brain’s metabolic response to sensory stimulation. eLife 11, e71016 (2022).

78. Wan, B., Riedl, V., Castrillon, G., Kirschner, M. & Valk, S. L. Bridging Glucose Metabolism and Intrinsic Functional Organization of the Human Cortex. *Commun*. Biol. 9, 377 (2026). doi:10.1038/s42003-026-09693-w.

79. Raichle, M. E. The brain’s dark energy. Science 314, 1249–1250 (2006).

80. Hyder, F., Fulbright, R. K., Shulman, R. G. & Rothman, D. L. Glutamatergic function in the resting awake human brain is supported by uniformly high oxidative energy. J. Cereb. Blood Flow Metab. 33, 339–347 (2013).

81. Cheng, A., Hou, Y. & Mattson, M. P. Mitochondria and neuroplasticity. ASN Neuro 2, e00045 (2010).

82. Jeanneteau, F. & Arango-Lievano, M. Linking mitochondria to synapses: New insights for stress-related neuropsychiatric disorders. Neural Plast. 2016, 3985063 (2016).

83. Rocca, M. A., Schoonheim, M. M., Valsasina, P., Geurts, J. J. G. & Filippi, M. Task- and resting-state fMRI studies in multiple sclerosis: From regions to systems and time-varying analysis. Current status and future perspective. NeuroImage: Clinical 35, 103076 (2022).

84. Turner, M. P. et al. Altered linear coupling between stimulus-evoked blood flow and oxygen metabolism in the aging human brain. Cereb. Cortex 33, 135–151 (2023).

85. Iadecola, C. The neurovascular unit coming of age: A journey through neurovascular coupling in health and disease. Neuron 96, 17–42 (2017).

86. Mastrobattista, E. et al. Late-life depression is associated with increased levels of GDF-15, a pro-aging mitokine. Am. J. Geriatr. Psychiatry 31, 1–9 (2023).

87. Azarsız, Y. D. et al. Serum FGF21 and GDF15 levels and their association with cognitive function in bipolar disorder. J. Affect. Disord. 397, 120979 (2026).

88. Jones-Murphy, A. et al. Circulating Growth Differentiation Factor-15 (GDF-15) levels in depressive disorders - A systematic review and meta-analysis of the association between GDF-15 and depression in adult populations. J. Affect. Disord. 408, 121837 (2026).

89. Beydoun, M. A. et al. GDF15 and its association with cognitive performance over time in a longitudinal study of middle-aged urban adults. Brain Behav. Immun. 108, 340–349 (2023).

90. Xue, X.-H., Tao, L.-L., Su, D.-Q., Guo, C.-J. & Liu, H. Diagnostic utility of GDF15 in neurodegenerative diseases: A systematic review and meta-analysis. Brain Behav. 12, e2502 (2022).

91. Behnke, A., Shaulson, E., Pontzer, H., Kempes, C. P. & Picard, M. Energy constraint on human health. Trends Endocrinol. Metab. (2026) doi:10.1016/j.tem.2026.02.010.

92. Bernardo, T. C. et al. Physical exercise and brain mitochondrial fitness: The possible role against Alzheimer’s disease. Brain Pathol. 26, 648–663 (2016).

93. Marseglia, A. et al. Metabolic syndrome is associated with poor cognition: A population-based study of 70-year-old adults without dementia. J. Gerontol. A Biol. Sci. Med. Sci. 76, 2275–2283 (2021).

94. Féart, C., Samieri, C. & Barberger-Gateau, P. Mediterranean diet and cognitive function in older adults. Curr. Opin. Clin. Nutr. Metab. Care 13, 14–18 (2010).

95. Maeda, K. et al. Clinical phenotype and segregation of mitochondrial 3243A>G mutation in 2 pairs of monozygotic twins. JAMA Neurol. 73, 990–993 (2016).

96. Picard, M. & Hirano, M. Disentangling (epi)genetic and environmental contributions to the mitochondrial 3243A>G mutation phenotype: Phenotypic Destiny in mitochondrial disease? JAMA Neurol. 73, 923–925 (2016).

97. Jain, I. H. et al. Hypoxia as a therapy for mitochondrial disease. Science 352, 54–61 (2016).

98. Kelly, C. et al. Perceived association of mood and symptom severity in adults with mitochondrial diseases. Mitochondrion 84, 102033 (2025).

99. López-Solà, M. et al. Towards a neurophysiological signature for fibromyalgia. Pain 158, 34–47 (2017).

100. Porcelli, A. J. An alternative to the traditional cold pressor test: The cold pressor arm wrap. J. Vis. Exp. 83, e50849 (2014). doi:10.3791/50849.

101. Barch, D. M. et al. Function in the human connectome: task-fMRI and individual differences in behavior. Neuroimage 80, 169–189 (2013).

102. Esteban, O. et al. fMRIPrep: a robust preprocessing pipeline for functional MRI. Nat. Methods 16, 111–116 (2019).

103. Wager, T. D., Keller, M. C., Lacey, S. C. & Jonides, J. Increased sensitivity in neuroimaging analyses using robust regression. Neuroimage 26, 99–113 (2005).

104. Yan, C. G., Wang, X. D., Zuo, X. N. & Zang, Y. F. DPABI: Data Processing & Analysis for (Resting-State) Brain Imaging. Neuroinformatics 14, 339–351 (2016).

105. Wetzels, R. & Wagenmakers, E.-J. A default Bayesian hypothesis test for correlations and partial correlations. Psychon. Bull. Rev. 19, 1057–1064 (2012). 10.3758/s13423-012-0295-x

106. Glasser, M. F. et al. A multi-modal parcellation of human cerebral cortex. Nature 536, 171–178 (2016).

107. Alexander-Bloch, A. F. et al. On testing for spatial correspondence between maps of human brain structure and function. Neuroimage 178, 540–551 (2018).

108. Markello, R. D. et al. neuromaps: structural and functional interpretation of brain maps. Nat. Methods 19, 1472–1479 (2022).

