## Supplementary material for "Mitochondrial Energy Transformation Capacity Influences Brain Activation During Sensory, Affective, and Cognitive Tasks": SI

Supplementary Methods and Results; Figures S1–S9; Tables S1–S5; Supplemental References.

### Supplementary Methods

#### Participants

The Mitochondrial Stress, Brain Imaging, and Epigenetics (MiSBIE) study was conducted in Columbia University Irving Medical Center in adherence to the directives outlined by the New York State Psychiatric Institute IRB protocol #7424, ClinicalTrials.gov NCT04831424. From 2018 to 2023, English-speaking adults aged 18–60 were enrolled through clinical and research networks at Columbia University (e.g., the Neuromuscular Clinic, RecruitMe), the Children's Hospital of Philadelphia, and national/international organizations focused on mitochondrial disease (e.g., NAMDC, UMDF). Participants were excluded for recent illness, pregnancy, current cancer, use of steroids, participation in other clinical trials, or cognitive impairment (TICS-41 ≤30). Participants visited Columbia University Irving Medical Center for two consecutive days of data collection, including biospecimen collection, medical exam, stress reactivity–recovery session, detailed physiological measurements, standing and functional capacity test, metabolic rate analysis, psychosocial questionnaires, and neuropsychological assessment. On the second day, participants visited The Zuckerman Institute at Columbia University for brain imaging, including T1, T2, field map, resting state, Diffusion-Weighted Imaging (DWI), and task fMRI [1].

In total, 110 participants are included in the Mitochondrial Stress, Brain Imaging, and Epigenetics (MISBIE) study. 40 MitoD patients, including 20 patients with m.3243A>G mutation, 15 patients with single deletion in mDNA, and 5 patients with Mitochondrial Encephalomyopathy (MELAS), with 70 healthy controls. The age and gender are matched between the two groups (Figure S9). High-resolution structural and functional MRI data were available for 91 participants (29 MitoD: 16 point mutations, 11 deletions, 2 MELAS; and 62 controls). Of these 110 participants, 19 did not complete the Day 2 imaging session for logistical or scheduling reasons unrelated to neurological function, such as travel constraints or time limitations; therefore, MRI data were never acquired for these individuals. High-resolution structural and functional MRI data were available for the remaining 91 participants, including 29 MitoD patients (16 point mutations, 11 deletions, 2 MELAS) and 62 controls.

Task-specific data availability differed slightly across paradigms. Three participants did not complete the N-back working-memory task, and one participant did not complete the multisensory task. Thus, N-back imaging data were available for 88 participants, cold-pain imaging data for 91 participants, and multisensory imaging data for 90 participants. Fourteen participants were further excluded from the primary N-back working-memory analyses based on a prespecified behavioral data-quality/task-engagement criterion: unavailable or severely incomplete behavioral response data (n = 12) or usable response logs with accuracy below 40% (n = 2). Of these 14 exclusions, 11 were healthy controls and 3 were MitoD patients. This criterion was used to exclude uninterpretable task engagement, not low-performing participants. In this binary forced-choice task, response-recording or button-mapping problems were considered potential explanations for the two usable response logs with accuracy below 40%. The criterion was applied identically across MitoD and control participants and had no systematic relationship to disease severity. Sensitivity analyses retaining all participants with available N-back imaging are reported in Figure S3. The overall pattern of results was similar, although the severe-patient versus control comparison of N-back brain pattern expression was no longer significant.

The final sample sizes differed by task: for the N-back working memory task, 88 participants (28 MitoD: 15 point mutations, 11 deletions, 2 MELAS) provided usable data, of whom 74 (25 MitoD: 14 point mutations, 9 deletions, 2 MELAS) remained after applying the behavioral data-quality/task-engagement criterion; for the cold-pain task, 91 participants (29 MitoD: 16 point mutations, 11 deletions, 2 MELAS) had complete data; and for the multisensory task, 90 participants (28 MitoD: 16 point mutations, 10 deletions, 2 MELAS) were included.

#### Materials and procedures

In addition to neuroimaging, all participants completed a comprehensive phenotypic battery covering eleven domains. Self-report questionnaires assessed key psychological factors (anxiety, depression, burnout, stress, personality traits, PTSD symptoms, tic severity), social and environmental exposures (social support, childhood trauma, life events), and health behaviors (physical activity, sleep quality). Clinically relevant symptom scales captured fatigability (physical, cognitive, psychosocial), autonomic dysfunction, and headache severity, while objective tests measured sit-to-stand performance, body composition, pulmonary function ($FEV_1$, FVC), and neurological status (NMDAS, CNS score, Karnofsky). Cognitive function was probed with standardized neuropsychological tasks of executive control, memory, and fluid intelligence (e.g., verbal fluency, trail-making, Stroop, RBANS, WASI). Resting energy metabolism and respiratory parameters were quantified via indirect calorimetry, and stress reactivity was indexed by heart rate, respiration, and salivary cortisol. Finally, blood and saliva assays provided markers of mitochondrial energetics (mtDNA copy number), metabolic health (glucose, lipids, electrolytes, $HbA_1c$), mitochondrial stress (GDF15, FGF21, lactate), and immune/inflammatory status (CBC, CRP, fibrinogen). (See [1] for more details)

A comprehensive assessment,including the Newcastle Mitochondrial Disease Adult Scale (NMDAS), Columbia Neurological Score (CNS), Karnofsky Performance Scale (KPS), Composite Autonomic Symptom Score (COMPASS), and Modified Fatigue Impact Scale (MFIS), was administered to capture the multisystem impact of mitochondrial disease. The NMDAS provides a standardized index of disease-specific symptom severity across neuromuscular, exercise intolerance, and metabolic domains. The CNS aggregates clinician-rated neurological findings (e.g., cranial nerve function, coordination, gait, and cognition) into a single severity score. The KPS rates overall functional capacity on a 0–100 scale, reflecting a patient’s ability to perform daily activities and self-care. The COMPASS quantifies autonomic dysfunction across orthostatic, vasomotor, sudomotor, gastrointestinal, bladder, and pupillomotor systems, yielding both domain-specific and total autonomic burden scores. Finally, the MFIS assesses the perceived impact of fatigability on physical, cognitive, and psychosocial functioning. Together, these instruments provide a comprehensive clinical profile—ranging from symptom burden and neurological impairment to daily-life functioning and autonomic stability—that we used both to characterize group differences and to relate systemic disease severity to neuroimaging and metabolic biomarkers.

We selected NMDAS as the primary patient-level severity index because it is the most widely validated, clinician-administered, and disease-specific composite scale in mitochondrial medicine, and is intentionally systems-agnostic — capturing overall disease burden across all organ systems rather than overweighting any particular symptom domain[2]. Specifically, it includes a standardized index of disease-specific symptom severity across neuromuscular, exercise intolerance, and metabolic domains. This property is particularly important for neuroimaging analyses, where we aim to capture overall bioenergetic stress rather than a specific clinical manifestation.

##### MRI Experimental paradigm

The MRI session for the MISBIE study took place on the second day of the participants' visit. The imaging acquisition followed a sequence that included a localizer (1 minute), a field map (1 minute), a T2 structural scan (11 minutes and 15 seconds), a T1 structural scan (5 minutes and 21 seconds), a multisensory task (5 minutes), a resting state scan (10 minutes and 51 seconds), an N-back task (2 runs, each lasting 4 minutes and 28 seconds), a social-evaluative task (6 minutes), and a cold pain task (6 minutes), ending with a DWI scan (Sequence 1-2, 6 minutes and 27 seconds). In the current analysis, we focused on the resting state and task data from the multisensory, N-back, and cold pain tasks. The speech preparation task was discarded due to misalignment issues between the experiment output and the MRI data.

##### Multisensory Task (Basic Sensory)

In the multisensory task, participants were exposed to both visual stimulation (a flickering checkerboard pattern) and auditory stimulation (a sequence of sounds with gradually increasing and decreasing pitch) presented in a 30-second block (See settings also in [3]). Each task block was preceded by a 30-second resting block during which participants were instructed to fixate on a central cross. The entire session lasted 5 minutes, with the rest-task cycle repeated five times. The contrast between the task and rest periods was used to model the brain's response to multisensory processing in the fMRI analysis.

##### N-back Task (Working Memory)

The N-back task followed a block design as described previously [4]. Within each task block, a word was presented for one second to indicate the block type (0-back or 2-back). Each block consisted of 10 trials featuring 10 different stimuli of the same type, and each block used a different set of stimuli. Participants were instructed to respond with a button press, indicating either "yes, this image matches the image presented n trials ago" (target response) or "no, this image does not match the image presented n trials ago" (nontarget response). Responses were recorded to calculate behavioral accuracy. Both the 2-back and 0-back blocks lasted 25 seconds, followed by a 15-second rest period. The contrast between the 2-back (high cognitive load) and 0-back (low cognitive load) conditions was used to model the brain's response during working memory tasks.

Cold Pain Task (Negative Affect)

In the cold pain task, the study coordinator equipped the participant with the MR-compatible Cold Pressor Arm Wrap (CPAW). No pain stimulation was administered prior to entering the scanner beyond explaining the procedure to participants.

During the fMRI scan, participants completed a single cold-pain block consisting of four consecutive phases applied to the right hand and wrist. At the beginning of the scan, the coordinator first placed a room-temperature wrap on the participant's right hand and wrist for 90 seconds. This was followed by a 30-second buffer period before applying the cold pack, which then covered the right hand and wrist for an additional two minutes. After a second 30-second buffer period, the session concluded with a 90-second recovery period using the room-temperature wrap. Thus, the cold pain task consisted of one block with a total duration of 6 minutes.

Inside the cold wrapper, Solution is MRI-safe gelpacs. The coldpack were stored in a small freezer set to approximately 0 °F (though temperatures could vary slightly) and were removed about 15 minutes before the task to reach a safe, near-freezing temperature. The procedure conducted in the experiment is as follows [5] . In the fMRI analysis, the contrast between the cold pressor and the initial room-temperature period was used to model the brain's response to cold pain.

To assess pain elicited by the CPAW, participants rated their discomfort on a 0–10 scale at the end of the experiment (0 = no pain; 10 = intolerable).

Imaging acquisition parameters

In the MRI session, we measured brain anatomy using T1- and T2-weighted magnetic resonance imaging on a 3 T Siemens Prisma scanner (Siemens Medical Solutions). The sequence parameters for T1 were: echo time, 0.00349 s; repetition time, 2.3 s; flip angle, 8°; voxel size, 1 × 1 × 1 mm; slice thickness, 1 mm. Parallel imaging with a reduction factor of 2 was used. The parameters for T2 were: echo time, 0.562 s; repetition time, 3.2 s; flip angle, 120°; voxel size, 0.7 × 0.7 × 0.7 mm; slice thickness, 0.7 mm. Parallel imaging with a reduction factor of 2 was used. We collected functional MRI images on the same scanner to measure BOLD activity during a series of psychological tasks. The sequence parameters were as follows: echo time, 0.029 s; repetition time, 0.46 s; flip angle, 44°; voxel size, 3.02 × 3.02 × 3 mm; slice thickness, 3 mm. Slices were acquired with a multiband acceleration factor of 8.

Data preprocessing

Data is transformed into the standard Brain Imaging Data Structure (BIDS) format and preprocessed by fMRIprep [6].

Anatomical images were processed with fMRIPrep (v20.2.1). T1-weighted volumes were bias-field corrected and skull-stripped. Tissue segmentation into gray matter, white matter, and cerebrospinal fluid was performed, and cortical surfaces were reconstructed. The resulting brain mask and surfaces were used to guide nonlinear normalization of the anatomical volume to the MNI152 template space.

Functional preprocessing was also carried out with fMRIPrep (v20.2.1). For each BOLD run, a reference volume was generated, and head-motion parameters were estimated. Slice-timing correction and motion realignment were applied, then functional data were co-registered to the T1-weighted image. Field-map–based distortion correction was disabled due to mismatched shim parameters between field maps and functional scans. Physiological and motion confounds—including framewise displacement, DVARS, and CompCor components—were computed, and motion outlier frames were flagged. Because outlier volumes were modeled as nuisance regressors rather than removed, there is no fixed scrubbing threshold or frame-removal percentage. Quantitative motion metrics (mean FD and DVARS) per task and group are reported in Fig. S2. Finally, preprocessed time series were normalized to MNI152 space and projected onto the fsaverage surface for surface-based analyses.

Resting state analysis

After preprocessing by fMRI prep, resting state data is further processed by regressing out nuisance covariate including 24 headmotion parameters extracted from fMRIprep and CSF signal and its derivatives. An additional covariate measurement includes spike indicator regressors that detected outliers based on Mahalanobis distance, which measures how different each image is from the rest of the images. An image is detected as a spike if its corresponding Mahalanobis distance is outside the 95% confidence region of the cloud of images in multidimensional space. Multiple comparison is controlled by $p<0.05$ (Bonferroni). In the regressor, each individual potential outlier was coded as 1, and other volumes were coded as 0.

Additionally, the data is detrended and smoothed using a 4mm Gaussian smooth kernel for further analysis.

To calculate the regional connectivity and spontaneous activity across brain, a Regional Homogeneity (Reho) analysis and an Amplitude of Low Frequency Fluctuations (ALFF) are done using Data Processing & Analysis of Brain Imaging (DIPABI) software [7]. Band-pass filter is applied for ALFF in 0.01-0.1Hz and 0.01-0.08Hz for Reho.

Task data analysis

Within-participant task effects were estimated using a general linear model (GLM). Robust regression was applied in the GLM to reduce the influence of outliers and improve sensitivity to the task signal [8]. Task and control blocks were modeled as regressors using boxcar functions convolved with the canonical hemodynamic response function (HRF) via CanlabTools (https://github.com/canlab).

The timing and duration of the regressors of interest were based on the onset and duration of each stimulus. A high-pass filter or detrending was applied to remove low-frequency temporal drift, depending on the task design. Specifically, both high-pass filtering and detrending were applied in the multisensory and N-back tasks. In contrast, only detrending was applied in the cold pain task because the relatively long task blocks could lead to the loss of task-related signals if a high-pass filter were used. Nuisance regressors included 24 head motion parameters (six rigid-body motion parameters, their squares, and their temporal derivatives), spike indicator regressors for potential outliers, and cerebrospinal fluid (CSF) time series along with their derivatives. CSF time series were extracted using CANlab tools from canonical CSF masks. The six head motion covariates were derived from the realignment parameters generated by fMRIPrep. Spike regressors were defined as described in the resting-state analysis section.

For each condition (task and control), whole-brain beta coefficient maps were generated separately at the participant level. At the group level, participants' beta maps were compared between task and control conditions. The contrasts of interest included ‘Cold’ vs. ‘Room Temperature’ in the cold pain task, ‘2-back’ vs. ‘0-back’ in the N-back task, and ‘Multisensory’ vs. ‘Rest’ in the multisensory task. In the second-level analysis, CSF signals were regressed out for each participant to control for potential confounds. The processed data were then used for final analyses.

Covariates and sensitivity analyses

To assess the robustness of primary findings to potential confounds, we conducted sensitivity analyses including sex, genotype/variant class (point mutation, single deletion, MELAS), fatigue subscales, and vision-related neurological measures as covariates in key regression models. Disease duration was not modeled as a covariate because mtDNA pathogenic variants are present from birth, and clinical onset varies widely across individuals, rendering a single duration metric ill-defined. Results of these analyses are reported in Tables S4 and S5.

Clustering analysis

For the clustering analysis, we computed pairwise partial Spearman correlations among all variables, statistically controlling for group membership (MitoD vs. control) to ensure that the resulting correlation structure reflected co-variation among variables independent of group differences. We then used a permutation test (Wager et al., 2008) to determine the optimal number of clusters, in which the observed clustering solution was compared against 1,000 random permutations of the data to identify the number of clusters that explained significantly more variance than chance. This analysis indicated that two clusters was the statistically optimal solution ($p < 0.05$). However, to capture finer-grained phenotypic structure, we selected $k = 11$, which corresponded to a local maximum in the permutation-based pseudo-Z statistic, balancing statistical rigor with biological interpretability. We then applied hierarchical clustering using Ward's linkage method on the distance matrix derived from the partial Spearman correlation matrix, which groups variables by minimizing within-cluster variance at each step.

Correlation robustness

To address the concern that all correlations are within-sample, we conducted a split-half robustness analysis for the primary brain–severity relationships. We repeatedly split the patient group ($n = 25$) into two random halves across 1,000 iterations and recomputed the age-controlled Spearman partial correlation between working-memory brain activation and NMDAS — and between NMDAS and behavioral performance — in each half independently. The mean of the two half-sample correlations was recorded per iteration, yielding an empirical distribution of split-half estimates.

Results were highly consistent across all splits. For NMDAS versus working-memory brain activation, the full-sample partial $r = -0.670$, with a median split-half $r = -0.646$ (95% CI [−0.762,

−0.474]), and 100% of iterations yielded a negative correlation consistent with the full-sample direction. For NMDAS versus behavioral performance, the full-sample age-adjusted r = −0.460, with a median split-half r = −0.453 (95% CI [−0.589, −0.277]), again with 100% directional consistency across all iterations.

The near-identical median split-half estimates relative to the full-sample values, combined with 100% directional consistency across 1,000 iterations, indicate that these associations are stable and not driven by any particular subset of patients. We note that the width of the confidence intervals reflects the expected increase in sampling variability when halving an already small patient sample (each half n ≈ 12), rather than any instability in the underlying relationship.

Machine learning classification

We used a multivariate pattern classification approach based on support vector machines (SVMs) to decode whole-brain neural representations distinguishing task and control states (e.g., 2-back vs. 0-back in the working memory task). Analyses were conducted in voxel space using whole-brain participant-level activation maps.

For each participant, first-level general linear models were estimated, and condition-specific beta maps were extracted for the task and control conditions. These beta maps served as feature vectors for classification, with each voxel treated as one feature. Condition labels (task vs. control) were assigned accordingly.

Classification was performed using a linear SVM, which is well suited for high-dimensional neuroimaging data and yields interpretable weight maps reflecting discriminative voxel patterns. To assess generalizability across individuals, we employed 5-fold cross-validation with participants as the unit of partitioning (leave-whole-subject-out). In each fold, beta maps from approximately 80% of participants were used to train the classifier, and beta maps from the held-out participants were used for testing. Classification performance was aggregated across folds.

A whole-brain SVM weight map was computed by averaging the trained classifier weights across cross-validation folds. To assess the stability and statistical reliability of voxel-wise weights, we conducted 5,000 bootstrap resampling iterations across participants. For each bootstrap sample, the SVM was retrained and voxel-wise weights were recomputed, yielding a bootstrap distribution for each voxel. These distributions were used to estimate voxel-wise significance of the SVM weights.

To quantify the degree to which each participant expressed the task-related neural pattern, we computed pattern expression values by taking the dot product between the SVM weight map estimated without that participant and each participant's task and control beta maps. The difference in pattern expression between task and control conditions was used as an index of task-related neural engagement for each participant, hereafter referred to as task brain expression. The paired plots in Figure 2A show condition scores from the final cross-validation repetition; Results report the mean task effect sizes across repetitions. For display only, cold-pain pairs were omitted if either condition exceeded three scaled median absolute deviations from its median. All pairs remained in effect-size calculations and statistical analyses.

Bayes factor analysis

Bayes factors (BFs) were used to evaluate the relative evidence for non-zero versus null age-adjusted NMDAS–brain associations. $BF_{10}$ is the ratio of the marginal likelihood under the alternative to that under the null hypothesis. Within each task, ranked NMDAS and ranked brain pattern expression were residualized against an intercept and ranked age. We used a Jeffreys–Zellner–Siow mixture of g-priors, $g \sim \text{inverse-gamma}(1/2, nr^2/2)$, with scale $r = 1$ and matching nuisance priors under both hypotheses[9]. For residual vectors x and y, $R^2 = (x'y)^2/[(x'x)(y'y)]$ and $v = n - 2$; the fixed-g marginal-likelihood ratio is $(1 + g)$^$((v - 1)/2)$ / $[1 + g(1 - R^2)]$^$(v/2)$. Numerical integration over g yielded $BF_{10}$ = 89.54, 0.88, and 0.23 for working memory, multisensory processing, and cold pain, respectively. Prior scales $r = 0.5$ and 2 yielded 86.50 and 64.11 for working memory, 1.33 and 0.49 for multisensory processing, and 0.40 and 0.12 for cold pain.

Preregistration statement

The primary hypotheses for the MiSBIE study were preregistered at https://clinicaltrials.gov/study/NCT04831424. However, the neuroimaging analyses were not. Group differences and brain–clinical associations were motivated by prior literature and theoretical considerations. Patient subgroups were identified using data-driven K-means clustering; the broader phenome-wide association analyses were exploratory.

#### Supplementary Results

##### Additional sensitivity analyses

Adjusting the key NMDAS–brain and NMDAS–behavior associations for sex and genotype/variant class did not materially alter the effect sizes or statistical significance. Sensitivity analyses involving fatigue, vision-related neurological measures, and psychiatric symptoms are reported in Tables S4 and S5.

In the subset of patients with available structural imaging data ($n = 23$), the association between working-memory brain activation and NMDAS remained significant after simultaneously controlling for cortical thickness and fractional anisotropy (partial $r = -0.536$, $p = 0.012$). Controlling for the NODDI microstructural measures ICVF and Max likewise left the association significant (partial $r = -0.662$, $p = 0.001$).

##### Resting state brain activity

Beyond task-related activity, spontaneous brain activity, including within large-scale resting-state networks like the default mode network and the frontoparietal network, accounts for a substantial portion of the brain's energy consumption [10–12]. To assess how mitochondrial MitoD-related energy transformation defects influence this activity, we examined spontaneous neural fluctuations using Amplitude of Low Frequency Fluctuation (ALFF) and Regional Homogeneity (ReHo) [13,14].

No significant group differences were observed between MitoD patients and healthy controls at FDR-corrected $q < 0.05$. However, at a more liberal threshold ($p < 0.001$ uncorrected, cluster size > 20 voxels), the MitoD group showed lower ALFF in the cerebellum and higher ReHo in the dorsomedial and ventromedial prefrontal cortices (dmPFC and vmPFC) (Figure S5A). We further tested whether disease severity, measured by the Newcastle Mitochondrial Disease Adult Scale (NMDAS), correlated with ALFF or ReHo within the MitoD group. No associations were significant at $p < 0.001$ level. Similarly, when we compared average ALFF and ReHo across seven canonical resting-state networks [15], we found no significant differences between MitoD and Controls or correlations with NMDAS, providing no evidence of widespread spontaneous activity abnormalities in MitoD.

Next, we compared resting-state functional connectivity differences between MitoD patients and controls among the same seven networks (Figure S5B), controlling for participant-wise head motion (average framewise displacement). Robust regression revealed selective increases in connectivity in the MitoD group at uncorrected thresholds ($p < 0.05$) for connections between (a) frontoparietal and somatosensory networks, and (b) frontoparietal and ventral attention networks. However, these results did not survive correction for multiple comparisons at FDR $q<0.05$. We also computed connectivity across 518 gray-matter parcels defined based on widely used atlases of the cortex, basal ganglia, thalamus, and brainstem (Figure S6), but again found no FDR-corrected differences; qualitatively, there was a trend toward increased cortical-to-cortical connectivity in MitoD patients, which decreased further with greater disease severity.

In summary, these results from rare individuals with genetic mitochondrial defects do not appear to cause strong or widespread alterations in spontaneous brain activity or network-level connectivity at rest. These measures also did not correlate with disease severity, consistent with largely preserved resting-state brain activity and connectivity.

For working-memory brain pattern response, several associations survived FDR correction:

- Positive correlations with cognitive performance (Letter Sequencing), the Columbia Neurologic Score (CNS), and DSM-5 Disinhibition;
- Negative correlations with NMDAS and GDF15.

At a more lenient threshold ($p < 0.05$ uncorrected), additional associations involved cognitive tests, fatigue, autonomic measures, and metabolic biomarkers (Table S2).
For behavioral performance, no associations survived FDR correction. At an uncorrected threshold, performance showed:

- Negative correlations with COMPASS secretomotor scores, GDF15, NMDAS, and several inflammatory/metabolic markers;
- Positive correlations with the CNS and cognitive task measures (Letter Sequencing, Number–Letter Sequencing).

Across these analyses, NMDAS and GDF15 were consistently associated with working-memory brain activation. The broader phenome-wide results are reported in Tables S2 and S3.

### SI References


1. Kelly, C. et al. A platform to map the mind-mitochondria connection and the hallmarks of psychobiology: the MiSBIE study. Trends Endocrinol. Metab. 35, 884–901 (2024).
2. Schaefer, A. M. et al. Mitochondrial disease in adults: a scale to monitor progression and treatment. Neurology 66, 1932–1934 (2006).
3. López-Solà, M. et al. Towards a neurophysiological signature for fibromyalgia. Pain 158, 34–47 (2017).
4. Barch, D. M. et al. Function in the human connectome: task-fMRI and individual differences in behavior. Neuroimage 80, 169–189 (2013).
5. Porcelli, A. J. An alternative to the traditional cold pressor test: The cold pressor arm wrap. J. Vis. Exp. 83, e50849 (2014). doi:10.3791/50849.
6. Esteban, O. et al. fMRIPrep: a robust preprocessing pipeline for functional MRI. Nat. Methods 16, 111–116 (2019).
7. Yan, C. G., Wang, X. D., Zuo, X. N. & Zang, Y. F. DPABI: Data Processing & Analysis for (Resting-State) Brain Imaging. Neuroinformatics 14, 339–351 (2016).
8. Wager, T. D., Keller, M. C., Lacey, S. C. & Jonides, J. Increased sensitivity in neuroimaging analyses using robust regression. Neuroimage 26, 99–113 (2005).
9. Wetzels, R. & Wagenmakers, E.-J. A default Bayesian hypothesis test for correlations and partial correlations. Psychon. Bull. Rev. 19, 1057–1064 (2012).
10. Raichle, M. E. & Gusnard, D. A. Appraising the brain's energy budget. Proc. Natl. Acad. Sci. U.S.A. 99, 10237–10239 (2002).
11. Tomasi, D., Wang, G.-J. & Volkow, N. D. Energetic cost of brain functional connectivity. Proc. Natl. Acad. Sci. U.S.A. 110, 13642–13647 (2013).
12. Godbersen, G. M. et al. Task-evoked metabolic demands of the posteromedial default mode network are shaped by dorsal attention and frontoparietal control networks. eLife 12, e84683 (2023).
13. Zou, Q.-H. et al. An improved approach to detection of amplitude of low-frequency fluctuation (ALFF) for resting-state fMRI: fractional ALFF. J. Neurosci. Methods 172, 137–141 (2008).
14. Zang, Y., Jiang, T., Lu, Y., He, Y. & Tian, L. Regional homogeneity approach to fMRI data analysis. Neuroimage 22, 394–400 (2004).
15. Yeo, B. T. T. et al. The organization of the human cerebral cortex estimated by intrinsic functional connectivity. J. Neurophysiol. 106, 1125–1165 (2011).

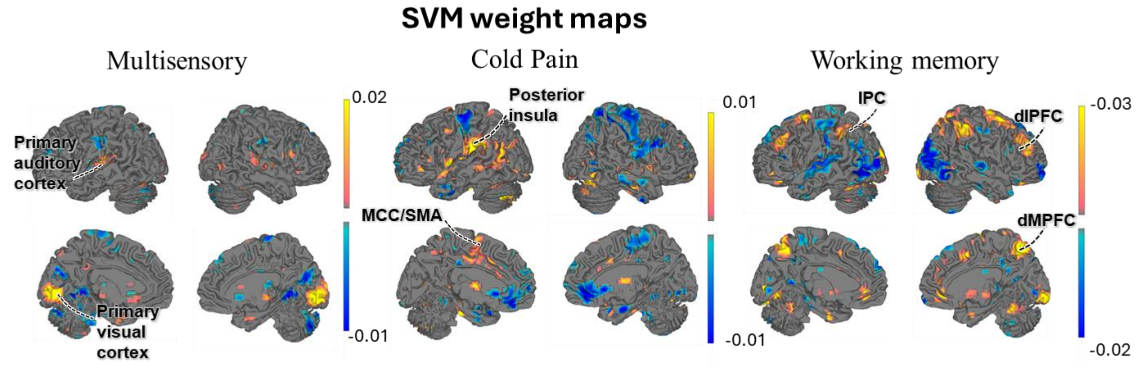


**Figure S1. Uncorrected task-discriminating SVM weight maps, related to Figure 2.** Maps are displayed at $p < 0.05$ uncorrected to provide a more inclusive visualization of the distributed task-related SVM patterns. These maps are intended for visualization only. Classification and individual-level brain pattern-expression scores were computed using the full, unthresholded group-level SVM weight map. Compared with the FDR-corrected maps shown in Figure 2B, the uncorrected maps show more spatially distributed pattern information but should be interpreted less robustly.

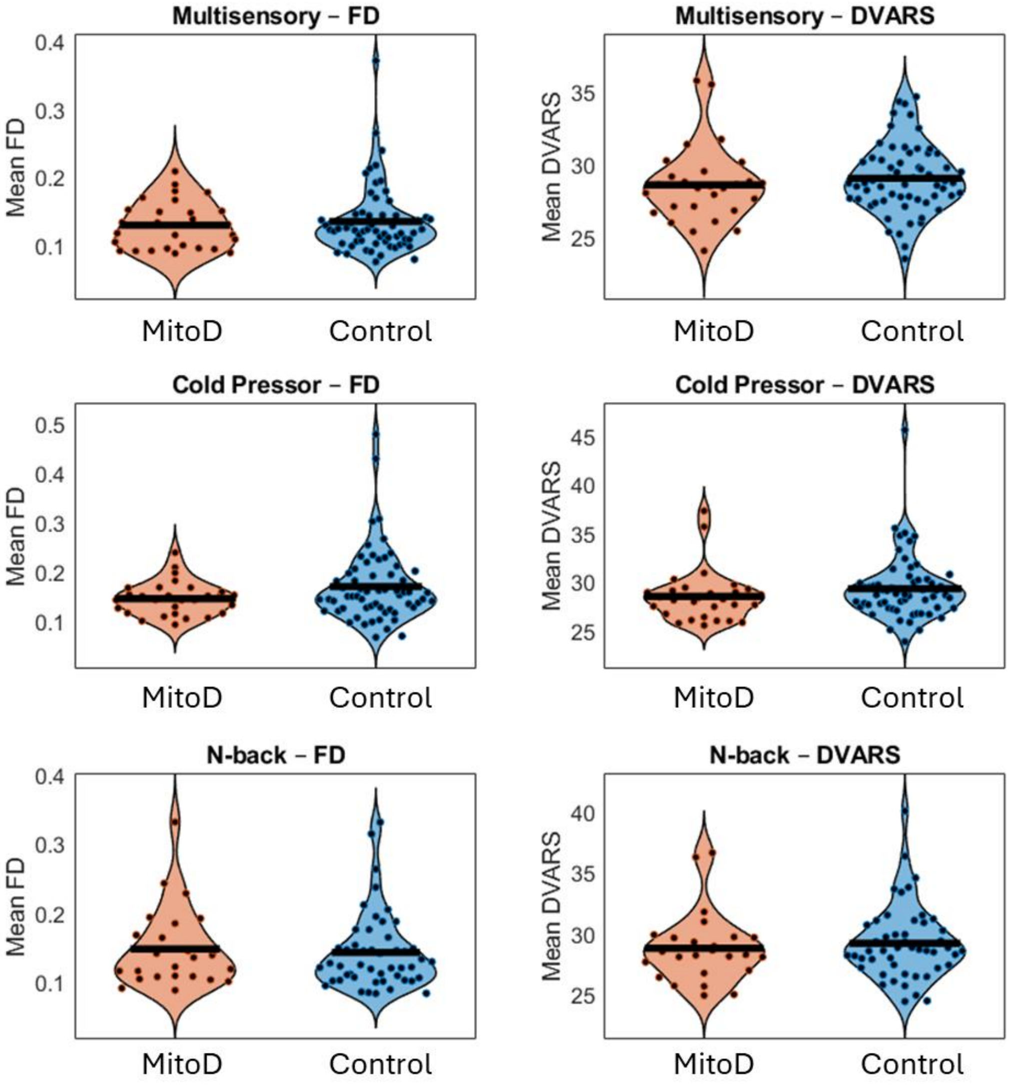


**Figure S2. Head-motion quality control across tasks, related to Figure 2 and Supplementary Methods.** Data QC: quantitative QC indices per task and group: mean/median FD, DVARS. The statistics: Multisensory: FD: $t(88) = -0.528$, $p = 0.5989$; DVARS: $t(88) = -0.818$, $p = 0.4158$; Cold Pressor; FD: $t(89) = -1.696$, $p = 0.0933$; DVARS: $t(89) = -1.151$, $p = 0.2526$; N-back, FD: $t(72) = 0.373$, $p = 0.7103$; DVARS: $t(72) = -0.561$, $p = 0.5762$.These results show that groups are balanced on head movement and movement-related brain features, and thus group differences are unlikely to be driven by movement.

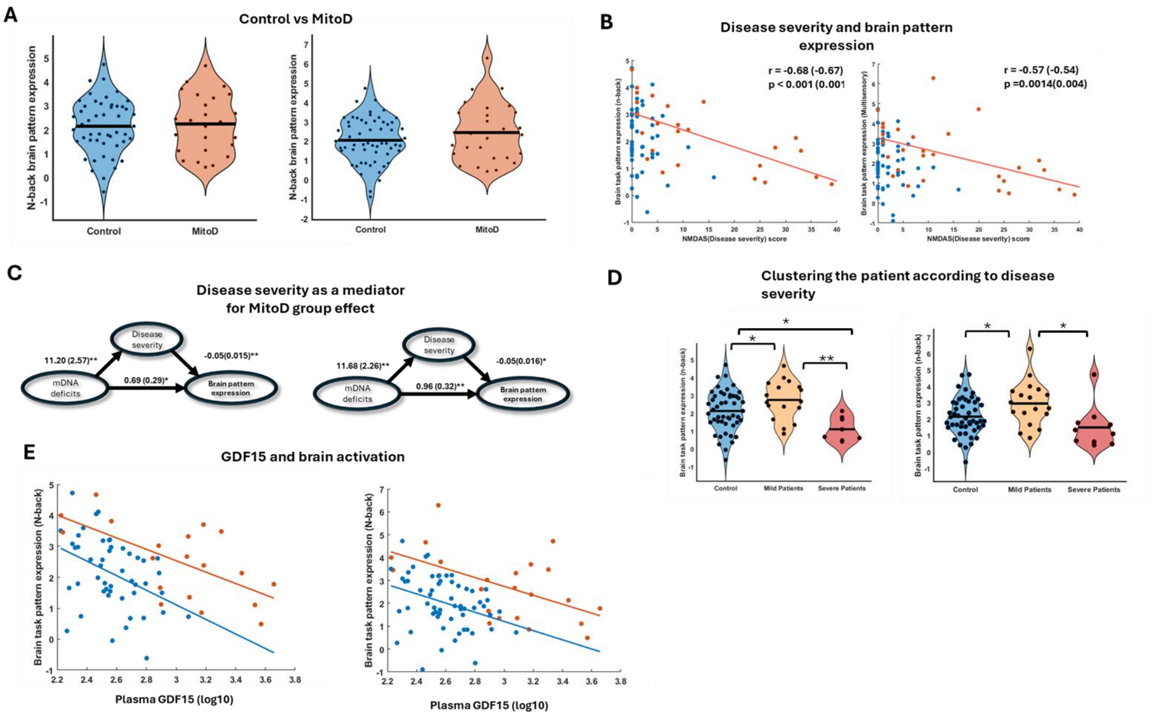


**Figure S3. Sensitivity to the N-back behavioral data-quality/task-engagement criterion, related to Figures 3 and 4.** Sensitivity analyses for all analysis figures involving brain-pattern expression after excluding participants with behavioral data-quality or task-engagement failures (left), compared with results before exclusion (right). A) Analysis for Figure S7A. B) Analysis for Figure 3C. C) Analysis for Figure 3D. D) Analysis for Figure 3E. E) Analysis for Figure 5B. The overall pattern of results was similar when the excluded participants were retained, although the severe-patient versus control comparison of N-back brain pattern expression in Figure 3E was no longer significant. *p <0.05. ** p<0.001

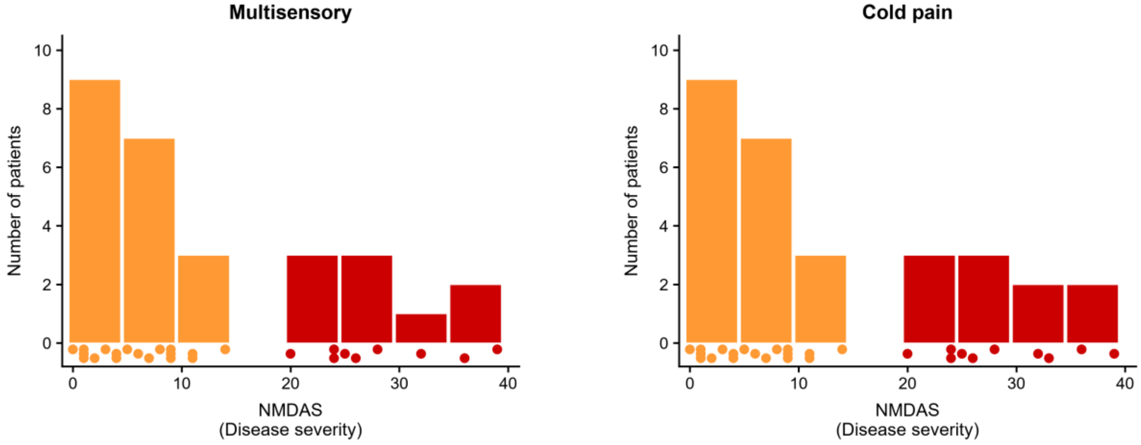


**Figure S4. NMDAS distributions in the multisensory and cold-pain samples, related to Figure 3.** Histograms show NMDAS scores in 5-point bins, with individual observations below. Amber and red indicate the resilient and severe patient subgroups identified using K-means clustering.

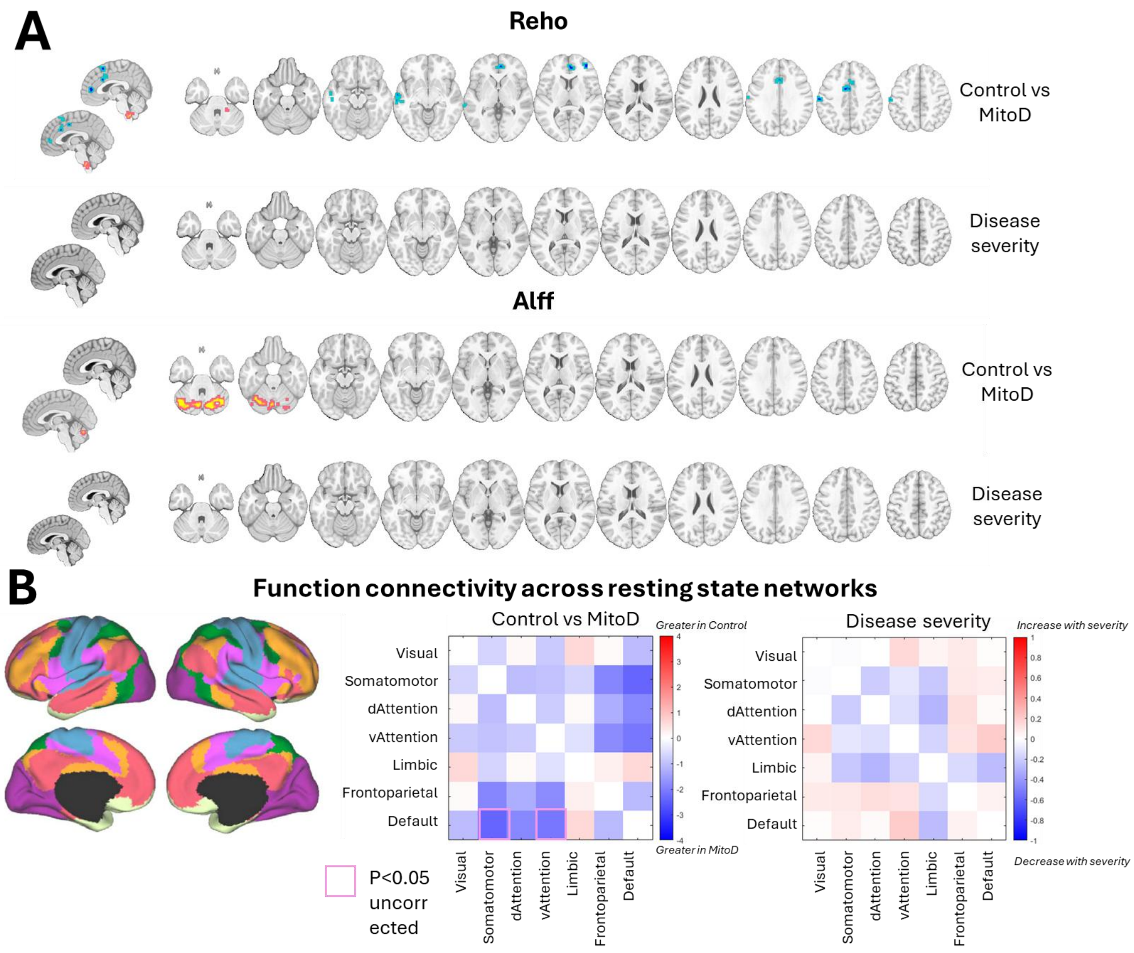


**Figure S5. Regional activity and network connectivity at rest, related to Figure 3 and Supplementary Methods.** A) Group comparison maps for Regional Homogeneity (ReHo) and Amplitude of Low Frequency Fluctuations (ALFF) between mitochondrial disease (MitoD) patients and healthy controls (top row), as well as correlations with disease severity within the MitoD group (bottom row). Voxel-wise threshold set at uncorrected p < 0.001 with cluster size > 20. No clusters survived FDR correction. B) Functional connectivity analysis across seven canonical resting-state networks defined by the Yeo et al. 16 atlas (left). Group-level connectivity matrices (right) show between-network connectivity for controls vs MitoD (middle) and associations with disease severity (right). Most effects did not reach statistical significance, indicating minimal alterations in resting-state connectivity in MitoD.

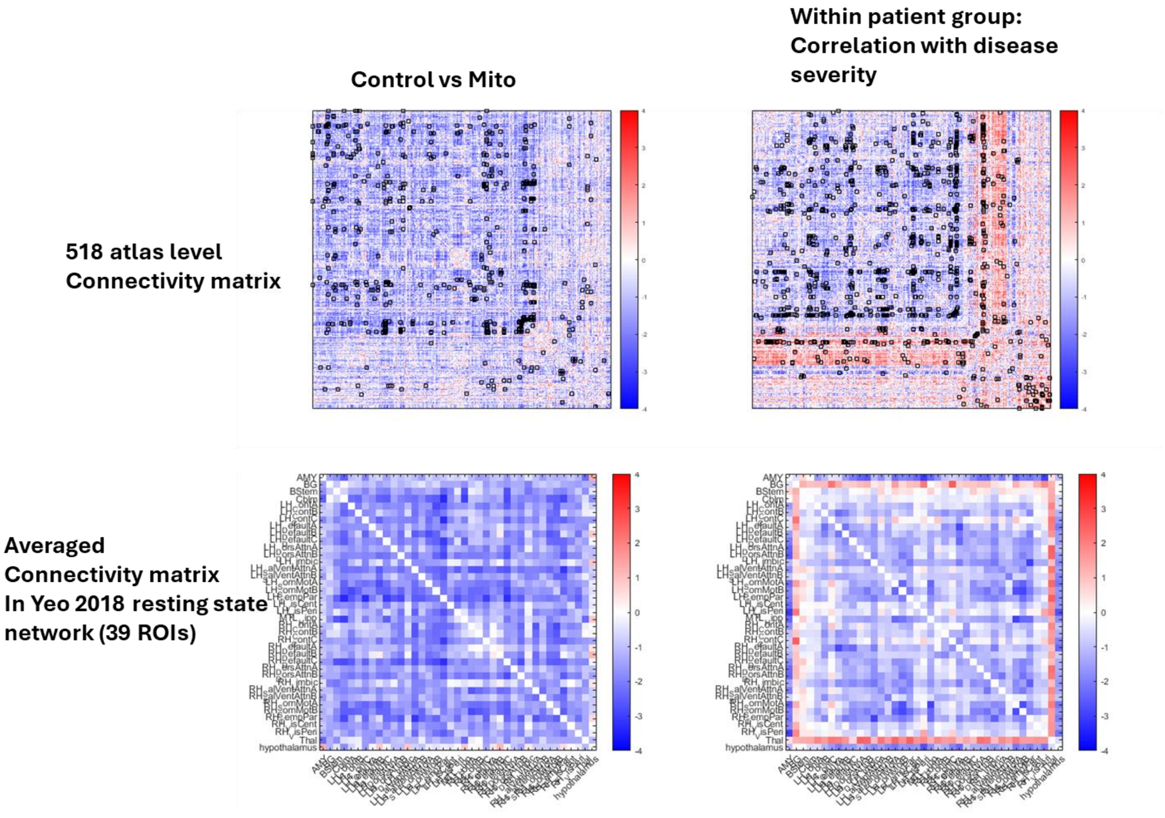


**Figure S6. Resting-state connectivity across 518 parcels, related to Figure 3 and Supplementary Methods.** A) Functional connectivity analysis was conducted across 518 atlas-defined parcels from the CANlab 2024 atlas (https://canlab.github.io/_pages/using_canlab_atlases/using_canlab_atlases.html). Group-level connectivity matrices show between-network connectivity for controls versus MitoD participants, and associations with disease severity. No results survived FDR correction. Effects with $p < 0.001$ are shown with black contours. B) Results from panel A were averaged within resting-state networks defined by Yeo et al. 16. No results survived FDR correction at $q < 0.05$ or the uncorrected threshold of $p < 0.001$.

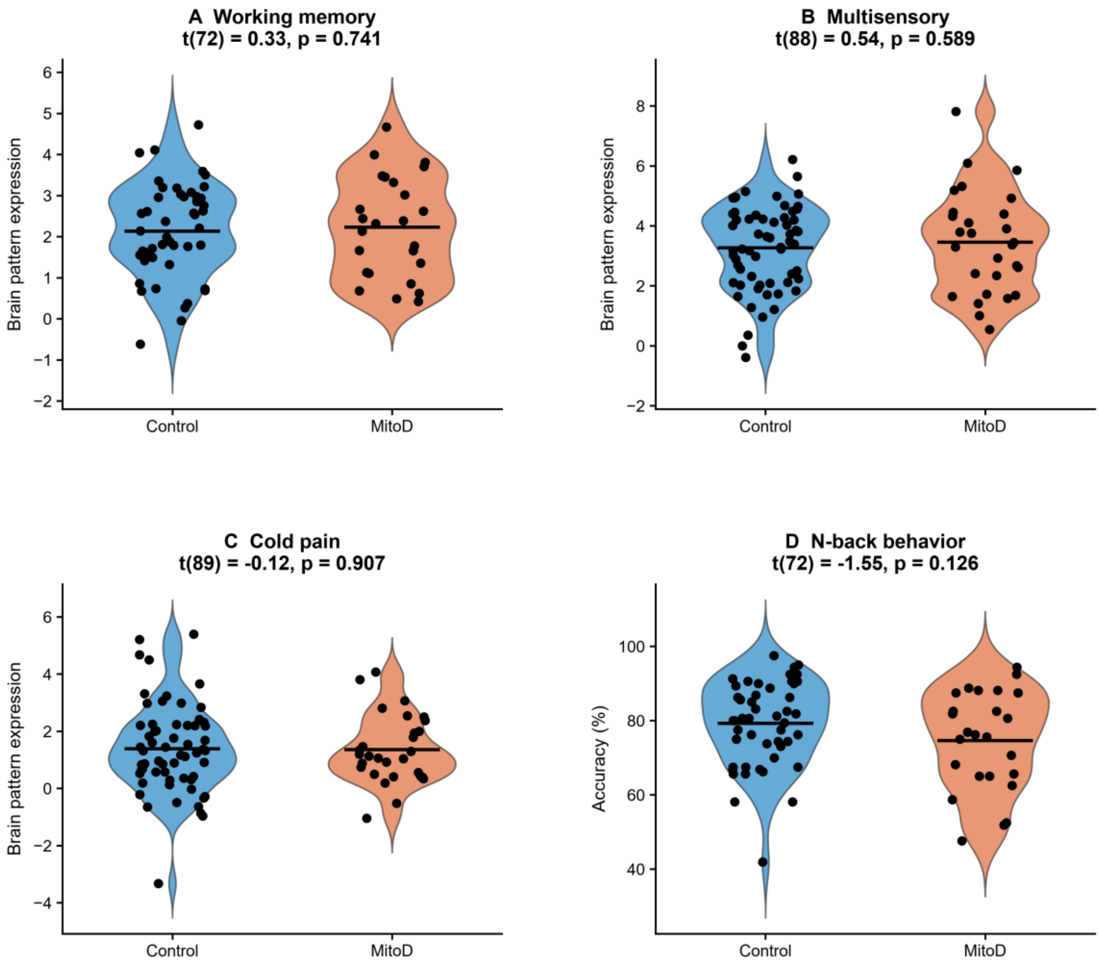


**Figure S7. Overall MitoD–control comparisons, related to Figures 2 and 3.** A–C) Task-minus-control brain pattern expression for working memory (25 MitoD, 49 controls), multisensory stimulation (28 MitoD, 62 controls), and cold pain (29 MitoD, 62 controls). D) N-back accuracy (25 MitoD, 49 controls). Blue denotes controls and orange denotes MitoD; dots represent participants and horizontal bars indicate means. Two-sided pooled-variance t tests compare MitoD minus control: working memory, $t(72) = 0.33$, $p = 0.741$; multisensory, $t(88) = 0.54$, $p = 0.589$; cold pain, $t(89) = -0.12$, $p = 0.907$; and accuracy, $t(72) = -1.55$, $p = 0.126$. The N-back eligibility criterion is retained. Null group means coexist with severity-related variation within patients.

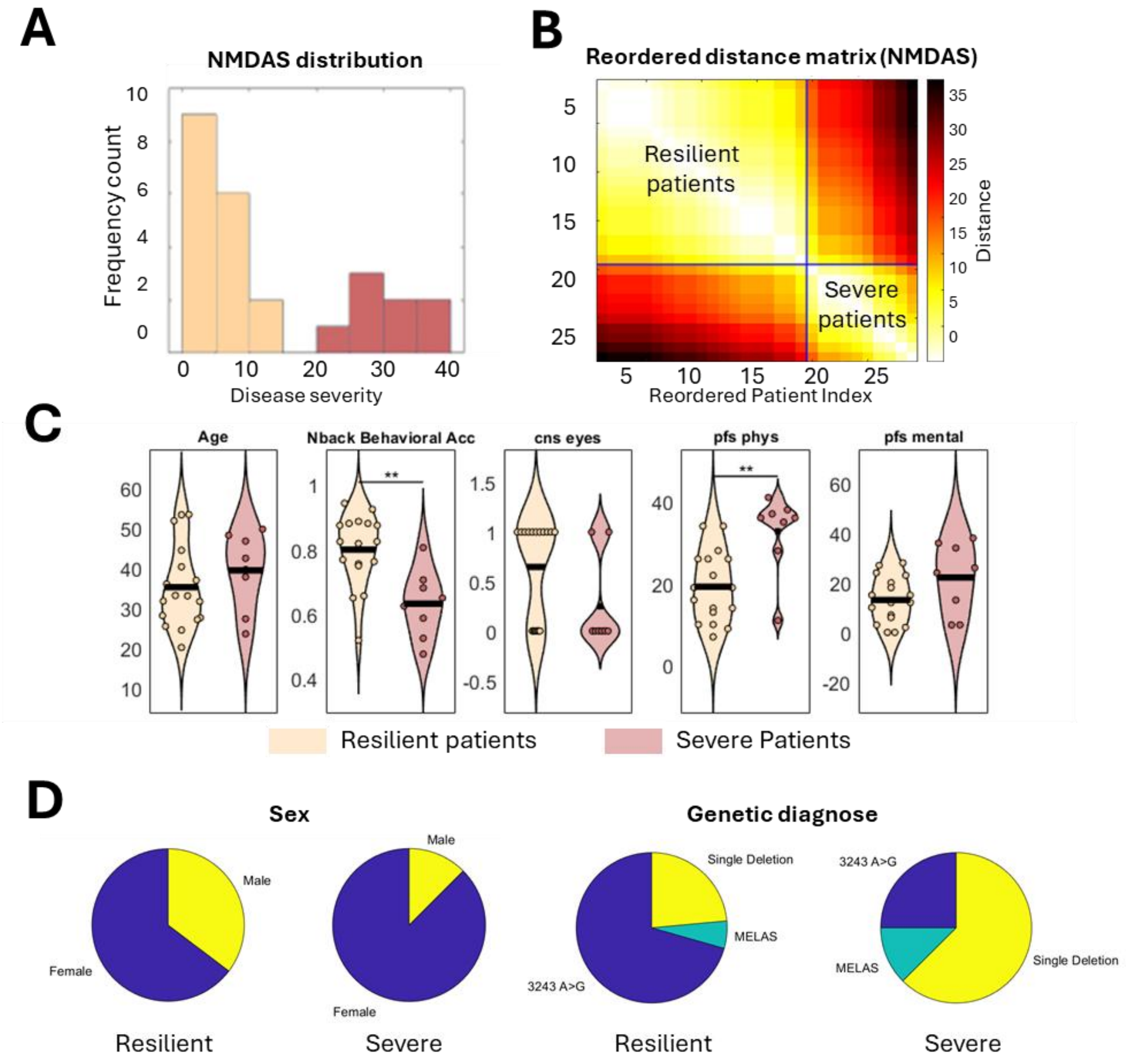


**Figure S8. Characteristics of resilient and severe MitoD subgroups, related to Figure 3.** A) distribution of NMDAS disease severity. B) Reordered pairwise distance matrix for the NMDAS-based clustering solution. Each cell represents the absolute difference in NMDAS score between pairs of patients, reordered by cluster assignment and sorted by NMDAS score within each cluster. C) Numerical basic information and important variables across each subgroup. ** p<0.05 under two-sample T test. D. Categorical basic information and important variables across each subgroup. These results indicate a significant difference in NMDAS score between the severe and resilient groups. Severe patients showed significantly higher physical fatigue and lower N-back task performance, whereas no significant differences were observed between groups in age, sex, genotype, general medical eye examination item, or mental fatigue. CNS eye: general medical eye examination item in columbia neurological score; Pfs phys, phs mental: Physical and mental fatigue.

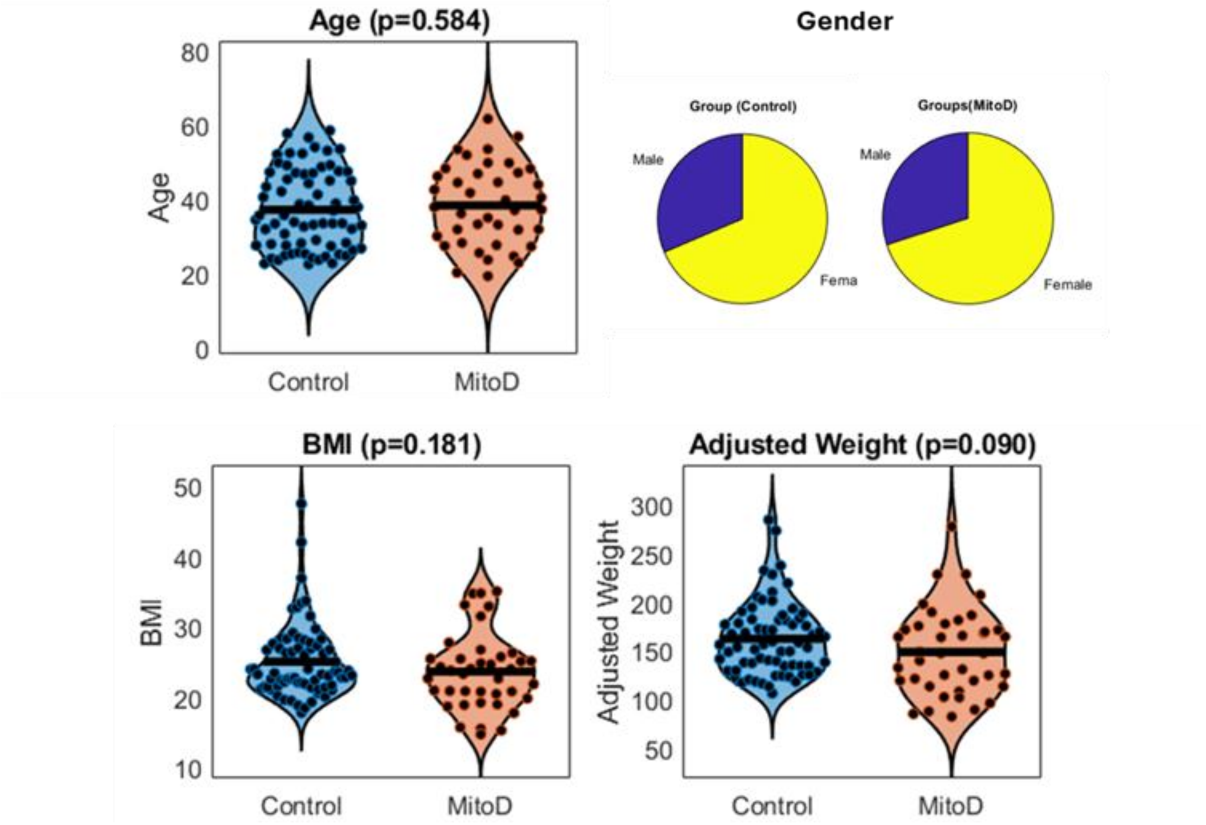


**Figure S9. Demographic and anthropometric characteristics, related to Figures 1–3 and Supplementary Methods.** Violin plots compare age, BMI, and adjusted weight between control and MitoD participants. Pie charts depict the sex distribution—percentage of males and females—in each group.

Supplemental tables

Table S1. Phenome-wide group comparisons, related to Figure 1.

g = Hedges' g (bias-corrected); M = mean; SD = standard deviation. p values are from Welch's t tests; q values are Benjamini–Hochberg FDR-adjusted across 211 testable comparisons. All 214 measurements are included, including those without significant group differences. Dashes denote unavailable statistics.

| Variable | Category | g | M (Controls) | SD (Controls) | N (Controls) | M (Patients) | SD (Patients) | N (Patients) | p | q (FDR) |
|---|---|---|---|---|---|---|---|---|---|---|
| stai_y1_state_total | Psychological Factors | 0.10 | 36.54 | 12.44 | 69 | 35.33 | 9.67 | 40 | 0.5729 | 0.7370 |
| stai_y1_trait_total | Psychological Factors | -0.06 | 40.54 | 12.84 | 69 | 41.23 | 10.73 | 40 | 0.7647 | 0.8866 |
| bdi_total | Psychological Factors | -0.27 | 7.94 | 8.32 | 69 | 10.25 | 8.78 | 40 | 0.1816 | 0.3515 |
| mbi_total | Psychological Factors | -0.23 | 6.49 | 2.43 | 69 | 7.05 | 2.42 | 39 | 0.2549 | 0.4337 |
| mbi_ineff | Psychological Factors | -0.05 | 4.33 | 1.53 | 69 | 4.41 | 1.47 | 39 | 0.7980 | 0.9101 |
| mbi_cyn | Psychological Factors | -0.00 | 1.92 | 1.45 | 69 | 1.93 | 1.52 | 39 | 0.9828 | 0.9876 |
| mbi_exhaus | Psychological Factors | -0.44 | 1.96 | 1.44 | 69 | 2.62 | 1.59 | 39 | 0.0349 | 0.1252 |
| pcl_total | Psychological Factors | -0.23 | 30.01 | 12.33 | 69 | 32.83 | 11.40 | 40 | 0.2319 | 0.4130 |
| dsm_5_cross_cutting | Psychological Factors | -0.03 | 12.14 | 12.23 | 63 | 12.54 | 9.89 | 35 | 0.8608 | 0.9509 |
| dsm_5_neg_aff | Psychological Factors | -0.21 | 3.38 | 3.29 | 66 | 4.05 | 2.95 | 39 | 0.2821 | 0.4651 |
| dsm_5_detachment | Psychological Factors | 0.13 | 2.85 | 3.00 | 67 | 2.46 | 2.90 | 39 | 0.5126 | 0.6717 |
| dsm_5_antagonism | Psychological Factors | 0.05 | 1.41 | 2.13 | 68 | 1.32 | 1.28 | 38 | 0.7725 | 0.8907 |
| dsm_5_disinhibition | Psychological Factors | 0.29 | 2.59 | 3.31 | 68 | 1.69 | 2.49 | 39 | 0.1169 | 0.2602 |
| dsm_5_psychoticism | Psychological Factors | 0.14 | 2.35 | 3.24 | 69 | 1.95 | 2.22 | 38 | 0.4525 | 0.6348 |
| dsm_5_substance | Psychological Factors | -0.56 | 1.71 | 1.89 | 7 | 3.33 | 4.04 | 3 | 0.5654 | 0.7319 |
| dsm_5_somatic | Psychological Factors | -0.33 | 10.12 | 5.91 | 8 | 12.00 | 5.25 | 14 | 0.4695 | 0.6348 |
| dsm_5_sleep_dist | Psychological Factors | -0.03 | 27.47 | 6.84 | 19 | 27.71 | 7.08 | 14 | 0.9227 | 0.9678 |
| dsm_5_repetitiv | Psychological Factors | 1.56 | 10.40 | 3.84 | 10 | 3.67 | 4.73 | 3 | 0.1143 | 0.2593 |
| dsm_5_mania | Psychological Factors | -0.50 | 8.90 | 3.55 | 21 | 10.83 | 4.31 | 6 | 0.3488 | 0.5333 |
| dsm_5_depre | Psychological Factors | 0.57 | 24.00 | 6.01 | 22 | 19.93 | 8.32 | 15 | 0.1171 | 0.2602 |
| dsm_5_anx | Psychological Factors | 0.25 | 20.61 | 6.06 | 23 | 19.06 | 6.29 | 16 | 0.4489 | 0.6348 |
| dsm_5_anger | Psychological Factors | 0.05 | 14.33 | 4.36 | 12 | 14.10 | 4.68 | 10 | 0.9056 | 0.9578 |
| pss_total | Psychological Factors | -0.03 | 17.14 | 7.35 | 69 | 17.35 | 6.26 | 40 | 0.8775 | 0.9577 |
| tics_total_score | Psychological Factors | -0.36 | 45.20 | 19.28 | 69 | 51.98 | 17.27 | 40 | 0.0620 | 0.1700 |
| dhs_score | Psychological Factors | -0.29 | 64.57 | 17.95 | 69 | 70.00 | 18.97 | 40 | 0.1455 | 0.3039 |
| neo_neuro | Psychological Factors | 0.09 | 33.74 | 10.33 | 69 | 32.85 | 8.99 | 40 | 0.6390 | 0.7931 |
| neo_extra | Psychological Factors | 0.28 | 40.71 | 6.45 | 69 | 38.80 | 7.21 | 40 | 0.1703 | 0.3422 |
| neo_open | Psychological Factors | 0.52 | 45.17 | 7.30 | 69 | 41.33 | 7.35 | 40 | 0.0099 | 0.0464 |
| neo_agree | Psychological Factors | 0.02 | 43.65 | 5.55 | 69 | 43.52 | 4.41 | 40 | 0.8956 | 0.9578 |
| neo_cons | Psychological Factors | 0.00 | 44.61 | 8.44 | 69 | 44.58 | 7.57 | 40 | 0.9829 | 0.9876 |
| mpss_score | Social and Environmental | -0.15 | 82.13 | 12.53 | 69 | 84.20 | 15.80 | 40 | 0.4808 | 0.6403 |

| Variable | Category | g | M (Controls) | SD (Controls) | N (Controls) | M (Patients) | SD (Patients) | N (Patients) | p | q (FDR) |
|---|---|---|---|---|---|---|---|---|---|---|
| ssq_s | Social and Environmental | -0.23 | 30.24 | 6.92 | 70 | 31.69 | 4.84 | 35 | 0.2179 | 0.3997 |
| ssq_n | Social and Environmental | -0.41 | 22.40 | 12.12 | 70 | 27.38 | 12.31 | 40 | 0.0436 | 0.1436 |
| ctq_ea | Social and Environmental | -0.02 | 8.62 | 4.53 | 69 | 8.69 | 4.60 | 39 | 0.9401 | 0.9723 |
| ctq_pa | Social and Environmental | -0.02 | 6.10 | 2.20 | 69 | 6.15 | 2.24 | 39 | 0.9069 | 0.9578 |
| ctq_sa | Social and Environmental | -0.03 | 6.04 | 3.55 | 69 | 6.15 | 3.71 | 39 | 0.8805 | 0.9577 |
| ctq_en | Social and Environmental | 0.18 | 10.78 | 5.88 | 69 | 9.79 | 4.76 | 39 | 0.3447 | 0.5333 |
| ctq_pn | Social and Environmental | 0.21 | 7.01 | 2.79 | 69 | 6.46 | 2.22 | 39 | 0.2613 | 0.4410 |
| ctq_md | Social and Environmental | -0.04 | 0.45 | 0.93 | 69 | 0.49 | 0.79 | 39 | 0.8232 | 0.9338 |
| leq_positive_events_score | Social and Environmental | 0.12 | 11.81 | 10.32 | 57 | 10.60 | 10.16 | 30 | 0.6022 | 0.7608 |
| leq_negative_events_score | Social and Environmental | 0.14 | 6.49 | 7.54 | 61 | 5.50 | 6.13 | 36 | 0.4825 | 0.6403 |
| leq_total_events_score | Social and Environmental | 0.07 | 22.54 | 15.69 | 67 | 21.31 | 19.38 | 39 | 0.7371 | 0.8593 |
| l_cat | Health Behaviors | 0.38 | 3.60 | 1.32 | 70 | 3.08 | 1.47 | 40 | 0.0663 | 0.1771 |
| ipaq_total_met | Health Behaviors | -0.05 | 4145.68 | 3739.07 | 69 | 4370.09 | 5866.84 | 39 | 0.8302 | 0.9362 |
| ipaq_intensity | Health Behaviors | 0.33 | 2.39 | 0.67 | 70 | 2.15 | 0.80 | 40 | 0.1200 | 0.2610 |
| scor_abp1_psqi_global | Health Behaviors | -0.37 | 6.09 | 3.38 | 68 | 7.46 | 4.22 | 39 | 0.0868 | 0.2180 |
| mfis_phy_sub | Health/Symptoms | -1.23 | 6.96 | 6.63 | 70 | 16.62 | 9.57 | 40 | 0.0000 | 0.0000 |
| mfis_cog_sub | Health/Symptoms | -1.00 | 8.30 | 7.60 | 70 | 16.75 | 9.60 | 40 | 0.0000 | 0.0001 |
| mfis_psy_sub | Health/Symptoms | -0.99 | 1.94 | 1.91 | 70 | 3.98 | 2.25 | 40 | 0.0000 | 0.0001 |
| mfis_total | Health/Symptoms | -1.19 | 17.20 | 15.12 | 70 | 37.35 | 19.35 | 40 | 0.0000 | 0.0000 |
| compass_orth_int | Health/Symptoms | -0.43 | 6.46 | 8.12 | 70 | 10.00 | 8.50 | 40 | 0.0357 | 0.1252 |
| compass_vasomotor | Health/Symptoms | -0.50 | 0.07 | 0.42 | 70 | 0.46 | 1.15 | 40 | 0.0456 | 0.1480 |
| compass_secretomotor | Health/Symptoms | -0.90 | 1.50 | 2.14 | 70 | 3.80 | 3.13 | 40 | 0.0001 | 0.0009 |
| compass_gastrointest | Health/Symptoms | -1.13 | 3.25 | 2.95 | 70 | 6.74 | 3.27 | 40 | 0.0000 | 0.0000 |
| compass_bladded | Health/Symptoms | -0.25 | 0.51 | 1.03 | 70 | 0.78 | 1.19 | 40 | 0.2329 | 0.4130 |
| compass_pupillomotor | Health/Symptoms | -1.30 | 0.85 | 0.90 | 70 | 2.28 | 1.36 | 40 | 0.0000 | 0.0000 |
| compass_total | Health/Symptoms | -1.03 | 12.64 | 10.19 | 70 | 24.06 | 12.34 | 40 | 0.0000 | 0.0001 |
| bipq_cog_ill_rep | Health/Symptoms | — | — | — | 0 | 26.73 | 7.90 | 37 | — | — |
| bipq_emo_rep | Health/Symptoms | — | — | — | 0 | 10.24 | 4.60 | 37 | — | — |
| bipq_ill_compreh | Health/Symptoms | — | — | — | 0 | 3.51 | 2.49 | 37 | — | — |
| pfs_phys | Health/Symptoms | -0.91 | 15.34 | 8.02 | 70 | 23.32 | 9.92 | 40 | 0.0000 | 0.0004 |

| Variable | Category | g | M (Controls) | SD (Controls) | N (Controls) | M (Patients) | SD (Patients) | N (Patients) | p | q (FDR) |
|---|---|---|---|---|---|---|---|---|---|---|
| pfs_mental | Health/Symptoms | -0.30 | 14.03 | 9.33 | 70 | 17.07 | 11.57 | 40 | 0.1596 | 0.3301 |
| mcc_score | Health/Symptoms | -0.21 | 74.99 | 34.06 | 69 | 82.30 | 36.92 | 40 | 0.3084 | 0.4968 |
| dcf_sit_stand_time | Symptoms/Clinical Assessmnent | 0.70 | 21.29 | 6.21 | 69 | 16.61 | 7.40 | 38 | 0.0015 | 0.0099 |
| dcf_bodyfat_avg | Symptoms/Clinical Assessmnent | 0.01 | 27.91 | 9.98 | 67 | 27.84 | 9.91 | 40 | 0.9720 | 0.9876 |
| fat_free_mass | Symptoms/Clinical Assessmnent | 0.14 | 47.56 | 25.90 | 67 | 43.87 | 24.62 | 40 | 0.4639 | 0.6348 |
| hip_waist_circum_ratio | Symptoms/Clinical Assessmnent | 0.20 | 1.14 | 0.10 | 69 | 1.12 | 0.13 | 40 | 0.3573 | 0.5410 |
| dcg_adj_wgt | Symptoms/Clinical Assessmnent | 0.34 | 162.22 | 37.01 | 70 | 148.77 | 43.96 | 40 | 0.1072 | 0.2542 |
| fev1_average | Symptoms/Clinical Assessmnent | 0.42 | 3.19 | 0.68 | 69 | 2.88 | 0.85 | 38 | 0.0569 | 0.1670 |
| fev1_predicted_average | Symptoms/Clinical Assessmnent | 0.59 | 93.34 | 16.08 | 69 | 83.91 | 15.67 | 38 | 0.0042 | 0.0232 |
| fvc_average | Symptoms/Clinical Assessmnent | 0.29 | 3.96 | 0.80 | 69 | 3.69 | 1.16 | 38 | 0.1946 | 0.3700 |
| fvc_predicted_average | Symptoms/Clinical Assessmnent | 0.52 | 95.40 | 15.85 | 69 | 87.23 | 15.40 | 38 | 0.0111 | 0.0500 |
| fev1fvc_average | Symptoms/Clinical Assessmnent | 0.45 | 80.62 | 6.89 | 69 | 77.34 | 7.95 | 38 | 0.0362 | 0.1252 |
| fev1_fvc_1_predicted_average | Symptoms/Clinical Assessmnent | 0.45 | 80.62 | 6.89 | 69 | 77.34 | 7.95 | 38 | 0.0362 | 0.1252 |
| met_ree_d2 | Systemic Energy Metabolism | 0.33 | 1569.58 | 399.73 | 45 | 1439.16 | 376.07 | 19 | 0.2216 | 0.4031 |
| met_vo2_d2 | Systemic Energy Metabolism | 0.45 | 228.96 | 58.43 | 45 | 200.26 | 71.29 | 19 | 0.1324 | 0.2822 |
| respiratory_rate_rr_breaths_min | Systemic Energy Metabolism | 0.48 | 12.07 | 4.47 | 42 | 9.79 | 5.11 | 18 | 0.1122 | 0.2593 |
| met_feo2_d2 | Systemic Energy Metabolism | -0.07 | 16.64 | 0.76 | 45 | 16.69 | 1.12 | 19 | 0.8341 | 0.9362 |
| met_tidal_d2 | Systemic Energy Metabolism | -0.37 | 0.72 | 0.35 | 45 | 0.86 | 0.42 | 19 | 0.2095 | 0.3877 |
| met_ve_d2 | Systemic Energy Metabolism | 0.04 | 7.47 | 2.61 | 45 | 7.37 | 2.68 | 19 | 0.8902 | 0.9578 |
| met_vo2_kg_d2 | Systemic Energy Metabolism | -0.06 | 3.11 | 0.75 | 45 | 3.16 | 1.03 | 19 | 0.8575 | 0.9509 |
| vf_c1_lf_cor_scaled | Cognitive Processes | 0.33 | 11.70 | 3.36 | 67 | 10.53 | 3.94 | 40 | 0.1194 | 0.2610 |
| vf_c1_cf_cor_scaled | Cognitive Processes | 0.19 | 11.70 | 3.47 | 67 | 11.00 | 3.85 | 40 | 0.3473 | 0.5333 |
| vf_c1_cs_cor_scaled | Cognitive Processes | 0.40 | 12.33 | 3.70 | 67 | 10.72 | 4.36 | 39 | 0.0569 | 0.1670 |
| vf_c1_cs_tsa_scaled | Cognitive Processes | 0.21 | 12.06 | 3.19 | 67 | 11.36 | 3.67 | 39 | 0.3233 | 0.5163 |
| nab_shl_irg_perc | Cognitive Processes | 0.09 | 65.91 | 24.78 | 67 | 63.72 | 25.00 | 39 | 0.6634 | 0.8045 |

| Variable | Category | g | M (Controls) | SD (Controls) | N (Controls) | M (Patients) | SD (Patients) | N (Patients) | p | q (FDR) |
|---|---|---|---|---|---|---|---|---|---|---|
| nab_shl_drg_perc | Cognitive Processes | 0.40 | 62.74 | 24.26 | 66 | 52.10 | 29.11 | 39 | 0.0587 | 0.1672 |
| rbans_cd_scaled | Cognitive Processes | 0.46 | 9.62 | 3.36 | 66 | 7.77 | 4.94 | 39 | 0.0426 | 0.1425 |
| rbans_ll_scaled | Cognitive Processes | 0.14 | 10.81 | 3.21 | 67 | 10.35 | 3.39 | 40 | 0.4945 | 0.6521 |
| nab_snl_spd_perc | Cognitive Processes | 0.48 | 51.82 | 29.21 | 66 | 37.22 | 32.71 | 36 | 0.0288 | 0.1126 |
| tm_ns_scaled | Cognitive Processes | 0.64 | 10.74 | 2.12 | 66 | 8.75 | 4.19 | 40 | 0.0073 | 0.0364 |
| tm_ls_scaled | Cognitive Processes | 0.52 | 10.93 | 2.80 | 67 | 9.18 | 4.04 | 40 | 0.0186 | 0.0755 |
| tm_nls_scaled | Cognitive Processes | 0.41 | 10.42 | 3.07 | 67 | 8.97 | 4.08 | 40 | 0.0574 | 0.1670 |
| nab_sdgf_perc | Cognitive Processes | 0.43 | 54.48 | 30.06 | 67 | 41.74 | 29.11 | 39 | 0.0349 | 0.1252 |
| nab_sdgb_perc | Cognitive Processes | 0.23 | 58.36 | 30.15 | 66 | 51.41 | 31.39 | 39 | 0.2692 | 0.4508 |
| cwi_cn_scaled | Cognitive Processes | 0.26 | 9.06 | 3.13 | 67 | 8.18 | 3.66 | 40 | 0.2065 | 0.3877 |
| cwi_wr_scaled | Cognitive Processes | 0.58 | 10.31 | 2.61 | 67 | 8.57 | 3.57 | 40 | 0.0092 | 0.0453 |
| cwi_inhib_scaled | Cognitive Processes | 0.48 | 11.10 | 2.83 | 67 | 9.53 | 3.88 | 40 | 0.0285 | 0.1126 |
| cwi_inh_cn_scaled | Cognitive Processes | 0.25 | 11.78 | 3.27 | 67 | 11.03 | 2.44 | 40 | 0.1796 | 0.3515 |
| st_free_corsort_scaled | Cognitive Processes | -0.27 | 11.63 | 2.97 | 67 | 12.47 | 3.24 | 34 | 0.2086 | 0.3877 |
| topf_standrd_score | Cognitive Processes | 0.08 | 110.63 | 12.39 | 67 | 109.62 | 12.67 | 39 | 0.6905 | 0.8278 |
| wasi_full_sum_c | Cognitive Processes | 0.10 | 109.99 | 15.35 | 67 | 108.38 | 16.69 | 40 | 0.6204 | 0.7745 |
| nmdas_i_ii_iii_score | Symptoms/Clinical Assessmnent | -1.65 | 1.97 | 2.91 | 68 | 15.88 | 13.26 | 40 | 0.0000 | 0.0000 |
| cns_score_total | Symptoms/Clinical Assessmnent | 1.61 | 72.85 | 1.84 | 68 | 65.41 | 7.19 | 40 | 0.0000 | 0.0000 |
| karnofsky_100 | Symptoms/Clinical Assessmnent | 1.83 | 98.67 | 4.05 | 45 | 88.75 | 8.06 | 16 | 0.0002 | 0.0014 |
| psychophysiology_arm_2.hr_avg_p5_300s | Stress Reactivity Systems | -0.42 | 74.60 | 11.18 | 55 | 79.65 | 13.14 | 37 | 0.0594 | 0.1672 |
| psychophysiology_arm_2.hr_avg_p10_300s | Stress Reactivity Systems | -0.31 | 74.45 | 10.72 | 58 | 78.13 | 13.04 | 35 | 0.1640 | 0.3328 |
| psychophysiology_arm_2.hr_avg_p5_60s_1 | Stress Reactivity Systems | -0.22 | 78.18 | 13.92 | 67 | 81.38 | 15.86 | 39 | 0.2989 | 0.4889 |
| psychophysiology_arm_2.hr_avg_p5_60s_2 | Stress Reactivity Systems | -0.34 | 73.93 | 13.19 | 67 | 78.52 | 13.70 | 37 | 0.1015 | 0.2435 |
| psychophysiology_arm_2.respr_count_p5_60s_1 | Stress Reactivity Systems | 0.17 | 19.78 | 4.17 | 68 | 19.06 | 4.12 | 34 | 0.4071 | 0.5883 |
| psychophysiology_arm_2.respr_count_p5_60s_2 | Stress Reactivity Systems | 0.15 | 18.92 | 4.11 | 68 | 18.30 | 4.11 | 35 | 0.4723 | 0.6348 |
| psychophysiology_arm_2.hr_avg_p10_60s_1 | Stress Reactivity Systems | -0.24 | 75.51 | 12.39 | 69 | 78.52 | 12.69 | 39 | 0.2363 | 0.4156 |
| psychophysiology_arm_2.hr_avg_p10_60s_2 | Stress Reactivity Systems | -0.31 | 73.14 | 11.68 | 65 | 76.86 | 12.69 | 39 | 0.1403 | 0.2960 |
| psychophysiology_arm_2.respr_count_p10_60s_1 | Stress Reactivity Systems | -0.16 | 19.59 | 3.72 | 68 | 20.21 | 4.22 | 35 | 0.4655 | 0.6348 |
| psychophysiology_arm_2.respr_count_p10_60s_2 | Stress Reactivity Systems | 0.00 | 19.09 | 3.68 | 68 | 19.10 | 4.22 | 36 | 0.9912 | 0.9912 |

| Variable | Category | g | M (Controls) | SD (Controls) | N (Controls) | M (Patients) | SD (Patients) | N (Patients) | p | q (FDR) |
|---|---|---|---|---|---|---|---|---|---|---|
| biological_sample_arm_2b.ln_nd1_copies_ul_saliva_m5 | Energetics | 0.47 | 7.04 | 1.93 | 60 | 6.13 | 1.84 | 30 | 0.0337 | 0.1252 |
| biological_sample_arm_2b.ln_b2m_copies_ul_saliva_m5 | Energetics | 0.44 | 3.88 | 1.68 | 55 | 3.14 | 1.56 | 23 | 0.0701 | 0.1804 |
| biological_sample_arm_2b.ln_nd1_copies_ul_saliva_5 | Energetics | 0.41 | 8.14 | 1.69 | 60 | 7.45 | 1.75 | 33 | 0.0662 | 0.1771 |
| biological_sample_arm_2b.ln_b2m_copies_ul_saliva_5 | Energetics | 0.53 | 4.66 | 1.78 | 53 | 3.76 | 1.45 | 30 | 0.0155 | 0.0641 |
| biological_sample_arm_2b.ln_nd1_copies_ul_saliva_10 | Energetics | 0.35 | 8.20 | 2.03 | 61 | 7.48 | 2.07 | 32 | 0.1140 | 0.2593 |
| biological_sample_arm_2b.ln_b2m_copies_ul_saliva_10 | Energetics | 0.40 | 4.58 | 1.93 | 57 | 3.82 | 1.81 | 29 | 0.0756 | 0.1921 |
| biological_sample_arm_2b.ln_nd1_copies_ul_saliva_20 | Energetics | 0.60 | 8.01 | 2.06 | 59 | 6.78 | 1.98 | 32 | 0.0068 | 0.0350 |
| biological_sample_arm_2b.ln_b2m_copies_ul_saliva_20 | Energetics | 0.55 | 4.50 | 2.08 | 54 | 3.37 | 1.88 | 30 | 0.0140 | 0.0589 |
| biological_sample_arm_2b.ln_nd1_copies_ul_saliva_30 | Energetics | 0.46 | 7.71 | 2.20 | 64 | 6.71 | 2.08 | 33 | 0.0308 | 0.1182 |
| biological_sample_arm_2b.ln_b2m_copies_ul_saliva_30 | Energetics | 0.43 | 4.37 | 2.03 | 59 | 3.52 | 1.68 | 27 | 0.0478 | 0.1528 |
| biological_sample_arm_2b.ln_nd1_copies_ul_saliva_60 | Energetics | 0.40 | 7.68 | 2.09 | 63 | 6.87 | 1.91 | 32 | 0.0619 | 0.1700 |
| biological_sample_arm_2b.ln_b2m_copies_ul_saliva_60 | Energetics | 0.42 | 4.29 | 2.02 | 59 | 3.47 | 1.73 | 27 | 0.0578 | 0.1670 |
| biological_sample_arm_2b.ln_nd1_copies_ul_saliva_90 | Energetics | 0.40 | 7.46 | 1.99 | 63 | 6.65 | 2.04 | 32 | 0.0698 | 0.1804 |
| biological_sample_arm_2b.ln_b2m_copies_ul_saliva_90 | Energetics | 0.44 | 4.29 | 1.89 | 54 | 3.45 | 1.87 | 29 | 0.0562 | 0.1670 |
| biological_sample_arm_2b.ln_nd1_copies_ul_saliva_120 | Energetics | 0.56 | 7.21 | 1.99 | 65 | 6.14 | 1.66 | 32 | 0.0067 | 0.0350 |
| biological_sample_arm_2b.ln_b2m_copies_ul_saliva_120 | Energetics | 0.38 | 4.03 | 1.83 | 55 | 3.36 | 1.51 | 23 | 0.0988 | 0.2408 |
| biological_sample_arm_2b.ln_nd1_copies_ul_serum_m5 | Energetics | 0.19 | 7.95 | 1.09 | 65 | 7.73 | 1.21 | 33 | 0.3953 | 0.5755 |
| biological_sample_arm_2b.ln_b2m_copies_ul_serum_m5 | Energetics | 0.26 | 3.33 | 0.79 | 65 | 3.09 | 1.12 | 33 | 0.2747 | 0.4564 |
| biological_sample_arm_2b.ln_nd1_copies_ul_serum_5 | Energetics | 0.19 | 7.92 | 1.05 | 64 | 7.71 | 1.21 | 34 | 0.3835 | 0.5677 |
| biological_sample_arm_2b.ln_b2m_copies_ul_serum_5 | Energetics | 0.17 | 3.39 | 0.79 | 64 | 3.23 | 1.08 | 33 | 0.4693 | 0.6348 |
| biological_sample_arm_2b.ln_nd1_copies_ul_serum_10 | Energetics | 0.02 | 7.91 | 1.09 | 63 | 7.89 | 1.20 | 35 | 0.9266 | 0.9678 |
| biological_sample_arm_2b.ln_b2m_copies_ul_serum_10 | Energetics | 0.27 | 3.30 | 0.78 | 63 | 3.07 | 1.01 | 35 | 0.2424 | 0.4193 |

| Variable | Category | g | M (Controls) | SD (Controls) | N (Controls) | M (Patients) | SD (Patients) | N (Patients) | p | q (FDR) |
|---|---|---|---|---|---|---|---|---|---|---|
| biological_sample_arm_2b.ln_nd1_copies_ul_serum_20 | Energetics | 0.03 | 7.96 | 1.07 | 63 | 7.94 | 1.19 | 34 | 0.9079 | 0.9578 |
| biological_sample_arm_2b.ln_b2m_copies_ul_serum_20 | Energetics | 0.17 | 3.23 | 0.83 | 63 | 3.08 | 1.00 | 34 | 0.4405 | 0.6279 |
| biological_sample_arm_2b.ln_nd1_copies_ul_serum_30 | Energetics | 0.10 | 8.03 | 1.12 | 62 | 7.92 | 1.23 | 33 | 0.6623 | 0.8045 |
| biological_sample_arm_2b.ln_b2m_copies_ul_serum_30 | Energetics | 0.38 | 3.30 | 0.82 | 62 | 2.96 | 0.99 | 33 | 0.0993 | 0.2408 |
| biological_sample_arm_2b.ln_nd1_copies_ul_serum_60 | Energetics | 0.08 | 8.17 | 1.11 | 61 | 8.08 | 1.13 | 31 | 0.7048 | 0.8402 |
| biological_sample_arm_2b.ln_b2m_copies_ul_serum_60 | Energetics | 0.28 | 3.51 | 0.87 | 61 | 3.24 | 1.08 | 31 | 0.2457 | 0.4214 |
| biological_sample_arm_2b.ln_nd1_copies_ul_serum_90 | Energetics | -0.01 | 8.18 | 1.12 | 62 | 8.19 | 1.21 | 30 | 0.9688 | 0.9876 |
| biological_sample_arm_2b.ln_b2m_copies_ul_serum_90 | Energetics | 0.30 | 3.53 | 0.99 | 62 | 3.23 | 0.99 | 30 | 0.1805 | 0.3515 |
| biological_sample_arm_2b.ln_b2m_copies_ul_serum_120 | Energetics | 0.21 | 3.59 | 0.88 | 63 | 3.39 | 1.02 | 31 | 0.3589 | 0.5410 |
| biological_sample_arm_2b.ln_nd1_copies_ul_serum_120 | Energetics | 0.08 | 8.17 | 1.22 | 63 | 8.07 | 1.33 | 31 | 0.7212 | 0.8501 |
| biological_sample_arm_2b.ln_nd1_copies_ul_plasma_m5 | Energetics | 0.09 | 8.35 | 0.83 | 64 | 8.27 | 0.78 | 34 | 0.6490 | 0.8008 |
| biological_sample_arm_2b.ln_b2m_copies_ul_plasma_m5 | Energetics | -0.03 | 1.91 | 0.67 | 62 | 1.93 | 0.63 | 33 | 0.8984 | 0.9578 |
| biological_sample_arm_2b.ln_nd1_copies_ul_plasma_5 | Energetics | 0.07 | 8.28 | 0.86 | 63 | 8.22 | 0.80 | 34 | 0.7181 | 0.8501 |
| biological_sample_arm_2b.ln_b2m_copies_ul_plasma_5 | Energetics | 0.13 | 1.90 | 0.69 | 62 | 1.81 | 0.67 | 32 | 0.5532 | 0.7205 |
| biological_sample_arm_2b.ln_nd1_copies_ul_plasma_10 | Energetics | 0.26 | 8.22 | 0.86 | 62 | 8.00 | 0.79 | 32 | 0.2238 | 0.4037 |
| biological_sample_arm_2b.ln_b2m_copies_ul_plasma_10 | Energetics | 0.11 | 1.89 | 0.71 | 61 | 1.80 | 0.75 | 30 | 0.6117 | 0.7682 |
| biological_sample_arm_2b.ln_nd1_copies_ul_plasma_20 | Energetics | 0.15 | 8.36 | 0.91 | 62 | 8.23 | 0.81 | 32 | 0.4662 | 0.6348 |
| biological_sample_arm_2b.ln_b2m_copies_ul_plasma_20 | Energetics | 0.22 | 2.00 | 0.80 | 61 | 1.83 | 0.82 | 32 | 0.3255 | 0.5163 |
| biological_sample_arm_2b.ln_nd1_copies_ul_plasma_30 | Energetics | 0.25 | 8.15 | 0.86 | 62 | 7.94 | 0.82 | 33 | 0.2417 | 0.4193 |
| biological_sample_arm_2b.ln_b2m_copies_ul_plasma_30 | Energetics | -0.12 | 1.93 | 0.66 | 60 | 2.07 | 1.79 | 31 | 0.6673 | 0.8046 |
| biological_sample_arm_2b.ln_nd1_copies_ul_plasma_60 | Energetics | 0.19 | 8.21 | 0.73 | 61 | 8.06 | 0.87 | 30 | 0.4202 | 0.6032 |
| biological_sample_arm_2b.ln_b2m_copies_ul_plasma_60 | Energetics | -0.01 | 1.88 | 0.64 | 59 | 1.89 | 0.69 | 30 | 0.9619 | 0.9876 |

| Variable | Category | g | M (Controls) | SD (Controls) | N (Controls) | M (Patients) | SD (Patients) | N (Patients) | p | q (FDR) |
|---|---|---|---|---|---|---|---|---|---|---|
| biological_sample_arm_2b.ln_nd1_copies_ul_plasma_90 | Energetics | 0.21 | 8.23 | 0.80 | 62 | 8.06 | 0.87 | 31 | 0.3459 | 0.5333 |
| biological_sample_arm_2b.ln_b2m_copies_ul_plasma_90 | Energetics | 0.43 | 1.97 | 0.64 | 61 | 1.68 | 0.69 | 29 | 0.0677 | 0.1786 |
| biological_sample_arm_2b.ln_nd1_copies_ul_plasma_120 | Energetics | 0.01 | 8.11 | 0.85 | 63 | 8.10 | 0.77 | 31 | 0.9780 | 0.9876 |
| biological_sample_arm_2b.ln_b2m_copies_ul_plasma_120 | Energetics | 0.24 | 2.03 | 0.62 | 61 | 1.87 | 0.74 | 30 | 0.3029 | 0.4916 |
| biological_sample_arm_2b.hgba1c_edta1 | Cellular and Molecular | -0.84 | 5.15 | 0.35 | 68 | 5.65 | 0.89 | 37 | 0.0019 | 0.0111 |
| biological_sample_arm_2b.glucose_serum | Cellular and Molecular | -0.46 | 92.49 | 9.12 | 69 | 106.86 | 52.09 | 36 | 0.1093 | 0.2562 |
| biological_sample_arm_2b.cholesterol_serum | Cellular and Molecular | -0.40 | 178.20 | 40.47 | 69 | 193.83 | 36.82 | 36 | 0.0496 | 0.1561 |
| biological_sample_arm_2b.insulin_serum | Cellular and Molecular | -0.47 | 8.30 | 6.43 | 68 | 13.25 | 15.76 | 32 | 0.0959 | 0.2380 |
| biological_sample_arm_2b.sodium_serum | Cellular and Molecular | -0.04 | 140.12 | 1.84 | 69 | 140.19 | 2.19 | 36 | 0.8546 | 0.9509 |
| biological_sample_arm_2b.potassium_serum | Cellular and Molecular | -0.13 | 4.45 | 0.32 | 69 | 4.51 | 0.67 | 36 | 0.5935 | 0.7590 |
| biological_sample_arm_2b.chloride_serum | Cellular and Molecular | 0.60 | 103.81 | 2.16 | 69 | 102.47 | 2.35 | 36 | 0.0058 | 0.0316 |
| biological_sample_arm_2b.co2_serum | Cellular and Molecular | 0.07 | 25.77 | 2.46 | 69 | 25.58 | 2.59 | 36 | 0.7253 | 0.8502 |
| biological_sample_arm_2b.bun_serum | Cellular and Molecular | -0.69 | 13.12 | 3.94 | 69 | 16.08 | 4.87 | 36 | 0.0025 | 0.0142 |
| biological_sample_arm_2b.creatinine_serum | Cellular and Molecular | 0.32 | 0.83 | 0.15 | 69 | 0.77 | 0.21 | 36 | 0.1638 | 0.3328 |
| biological_sample_arm_2b.albumin_serum | Cellular and Molecular | -0.30 | 4.61 | 0.26 | 69 | 4.70 | 0.36 | 36 | 0.1847 | 0.3542 |
| biological_sample_arm_2b.calcium_serum | Cellular and Molecular | 0.03 | 9.49 | 0.32 | 69 | 9.48 | 0.41 | 36 | 0.8772 | 0.9577 |
| biological_sample_arm_2b.triglycerides_serum | Cellular and Molecular | -0.47 | 87.94 | 66.29 | 69 | 121.86 | 82.69 | 36 | 0.0374 | 0.1272 |
| biological_sample_arm_2b.hdl_serum | Cellular and Molecular | 0.17 | 61.12 | 16.88 | 69 | 58.19 | 16.49 | 36 | 0.3955 | 0.5755 |
| biological_sample_arm_2b.cpeptide_serum | Cellular and Molecular | -0.22 | 1.42 | 0.68 | 69 | 1.59 | 0.99 | 31 | 0.3848 | 0.5677 |
| biological_sample_arm_2b.wbc_edta1 | Immune Regulation and Inflammation | -0.32 | 5.56 | 1.63 | 69 | 6.10 | 1.72 | 37 | 0.1251 | 0.2693 |
| biological_sample_arm_2b.neutroabsolute_edta1 | Immune Regulation and Inflammation | -0.20 | 3.24 | 1.31 | 69 | 3.52 | 1.44 | 37 | 0.3373 | 0.5312 |
| biological_sample_arm_2b.lymphabsolute_edta1 | Immune Regulation and Inflammation | -0.51 | 1.65 | 0.46 | 69 | 1.88 | 0.41 | 37 | 0.0118 | 0.0518 |
| biological_sample_arm_2b.monoabsolute_edta1 | Immune Regulation and Inflammation | -0.11 | 0.46 | 0.12 | 69 | 0.47 | 0.15 | 37 | 0.5976 | 0.7596 |
| biological_sample_arm_2b.eosabsolute_edta1 | Immune Regulation and Inflammation | -0.05 | 0.16 | 0.14 | 69 | 0.16 | 0.12 | 37 | 0.7951 | 0.9101 |
| biological_sample_arm_2b.basoabsolute_edta1 | Immune Regulation and Inflammation | -0.02 | 0.05 | 0.05 | 69 | 0.05 | 0.03 | 37 | 0.9324 | 0.9691 |
| biological_sample_arm_2b.fibrinogen_blue | Immune Regulation and Inflammation | -0.20 | 289.19 | 60.83 | 67 | 301.63 | 67.04 | 35 | 0.3623 | 0.5422 |
| biological_sample_arm_2b.crp_serum | Immune Regulation and Inflammation | -0.11 | 1.96 | 2.82 | 66 | 2.43 | 5.95 | 35 | 0.6596 | 0.8045 |

| Variable | Category | g | M (Controls) | SD (Controls) | N (Controls) | M (Patients) | SD (Patients) | N (Patients) | p | q (FDR) |
|---|---|---|---|---|---|---|---|---|---|---|
| biological_sample_arm_2b.scr_max_sc_5_min | Stress Reactivity Systems | -0.32 | 7.13 | 3.88 | 52 | 8.54 | 5.00 | 32 | 0.1763 | 0.3509 |
| biological_sample_arm_2b.scr_max_sc_10_min | Stress Reactivity Systems | -0.17 | 7.48 | 4.63 | 50 | 8.33 | 5.25 | 30 | 0.4664 | 0.6348 |
| biological_sample_arm_2b.fgf21_fasting_pgml | Cellular and Molecular | -0.87 | 234.99 | 345.29 | 65 | 657.79 | 668.22 | 34 | 0.0013 | 0.0086 |
| biological_sample_arm_2b.fgf21_s1_pgml | Cellular and Molecular | -1.27 | 109.01 | 152.67 | 65 | 567.82 | 580.42 | 33 | 0.0001 | 0.0007 |
| biological_sample_arm_2b.fgf21_s2_pgml | Cellular and Molecular | -1.42 | 103.77 | 151.52 | 64 | 594.72 | 546.91 | 34 | 0.0000 | 0.0001 |
| biological_sample_arm_2b.fgf21_s3_pgml | Cellular and Molecular | -1.34 | 99.48 | 145.73 | 62 | 608.53 | 600.71 | 35 | 0.0000 | 0.0002 |
| biological_sample_arm_2b.fgf21_s4_pgml | Cellular and Molecular | -1.32 | 99.77 | 150.26 | 62 | 565.67 | 553.99 | 34 | 0.0000 | 0.0003 |
| biological_sample_arm_2b.fgf21_s5_pgml | Cellular and Molecular | -1.35 | 98.64 | 151.28 | 62 | 568.87 | 549.60 | 33 | 0.0000 | 0.0003 |
| biological_sample_arm_2b.fgf21_s6_pgml | Cellular and Molecular | -1.34 | 106.02 | 161.76 | 61 | 643.75 | 652.89 | 31 | 0.0001 | 0.0007 |
| biological_sample_arm_2b.fgf21_s7_pgml | Cellular and Molecular | -1.29 | 111.98 | 179.16 | 61 | 682.04 | 730.54 | 29 | 0.0003 | 0.0019 |
| biological_sample_arm_2b.fgf21_s8_pgml | Cellular and Molecular | -1.27 | 105.83 | 185.73 | 63 | 678.43 | 739.01 | 31 | 0.0002 | 0.0014 |
| biological_sample_arm_2b.gdf15_t_fasting1 | Cellular and Molecular | -1.57 | 436.97 | 196.48 | 68 | 1534.62 | 1200.21 | 32 | 0.0000 | 0.0001 |
| biological_sample_arm_2b.gdf15_t_minus5 | Cellular and Molecular | -1.72 | 385.27 | 157.77 | 65 | 1406.12 | 983.11 | 34 | 0.0000 | 0.0000 |
| biological_sample_arm_2b.gdf15_t_5 | Cellular and Molecular | -1.74 | 395.35 | 164.50 | 64 | 1371.98 | 921.34 | 34 | 0.0000 | 0.0000 |
| biological_sample_arm_2b.gdf15_t_10 | Cellular and Molecular | -1.70 | 404.21 | 165.04 | 63 | 1372.57 | 942.93 | 33 | 0.0000 | 0.0000 |
| biological_sample_arm_2b.gdf15_t_20 | Cellular and Molecular | -1.87 | 395.98 | 171.24 | 63 | 1427.93 | 918.28 | 32 | 0.0000 | 0.0000 |
| biological_sample_arm_2b.gdf15_t_30 | Cellular and Molecular | -1.69 | 382.09 | 162.83 | 62 | 1379.35 | 973.31 | 33 | 0.0000 | 0.0000 |
| biological_sample_arm_2b.gdf15_t_60 | Cellular and Molecular | -1.71 | 406.78 | 182.92 | 61 | 1395.11 | 969.21 | 30 | 0.0000 | 0.0001 |
| biological_sample_arm_2b.gdf15_t_90 | Cellular and Molecular | -1.69 | 412.69 | 195.19 | 62 | 1453.23 | 1026.61 | 31 | 0.0000 | 0.0001 |
| biological_sample_arm_2b.gdf15_t_120 | Cellular and Molecular | -1.57 | 415.74 | 210.30 | 63 | 1422.91 | 1071.36 | 31 | 0.0000 | 0.0001 |
| biological_sample_arm_2b.lactate_nmolul__t_m5 | Cellular and Molecular | -1.15 | 0.68 | 0.30 | 50 | 1.15 | 0.57 | 24 | 0.0007 | 0.0046 |
| biological_sample_arm_2b.lactate_nmolul__t_5 | Cellular and Molecular | -1.01 | 0.79 | 0.35 | 50 | 1.25 | 0.61 | 24 | 0.0017 | 0.0104 |
| biological_sample_arm_2b.lactate_nmolul__t_10 | Cellular and Molecular | -1.05 | 0.72 | 0.34 | 50 | 1.22 | 0.65 | 24 | 0.0016 | 0.0099 |
| biological_sample_arm_2b.lactate_nmolul__t_20 | Cellular and Molecular | -1.05 | 0.60 | 0.35 | 50 | 1.11 | 0.67 | 24 | 0.0017 | 0.0104 |
| biological_sample_arm_2b.lactate_nmolul__t_30 | Cellular and Molecular | -0.70 | 0.68 | 0.49 | 50 | 1.06 | 0.62 | 24 | 0.0122 | 0.0524 |
| biological_sample_arm_2b.lactate_nmolul__t_60 | Cellular and Molecular | -0.82 | 0.60 | 0.36 | 50 | 1.01 | 0.69 | 24 | 0.0108 | 0.0496 |
| biological_sample_arm_2b.lactate_nmolul__t_90 | Cellular and Molecular | -0.62 | 0.48 | 0.33 | 50 | 0.77 | 0.67 | 24 | 0.0518 | 0.1606 |
| biological_sample_arm_2b.lactate_nmolul__t_120 | Cellular and Molecular | -0.91 | 0.47 | 0.24 | 50 | 0.86 | 0.66 | 24 | 0.0098 | 0.0464 |

Table S2. Phenome-wide associations with working-memory brain activation in MitoD patients, related to Figure 3.

Age-controlled Spearman partial correlations in patients. Benjamini–Hochberg FDR correction was applied across 304 tests. Variables are sorted by |R|. Sig_FDR: 1 = survives FDR q < 0.05; 0 = does not. N denotes the available sample for each variable.

| Variable | R | p | p_FDR | Sig_FDR | N |
|---|---|---|---|---|---|
| gdf15_t_120 | -0.707 | 0.0007 | 0.0310 | 1 | 20 |
| gdf15_t_10 | -0.697 | 0.0006 | 0.0310 | 1 | 21 |
| gdf15_t_60 | -0.681 | 0.0018 | 0.0431 | 1 | 19 |
| nmdas_i_ii_iii_score | -0.670 | 0.0003 | 0.0310 | 1 | 25 |
| nmdas_i_ii_iii_score_1 | -0.670 | 0.0003 | 0.0310 | 1 | 25 |
| nmdas_i_ii_iii_score_2 | -0.670 | 0.0003 | 0.0310 | 1 | 25 |
| gdf15_t_20 | -0.660 | 0.0021 | 0.0435 | 1 | 20 |
| tm_ls_scaled | 0.646 | 0.0006 | 0.0310 | 1 | 25 |
| nmdas_cca_7 | -0.646 | 0.0006 | 0.0310 | 1 | 25 |
| gdf15_t_30 | -0.644 | 0.0022 | 0.0435 | 1 | 21 |
| fgf21_s7_pgml | -0.640 | 0.0057 | 0.0722 | 0 | 18 |
| gdf15_t_90 | -0.634 | 0.0036 | 0.0573 | 0 | 20 |
| gdf15_t_minus5 | -0.625 | 0.0024 | 0.0435 | 1 | 22 |
| cns_score_stancegait | 0.624 | 0.0011 | 0.0430 | 1 | 25 |
| gdf15_t_5 | -0.616 | 0.0029 | 0.0493 | 1 | 22 |
| dsm_5_disinhibition_imputed | 0.614 | 0.0014 | 0.0431 | 1 | 25 |
| nmdas_cf_10 | -0.613 | 0.0014 | 0.0431 | 1 | 25 |
| triglycerides_serum | -0.610 | 0.0043 | 0.0626 | 0 | 21 |
| cns_score_total | 0.604 | 0.0018 | 0.0431 | 1 | 25 |
| cns_score_total_1 | 0.604 | 0.0018 | 0.0431 | 1 | 25 |
| cns_tndm_wlk_tst | 0.590 | 0.0024 | 0.0435 | 1 | 25 |
| lymphabsolute_edta1 | -0.570 | 0.0070 | 0.0792 | 0 | 22 |
| rbans_cd_scaled | 0.566 | 0.0049 | 0.0643 | 0 | 24 |
| pfs_phys | -0.563 | 0.0042 | 0.0626 | 0 | 25 |
| glucose_serum | -0.563 | 0.0098 | 0.0995 | 0 | 21 |
| lactate_nmolul__t_20 | -0.562 | 0.0188 | 0.1462 | 0 | 18 |
| cns_proximalstr_l | 0.559 | 0.0045 | 0.0628 | 0 | 25 |
| respiratory_rate_rr_breaths_min | -0.556 | 0.1947 | 0.4368 | 0 | 8 |
| gdf15_t_fasting1 | -0.545 | 0.0192 | 0.1462 | 0 | 19 |
| cns_score_musclebulk | 0.544 | 0.0061 | 0.0736 | 0 | 25 |
| fgf21_s8_pgml | -0.542 | 0.0166 | 0.1326 | 0 | 20 |
| cwi_wr_scaled | 0.539 | 0.0065 | 0.0764 | 0 | 25 |
| cpeptide_serum | -0.532 | 0.0230 | 0.1635 | 0 | 19 |

| Variable | R | p | p_FDR | Sig_FDR | N |
|---|---|---|---|---|---|
| tm_nls_scaled | 0.524 | 0.0086 | 0.0938 | 0 | 25 |
| nmdas_cca_5 | -0.516 | 0.0098 | 0.0995 | 0 | 25 |
| lactate_nmolul__t_10 | -0.515 | 0.0345 | 0.1899 | 0 | 18 |
| nab_snl_spd_perc | 0.513 | 0.0123 | 0.1132 | 0 | 24 |
| tm_ns_scaled | 0.511 | 0.0107 | 0.1051 | 0 | 25 |
| fgf21_s4_pgml | -0.503 | 0.0237 | 0.1635 | 0 | 21 |
| cns_kj_reflex_l | 0.503 | 0.0123 | 0.1132 | 0 | 25 |
| nmdas_cf_4 | -0.500 | 0.0129 | 0.1132 | 0 | 25 |
| nmdas_cca_6 | -0.499 | 0.0130 | 0.1132 | 0 | 25 |
| cns_score_cranialnerves | 0.494 | 0.0141 | 0.1189 | 0 | 25 |
| potassium_serum | 0.494 | 0.0269 | 0.1635 | 0 | 21 |
| lactate_nmolul__t_30 | -0.491 | 0.0452 | 0.2035 | 0 | 18 |
| fgf21_s5_pgml | -0.490 | 0.0283 | 0.1646 | 0 | 21 |
| lactate_nmolul__t_5 | -0.489 | 0.0462 | 0.2035 | 0 | 18 |
| cns_kj_reflex_r | 0.488 | 0.0156 | 0.1282 | 0 | 25 |
| insulin_serum | -0.486 | 0.0350 | 0.1899 | 0 | 20 |
| met_ve_d2 | -0.474 | 0.2358 | 0.4910 | 0 | 9 |
| nab_shl_drg_perc | 0.467 | 0.0246 | 0.1635 | 0 | 24 |
| nmdas_ssi_1 | -0.461 | 0.0233 | 0.1635 | 0 | 25 |
| fgf21_s6_pgml | -0.455 | 0.0500 | 0.2056 | 0 | 20 |
| cns_superficial | 0.453 | 0.0261 | 0.1635 | 0 | 25 |
| cns_deep | 0.453 | 0.0261 | 0.1635 | 0 | 25 |
| cns_score_sensation | 0.453 | 0.0261 | 0.1635 | 0 | 25 |
| nmdas_cf_7 | -0.453 | 0.0261 | 0.1635 | 0 | 25 |
| fgf21_s3_pgml | -0.453 | 0.0392 | 0.2035 | 0 | 22 |
| compass_pupillomotor | -0.452 | 0.0265 | 0.1635 | 0 | 25 |
| ln_b2m_copies_ul_saliva_m5 | -0.450 | 0.1229 | 0.3460 | 0 | 14 |
| dcf_sit_stand_time | 0.450 | 0.0275 | 0.1641 | 0 | 25 |
| cwi_inhib_scaled | 0.447 | 0.0287 | 0.1646 | 0 | 25 |
| lactate_nmolul__t_60 | -0.441 | 0.0763 | 0.2668 | 0 | 18 |
| fgf21_s2_pgml | -0.433 | 0.0499 | 0.2056 | 0 | 22 |
| pfs_mental | -0.432 | 0.0348 | 0.1899 | 0 | 25 |
| met_vo2_kg_d2 | -0.431 | 0.2864 | 0.5087 | 0 | 9 |
| cns_gait_statn | 0.430 | 0.0359 | 0.1915 | 0 | 25 |
| fgf21_s1_pgml | -0.422 | 0.0567 | 0.2239 | 0 | 22 |
| nmdas_cf_2 | -0.420 | 0.0409 | 0.2035 | 0 | 25 |
| cns_other | 0.420 | 0.0411 | 0.2035 | 0 | 25 |

| Variable | R | p | p_FDR | Sig_FDR | N |
|---|---|---|---|---|---|
| cns_score_otherfindings | 0.420 | 0.0411 | 0.2035 | 0 | 25 |
| pi_geneticdiagnosistype | -0.419 | 0.0416 | 0.2035 | 0 | 25 |
| cns_height | 0.418 | 0.2627 | 0.4927 | 0 | 10 |
| cns_crnl_nrv_x | 0.413 | 0.0447 | 0.2035 | 0 | 25 |
| cns_hp_tst | 0.413 | 0.0447 | 0.2035 | 0 | 25 |
| cns_distalstrength_l | 0.413 | 0.0447 | 0.2035 | 0 | 25 |
| cns_proximalstr_r | 0.412 | 0.0453 | 0.2035 | 0 | 25 |
| fgf21_fasting_pgml | -0.412 | 0.0798 | 0.2727 | 0 | 20 |
| nmdas_cca_4 | -0.411 | 0.0458 | 0.2035 | 0 | 25 |
| hdl_serum | 0.411 | 0.0720 | 0.2543 | 0 | 21 |
| wbc_edta1 | -0.410 | 0.0648 | 0.2385 | 0 | 22 |
| nmdas_cf_5 | -0.408 | 0.0478 | 0.2056 | 0 | 25 |
| cns_crnl_nrv_vii | 0.404 | 0.0499 | 0.2056 | 0 | 25 |
| cns_crnl_nrv_ix | 0.404 | 0.0499 | 0.2056 | 0 | 25 |
| nmdas_cf_9 | -0.403 | 0.0508 | 0.2061 | 0 | 25 |
| mfis_phy_sub | -0.394 | 0.0564 | 0.2239 | 0 | 25 |
| hr_avg_p5_60s_2 | -0.394 | 0.0696 | 0.2487 | 0 | 23 |
| karnofsky_100 | 0.393 | 0.3836 | 0.6011 | 0 | 8 |
| neutroabsolute_edta1 | -0.391 | 0.0794 | 0.2727 | 0 | 22 |
| lactate_nmolul__t_120 | -0.391 | 0.1211 | 0.3460 | 0 | 18 |
| nmdas_cca_2 | -0.390 | 0.0594 | 0.2311 | 0 | 25 |
| rbans_ll_scaled | 0.389 | 0.0601 | 0.2311 | 0 | 25 |
| cwi_cn_scaled | 0.387 | 0.0621 | 0.2359 | 0 | 25 |
| cwi_inh_cn_scaled | -0.383 | 0.0650 | 0.2385 | 0 | 25 |
| compass_secretomotor | -0.382 | 0.0651 | 0.2385 | 0 | 25 |
| ipaq_intensity | 0.377 | 0.0693 | 0.2487 | 0 | 25 |
| hgba1c_edta1 | -0.376 | 0.0930 | 0.3073 | 0 | 22 |
| leq_positive_events_score | 0.371 | 0.1177 | 0.3440 | 0 | 20 |
| crp_serum | -0.362 | 0.1165 | 0.3438 | 0 | 21 |
| stai_y1_state_total | -0.362 | 0.0826 | 0.2791 | 0 | 25 |
| nmdas_cca_3 | -0.358 | 0.0857 | 0.2863 | 0 | 25 |
| adj_wgt_perc | -0.343 | 0.1007 | 0.3257 | 0 | 25 |
| cns_reg_wlk_tst | 0.343 | 0.1007 | 0.3257 | 0 | 25 |
| nab_sdgf_perc | 0.343 | 0.1091 | 0.3418 | 0 | 24 |
| compass_total | -0.341 | 0.1027 | 0.3287 | 0 | 25 |
| hr_avg_p5_60s_1 | -0.341 | 0.1111 | 0.3433 | 0 | 24 |
| cns_score_ocularfundi | 0.336 | 0.1085 | 0.3418 | 0 | 25 |

| Variable | R | p | p_FDR | Sig_FDR | N |
|---|---|---|---|---|---|
| eosabsolute_edta1 | 0.335 | 0.1377 | 0.3703 | 0 | 22 |
| neo_agree | 0.331 | 0.1146 | 0.3433 | 0 | 25 |
| cns_score_weight | 0.330 | 0.1152 | 0.3433 | 0 | 25 |
| cns_score_height | 0.330 | 0.1152 | 0.3433 | 0 | 25 |
| hgt_perc | -0.330 | 0.1152 | 0.3433 | 0 | 25 |
| met_tidal_d2 | 0.330 | 0.4247 | 0.6392 | 0 | 9 |
| ln_b2m_copies_ul_saliva_5 | -0.329 | 0.1968 | 0.4368 | 0 | 18 |
| albumin_serum | 0.328 | 0.1584 | 0.3852 | 0 | 21 |
| cns_score_cerebellarfunction | 0.324 | 0.1228 | 0.3460 | 0 | 25 |
| cns_crnl_nrv_iv | 0.323 | 0.1240 | 0.3460 | 0 | 25 |
| cns_crnl_nrv_vi | 0.323 | 0.1240 | 0.3460 | 0 | 25 |
| nmdas_cca_1 | -0.319 | 0.1292 | 0.3572 | 0 | 25 |
| nmdas_cf_3 | -0.317 | 0.1309 | 0.3585 | 0 | 25 |
| cns_distalstrength_r | 0.314 | 0.1354 | 0.3675 | 0 | 25 |
| vf_c1_cs_cor_scaled | 0.311 | 0.1491 | 0.3819 | 0 | 24 |
| cns_distalbulk_r | 0.305 | 0.1479 | 0.3819 | 0 | 25 |
| cns_distalbulk_l | 0.305 | 0.1479 | 0.3819 | 0 | 25 |
| cns_score_myotaticreflexes | 0.305 | 0.1479 | 0.3819 | 0 | 25 |
| cns_peripheral_rtn | 0.304 | 0.1485 | 0.3819 | 0 | 25 |
| mfis_psy_sub | -0.303 | 0.1495 | 0.3819 | 0 | 25 |
| vf_c1_cf_cor_scaled | 0.302 | 0.1509 | 0.3822 | 0 | 25 |
| ln_nd1_copies_ul_saliva_120 | -0.302 | 0.2090 | 0.4539 | 0 | 20 |
| nmdas_cf_1 | -0.301 | 0.1529 | 0.3837 | 0 | 25 |
| mbi_cyn | -0.301 | 0.1635 | 0.3944 | 0 | 24 |
| cns_speech_coord | 0.300 | 0.1545 | 0.3837 | 0 | 25 |
| vf_c1_lf_cor_scaled | 0.299 | 0.1552 | 0.3837 | 0 | 25 |
| bdi_total | -0.297 | 0.1583 | 0.3852 | 0 | 25 |
| calcium_serum | 0.294 | 0.2091 | 0.4539 | 0 | 21 |
| compass_vasomotor | -0.291 | 0.1671 | 0.4001 | 0 | 25 |
| nmdas_cf_6 | -0.290 | 0.1693 | 0.4022 | 0 | 25 |
| met_feo2_d2 | -0.285 | 0.4946 | 0.7059 | 0 | 9 |
| cns_wj_reflex_l | 0.284 | 0.1791 | 0.4187 | 0 | 25 |
| cns_wj_reflex_r | 0.284 | 0.1791 | 0.4187 | 0 | 25 |
| vf_c1_cs_tsa_scaled | 0.282 | 0.1928 | 0.4368 | 0 | 24 |
| cns_heel_wlk_tst | 0.281 | 0.1838 | 0.4266 | 0 | 25 |
| compass_gastrointest | -0.278 | 0.1878 | 0.4326 | 0 | 25 |
| ln_nd1_copies_ul_saliva_10 | -0.278 | 0.2494 | 0.4927 | 0 | 20 |

| Variable | R | p | p_FDR | Sig_FDR | N |
|---|---|---|---|---|---|
| cns_crnl_nrv_iii | 0.275 | 0.1932 | 0.4368 | 0 | 25 |
| ln_b2m_copies_ul_saliva_120 | -0.274 | 0.3654 | 0.5796 | 0 | 14 |
| mfis_cog_sub | -0.274 | 0.1957 | 0.4368 | 0 | 25 |
| nmdas_ssi_2 | -0.267 | 0.2074 | 0.4539 | 0 | 25 |
| ln_b2m_copies_ul_plasma_90 | -0.265 | 0.3034 | 0.5270 | 0 | 18 |
| ln_b2m_copies_ul_plasma_m5 | 0.265 | 0.2595 | 0.4927 | 0 | 21 |
| srh_score | 0.265 | 0.2115 | 0.4561 | 0 | 25 |
| nab_shl_irg_perc | 0.258 | 0.2351 | 0.4910 | 0 | 24 |
| neo_extra | 0.257 | 0.2253 | 0.4824 | 0 | 25 |
| hr_avg_p10_60s_2 | -0.256 | 0.2378 | 0.4918 | 0 | 24 |
| ln_b2m_copies_ul_plasma_5 | 0.256 | 0.2900 | 0.5095 | 0 | 20 |
| scor_abp1_psqi_global | -0.255 | 0.2289 | 0.4846 | 0 | 25 |
| nmdas_ssi_6 | -0.255 | 0.2296 | 0.4846 | 0 | 25 |
| lactate_nmolul__t_90 | -0.245 | 0.3426 | 0.5600 | 0 | 18 |
| hr_avg_p5_300s | -0.245 | 0.2593 | 0.4927 | 0 | 24 |
| respr_count_p10_60s_2 | -0.245 | 0.2979 | 0.5204 | 0 | 21 |
| respr_count_p5_60s_2 | -0.242 | 0.3181 | 0.5388 | 0 | 20 |
| ln_b2m_copies_ul_serum_30 | -0.241 | 0.3062 | 0.5289 | 0 | 21 |
| nmdas_ssi_5 | -0.241 | 0.2574 | 0.4927 | 0 | 25 |
| nmdas_ssi_9 | -0.240 | 0.2586 | 0.4927 | 0 | 25 |
| dsm_5_antagonism_imputed | 0.236 | 0.2660 | 0.4927 | 0 | 25 |
| cns_crnl_nrv_xi | 0.236 | 0.2665 | 0.4927 | 0 | 25 |
| met_ree_d2 | -0.236 | 0.5738 | 0.7640 | 0 | 9 |
| nmdas_ssi_3 | -0.236 | 0.2674 | 0.4927 | 0 | 25 |
| nmdas_cca_8 | -0.236 | 0.2674 | 0.4927 | 0 | 25 |
| cns_nasopharynx | 0.236 | 0.2674 | 0.4927 | 0 | 25 |
| cns_crnl_nrv_v | 0.236 | 0.2674 | 0.4927 | 0 | 25 |
| cns_crnl_nrv_viii | 0.236 | 0.2674 | 0.4927 | 0 | 25 |
| cns_toe_wlk_tst | 0.236 | 0.2674 | 0.4927 | 0 | 25 |
| cns_romberg_tst | 0.236 | 0.2674 | 0.4927 | 0 | 25 |
| cns_fng_ns_fng | 0.236 | 0.2674 | 0.4927 | 0 | 25 |
| cns_left_toe | 0.236 | 0.2674 | 0.4927 | 0 | 25 |
| cns_score_toesign | 0.236 | 0.2674 | 0.4927 | 0 | 25 |
| ipaq_total_met | 0.232 | 0.2878 | 0.5087 | 0 | 24 |
| cns_tj_reflex_l | 0.230 | 0.2794 | 0.5020 | 0 | 25 |
| cns_tj_reflex_r | 0.230 | 0.2794 | 0.5020 | 0 | 25 |
| cns_proximalbulk_r | 0.229 | 0.2807 | 0.5020 | 0 | 25 |

| Variable | R | p | p_FDR | Sig_FDR | N |
|---|---|---|---|---|---|
| cns_proximalbulk_l | 0.229 | 0.2807 | 0.5020 | 0 | 25 |
| nmdas_cf_8 | -0.229 | 0.2807 | 0.5020 | 0 | 25 |
| hr_avg_p10_300s | -0.220 | 0.3120 | 0.5354 | 0 | 24 |
| met_vo2_d2 | -0.220 | 0.6012 | 0.7849 | 0 | 9 |
| ln_b2m_copies_ul_serum_10 | -0.216 | 0.3476 | 0.5621 | 0 | 22 |
| dcg_adj_wgt | 0.215 | 0.3135 | 0.5354 | 0 | 25 |
| ctq_pa | 0.212 | 0.3191 | 0.5388 | 0 | 25 |
| general_medical_and_neurological_examination_complete | -0.211 | 0.3223 | 0.5397 | 0 | 25 |
| dsm_5_neg_aff_imputed | 0.211 | 0.3231 | 0.5397 | 0 | 25 |
| fvc_average | 0.209 | 0.3392 | 0.5600 | 0 | 24 |
| hr_avg_p10_60s_1 | -0.208 | 0.3409 | 0.5600 | 0 | 24 |
| ln_nd1_copies_ul_saliva_30 | -0.208 | 0.3661 | 0.5796 | 0 | 22 |
| stai_y1_trait_total | -0.208 | 0.3306 | 0.5492 | 0 | 25 |
| lactate_nmolul__t_m5 | -0.205 | 0.4310 | 0.6392 | 0 | 18 |
| cns_score_generalmedical | 0.201 | 0.3451 | 0.5610 | 0 | 25 |
| wasi_full_sum_c | 0.199 | 0.3512 | 0.5646 | 0 | 25 |
| l_cat | 0.198 | 0.3529 | 0.5646 | 0 | 25 |
| ln_b2m_copies_ul_saliva_10 | -0.194 | 0.4564 | 0.6671 | 0 | 18 |
| mbi_ineff | 0.192 | 0.3794 | 0.5975 | 0 | 24 |
| fev1_average | 0.188 | 0.3912 | 0.6082 | 0 | 24 |
| topf_standrd_score | 0.187 | 0.3921 | 0.6082 | 0 | 24 |
| creatinine_serum | 0.187 | 0.4298 | 0.6392 | 0 | 21 |
| leq_negative_events_score | -0.186 | 0.4081 | 0.6238 | 0 | 23 |
| scr_max_sc_5_min | 0.185 | 0.4227 | 0.6392 | 0 | 22 |
| ln_nd1_copies_ul_plasma_120 | -0.184 | 0.4501 | 0.6610 | 0 | 20 |
| neo_cons | -0.179 | 0.4021 | 0.6205 | 0 | 25 |
| cns_truncal_coord | 0.177 | 0.4084 | 0.6238 | 0 | 25 |
| cns_macula_regn | 0.175 | 0.4123 | 0.6267 | 0 | 25 |
| cns_eye_mvt | 0.170 | 0.4274 | 0.6392 | 0 | 25 |
| cns_crnl_nrv_ii | 0.168 | 0.4335 | 0.6397 | 0 | 25 |
| fibrinogen_blue | -0.158 | 0.5055 | 0.7114 | 0 | 21 |
| dcf_bodyfat_avg | -0.158 | 0.4610 | 0.6705 | 0 | 25 |
| compass_orth_int | -0.156 | 0.4656 | 0.6740 | 0 | 25 |
| ln_nd1_copies_ul_saliva_5 | -0.156 | 0.5106 | 0.7139 | 0 | 21 |
| ln_nd1_copies_ul_serum_m5 | -0.156 | 0.5004 | 0.7079 | 0 | 22 |
| cns_alt_supin_pron | 0.154 | 0.4724 | 0.6807 | 0 | 25 |
| nmdas_ssi_7 | -0.150 | 0.4844 | 0.6947 | 0 | 25 |

| Variable | R | p | p_FDR | Sig_FDR | N |
|---|---|---|---|---|---|
| ln_b2m_copies_ul_serum_120 | 0.146 | 0.5498 | 0.7429 | 0 | 20 |
| cns_aj_reflex_l | 0.144 | 0.5007 | 0.7079 | 0 | 25 |
| ln_b2m_copies_ul_serum_5 | -0.143 | 0.5464 | 0.7416 | 0 | 21 |
| cns_skeletal | 0.141 | 0.5119 | 0.7139 | 0 | 25 |
| hip_waist_circum_ratio | -0.139 | 0.5167 | 0.7172 | 0 | 25 |
| fvc_predicted_average | 0.135 | 0.5389 | 0.7346 | 0 | 24 |
| tics_total_score | 0.134 | 0.5316 | 0.7346 | 0 | 25 |
| mcc_score | -0.133 | 0.5347 | 0.7346 | 0 | 25 |
| cns_retinal_vssls | -0.132 | 0.5386 | 0.7346 | 0 | 25 |
| ln_nd1_copies_ul_plasma_m5 | -0.131 | 0.5715 | 0.7640 | 0 | 22 |
| ln_b2m_copies_ul_plasma_10 | 0.123 | 0.6146 | 0.7964 | 0 | 20 |
| ln_nd1_copies_ul_plasma_5 | 0.121 | 0.6010 | 0.7849 | 0 | 22 |
| neo_neuro | -0.121 | 0.5738 | 0.7640 | 0 | 25 |
| cns_aj_reflex_r | 0.120 | 0.5755 | 0.7640 | 0 | 25 |
| respr_count_p10_60s_1 | -0.119 | 0.6269 | 0.8008 | 0 | 20 |
| cholesterol_serum | 0.118 | 0.6204 | 0.7964 | 0 | 21 |
| cns_eyes | 0.118 | 0.5832 | 0.7708 | 0 | 25 |
| ln_b2m_copies_ul_serum_20 | -0.118 | 0.6209 | 0.7964 | 0 | 21 |
| ln_b2m_copies_ul_plasma_20 | 0.117 | 0.6338 | 0.8040 | 0 | 20 |
| dsm_5_psychoticism | -0.115 | 0.6016 | 0.7849 | 0 | 24 |
| ln_nd1_copies_ul_saliva_m5 | -0.110 | 0.6651 | 0.8248 | 0 | 19 |
| ln_nd1_copies_ul_serum_20 | -0.110 | 0.6457 | 0.8145 | 0 | 21 |
| met_ree_d1 | 0.108 | 0.7991 | 0.8932 | 0 | 9 |
| dsm_5_detachment_imputed | -0.107 | 0.6183 | 0.7964 | 0 | 25 |
| ln_nd1_copies_ul_serum_120 | 0.105 | 0.6674 | 0.8248 | 0 | 20 |
| cns_ocular_fundi | 0.102 | 0.6348 | 0.8040 | 0 | 25 |
| ln_nd1_copies_ul_plasma_10 | -0.099 | 0.6788 | 0.8354 | 0 | 21 |
| cns_hc | 0.099 | 0.6627 | 0.8248 | 0 | 23 |
| ln_nd1_copies_ul_serum_90 | 0.098 | 0.6993 | 0.8537 | 0 | 19 |
| ssq_n | 0.097 | 0.6505 | 0.8171 | 0 | 25 |
| cns_abdomen | 0.096 | 0.6566 | 0.8214 | 0 | 25 |
| ln_b2m_copies_ul_serum_m5 | -0.093 | 0.6875 | 0.8427 | 0 | 22 |
| ln_nd1_copies_ul_saliva_20 | 0.092 | 0.7080 | 0.8574 | 0 | 20 |
| ln_b2m_copies_ul_plasma_30 | -0.092 | 0.7169 | 0.8600 | 0 | 19 |
| ln_b2m_copies_ul_saliva_90 | -0.090 | 0.7300 | 0.8600 | 0 | 18 |
| ln_nd1_copies_ul_plasma_90 | -0.089 | 0.7168 | 0.8600 | 0 | 20 |
| ln_nd1_copies_ul_serum_60 | -0.085 | 0.7305 | 0.8600 | 0 | 20 |

| Variable | R | p | p_FDR | Sig_FDR | N |
|---|---|---|---|---|---|
| dsm_5_cross_cutting_imputed | -0.082 | 0.7034 | 0.8553 | 0 | 25 |
| scr_max_sc_10_min | 0.081 | 0.7413 | 0.8600 | 0 | 20 |
| ctq_pn | 0.077 | 0.7193 | 0.8600 | 0 | 25 |
| ln_nd1_copies_ul_saliva_60 | 0.077 | 0.7553 | 0.8600 | 0 | 20 |
| dhs_score | 0.075 | 0.7259 | 0.8600 | 0 | 25 |
| monoabsolute_edta1 | -0.074 | 0.7499 | 0.8600 | 0 | 22 |
| ctq_ea | 0.073 | 0.7352 | 0.8600 | 0 | 25 |
| pss_total | -0.073 | 0.7357 | 0.8600 | 0 | 25 |
| fev1fvc_average | -0.072 | 0.7454 | 0.8600 | 0 | 24 |
| compass_bladded | 0.071 | 0.7410 | 0.8600 | 0 | 25 |
| ln_b2m_copies_ul_saliva_30 | -0.070 | 0.8049 | 0.8963 | 0 | 16 |
| nab_sdgb_perc | -0.070 | 0.7521 | 0.8600 | 0 | 24 |
| mbi_exhaus | -0.069 | 0.7528 | 0.8600 | 0 | 24 |
| cns_weight | 0.069 | 0.8607 | 0.9269 | 0 | 10 |
| ln_nd1_copies_ul_serum_5 | -0.068 | 0.7704 | 0.8707 | 0 | 22 |
| pcl_total | 0.067 | 0.7541 | 0.8600 | 0 | 25 |
| nmdas_complete | -0.065 | 0.7644 | 0.8671 | 0 | 25 |
| ln_b2m_copies_ul_serum_90 | -0.065 | 0.7992 | 0.8932 | 0 | 19 |
| st_free_corsort_scaled | -0.061 | 0.7979 | 0.8932 | 0 | 21 |
| respr_count_p5_60s_1 | 0.057 | 0.8211 | 0.9110 | 0 | 19 |
| basoabsolute_edta1 | -0.048 | 0.8359 | 0.9198 | 0 | 22 |
| fev1_predicted_average | 0.048 | 0.8292 | 0.9167 | 0 | 24 |
| ln_b2m_copies_ul_saliva_20 | 0.047 | 0.8542 | 0.9269 | 0 | 19 |
| ln_b2m_copies_ul_plasma_120 | -0.045 | 0.8605 | 0.9269 | 0 | 19 |
| fat_free_mass | 0.044 | 0.8381 | 0.9198 | 0 | 25 |
| chloride_serum | 0.043 | 0.8572 | 0.9269 | 0 | 21 |
| neo_open | 0.039 | 0.8561 | 0.9269 | 0 | 25 |
| ln_nd1_copies_ul_serum_30 | 0.039 | 0.8719 | 0.9333 | 0 | 21 |
| ctq_en | 0.037 | 0.8629 | 0.9269 | 0 | 25 |
| ln_nd1_copies_ul_plasma_20 | -0.036 | 0.8849 | 0.9374 | 0 | 20 |
| ln_nd1_copies_ul_plasma_30 | 0.033 | 0.8900 | 0.9394 | 0 | 21 |
| ctq_sa | 0.032 | 0.8834 | 0.9374 | 0 | 25 |
| cns_skin | -0.031 | 0.8850 | 0.9374 | 0 | 25 |
| ssq_s | 0.030 | 0.8935 | 0.9398 | 0 | 23 |
| ln_nd1_copies_ul_saliva_90 | -0.027 | 0.9119 | 0.9510 | 0 | 20 |
| ln_b2m_copies_ul_saliva_60 | -0.027 | 0.9150 | 0.9510 | 0 | 19 |
| ln_b2m_copies_ul_serum_60 | -0.026 | 0.9173 | 0.9510 | 0 | 20 |

| Variable | R | p | p_FDR | Sig_FDR | N |
|---|---|---|---|---|---|
| nmdas_ssi_8 | 0.025 | 0.9068 | 0.9506 | 0 | 25 |
| ln_nd1_copies_ul_serum_10 | 0.023 | 0.9221 | 0.9510 | 0 | 22 |
| cns_ears | -0.021 | 0.9228 | 0.9510 | 0 | 25 |
| bun_serum | 0.018 | 0.9394 | 0.9648 | 0 | 21 |
| ctq_md | 0.015 | 0.9461 | 0.9684 | 0 | 25 |
| ln_nd1_copies_ul_plasma_60 | 0.014 | 0.9551 | 0.9743 | 0 | 19 |
| ln_b2m_copies_ul_plasma_60 | 0.010 | 0.9696 | 0.9843 | 0 | 19 |
| co2_serum | -0.009 | 0.9713 | 0.9843 | 0 | 21 |
| leq_total_events_score | 0.006 | 0.9761 | 0.9858 | 0 | 25 |
| sodium_serum | 0.003 | 0.9891 | 0.9924 | 0 | 21 |
| cns_bj_reflex_l | -0.002 | 0.9915 | 0.9924 | 0 | 25 |
| cns_bj_reflex_r | -0.002 | 0.9924 | 0.9924 | 0 | 25 |

Table S3. Phenome-wide associations with working-memory behavioral performance in MitoD patients, related to Figure 3.

Age-controlled Spearman partial correlations in patients. Benjamini–Hochberg FDR correction was applied across 304 tests. Variables are sorted by |R|. Sig_FDR: 1 = survives FDR q < 0.05; 0 = does not. N denotes the available sample for each variable.

| Variable | R | p | p_FDR | Sig_FDR | N |
|---|---|---|---|---|---|
| respiratory_rate_rr_breaths_min | -0.724 | 0.0656 | 0.3067 | 0 | 8 |
| tm_nls_scaled | 0.665 | 0.0004 | 0.0948 | 0 | 25 |
| wbc_edta1 | -0.662 | 0.0011 | 0.0948 | 0 | 22 |
| neutroabsolute_edta1 | -0.653 | 0.0013 | 0.0948 | 0 | 22 |
| met_vo2_kg_d2 | -0.633 | 0.0924 | 0.3546 | 0 | 9 |
| cns_score_musclebulk | 0.618 | 0.0013 | 0.0948 | 0 | 25 |
| hdl_serum | 0.614 | 0.0040 | 0.0948 | 0 | 21 |
| gdf15_t_120 | -0.605 | 0.0061 | 0.1228 | 0 | 20 |
| cns_proximalstr_l | 0.600 | 0.0019 | 0.0948 | 0 | 25 |
| tm_ls_scaled | 0.583 | 0.0028 | 0.0948 | 0 | 25 |
| wasi_full_sum_c | 0.582 | 0.0029 | 0.0948 | 0 | 25 |
| nmdas_cca_4 | -0.573 | 0.0034 | 0.0948 | 0 | 25 |
| cns_height | 0.573 | 0.1068 | 0.3591 | 0 | 10 |
| cns_crnl_nrv_x | 0.571 | 0.0036 | 0.0948 | 0 | 25 |
| cns_hp_tst | 0.571 | 0.0036 | 0.0948 | 0 | 25 |
| cns_distalstrength_l | 0.571 | 0.0036 | 0.0948 | 0 | 25 |
| gdf15_t_60 | -0.570 | 0.0135 | 0.1866 | 0 | 19 |
| gdf15_t_10 | -0.569 | 0.0088 | 0.1576 | 0 | 21 |
| cns_score_stancegait | 0.564 | 0.0041 | 0.0948 | 0 | 25 |
| nmdas_cca_7 | -0.561 | 0.0044 | 0.0948 | 0 | 25 |
| fgf21_s7_pgml | -0.550 | 0.0222 | 0.1885 | 0 | 18 |
| cns_proximalstr_r | 0.528 | 0.0080 | 0.1527 | 0 | 25 |
| gdf15_t_minus5 | -0.519 | 0.0159 | 0.1885 | 0 | 22 |
| cns_distalstrength_r | 0.514 | 0.0102 | 0.1715 | 0 | 25 |
| triglycerides_serum | -0.513 | 0.0207 | 0.1885 | 0 | 21 |
| gdf15_t_5 | -0.510 | 0.0182 | 0.1885 | 0 | 22 |
| dsm_5_antagonism_imputed | 0.509 | 0.0112 | 0.1786 | 0 | 25 |
| ln_b2m_copies_ul_serum_30 | -0.505 | 0.0232 | 0.1885 | 0 | 21 |
| gdf15_t_30 | -0.504 | 0.0234 | 0.1885 | 0 | 21 |
| cwi_inh_cn_scaled | -0.502 | 0.0125 | 0.1841 | 0 | 25 |
| gdf15_t_20 | -0.501 | 0.0288 | 0.2035 | 0 | 20 |
| ctq_pa | 0.501 | 0.0127 | 0.1841 | 0 | 25 |
| fev1_average | 0.495 | 0.0164 | 0.1885 | 0 | 24 |

| Variable | R | p | p_FDR | Sig_FDR | N |
|---|---|---|---|---|---|
| cholesterol_serum | 0.485 | 0.0301 | 0.2062 | 0 | 21 |
| lactate_nmolul__t_20 | -0.474 | 0.0548 | 0.2830 | 0 | 18 |
| cns_crnl_nrv_vii | 0.470 | 0.0204 | 0.1885 | 0 | 25 |
| cns_crnl_nrv_ix | 0.470 | 0.0204 | 0.1885 | 0 | 25 |
| met_ve_d2 | -0.467 | 0.2434 | 0.5081 | 0 | 9 |
| nmdas_i_ii_iii_score | -0.460 | 0.0238 | 0.1885 | 0 | 25 |
| nmdas_i_ii_iii_score_1 | -0.460 | 0.0238 | 0.1885 | 0 | 25 |
| nmdas_i_ii_iii_score_2 | -0.460 | 0.0238 | 0.1885 | 0 | 25 |
| dcg_adj_wgt | 0.459 | 0.0240 | 0.1885 | 0 | 25 |
| cns_tndm_wlk_tst | 0.459 | 0.0242 | 0.1885 | 0 | 25 |
| nmdas_cca_5 | -0.457 | 0.0248 | 0.1885 | 0 | 25 |
| compass_secretomotor | -0.457 | 0.0248 | 0.1885 | 0 | 25 |
| cns_score_total | 0.457 | 0.0248 | 0.1885 | 0 | 25 |
| cns_score_total_1 | 0.457 | 0.0248 | 0.1885 | 0 | 25 |
| nmdas_cf_10 | -0.454 | 0.0260 | 0.1928 | 0 | 25 |
| gdf15_t_90 | -0.452 | 0.0517 | 0.2830 | 0 | 20 |
| nmdas_cca_1 | -0.449 | 0.0278 | 0.2011 | 0 | 25 |
| nmdas_cf_3 | -0.442 | 0.0305 | 0.2062 | 0 | 25 |
| met_ree_d1 | 0.435 | 0.2816 | 0.5189 | 0 | 9 |
| vf_c1_cs_cor_scaled | 0.433 | 0.0389 | 0.2517 | 0 | 24 |
| vf_c1_lf_cor_scaled | 0.431 | 0.0355 | 0.2348 | 0 | 25 |
| nab_shl_drg_perc | 0.429 | 0.0411 | 0.2603 | 0 | 24 |
| respr_count_p5_60s_2 | -0.428 | 0.0672 | 0.3097 | 0 | 20 |
| lactate_nmolul__t_90 | -0.424 | 0.0897 | 0.3546 | 0 | 18 |
| ln_b2m_copies_ul_serum_5 | -0.423 | 0.0630 | 0.2994 | 0 | 21 |
| monoabsolute_edta1 | -0.422 | 0.0568 | 0.2830 | 0 | 22 |
| cns_distalbulk_r | 0.414 | 0.0440 | 0.2676 | 0 | 25 |
| cns_distalbulk_l | 0.414 | 0.0440 | 0.2676 | 0 | 25 |
| nmdas_cf_4 | -0.413 | 0.0449 | 0.2676 | 0 | 25 |
| cns_speech_coord | 0.410 | 0.0464 | 0.2712 | 0 | 25 |
| tm_ns_scaled | 0.408 | 0.0477 | 0.2736 | 0 | 25 |
| glucose_serum | -0.408 | 0.0743 | 0.3323 | 0 | 21 |
| met_tidal_d2 | 0.404 | 0.3214 | 0.5629 | 0 | 9 |
| fat_free_mass | 0.400 | 0.0529 | 0.2830 | 0 | 25 |
| hgt_perc | -0.396 | 0.0552 | 0.2830 | 0 | 25 |
| cns_score_weight | 0.396 | 0.0552 | 0.2830 | 0 | 25 |
| cns_score_height | 0.396 | 0.0552 | 0.2830 | 0 | 25 |

| Variable | R | p | p_FDR | Sig_FDR | N |
|---|---|---|---|---|---|
| ctq_ea | 0.395 | 0.0559 | 0.2830 | 0 | 25 |
| cns_score_cranialnerves | 0.392 | 0.0581 | 0.2848 | 0 | 25 |
| gdf15_t_fasting1 | -0.390 | 0.1093 | 0.3591 | 0 | 19 |
| cwi_wr_scaled | 0.386 | 0.0624 | 0.2994 | 0 | 25 |
| fgf21_s8_pgml | -0.385 | 0.1035 | 0.3591 | 0 | 20 |
| fvc_average | 0.383 | 0.0710 | 0.3223 | 0 | 24 |
| ln_b2m_copies_ul_serum_10 | -0.378 | 0.0912 | 0.3546 | 0 | 22 |
| lactate_nmolul__t_30 | -0.373 | 0.1405 | 0.3931 | 0 | 18 |
| cns_hc | 0.370 | 0.0899 | 0.3546 | 0 | 23 |
| cns_crnl_nrv_xi | 0.370 | 0.0754 | 0.3323 | 0 | 25 |
| fgf21_s4_pgml | -0.369 | 0.1099 | 0.3591 | 0 | 21 |
| topf_standrd_score | 0.368 | 0.0844 | 0.3546 | 0 | 24 |
| lymphabsolute_edta1 | -0.367 | 0.1015 | 0.3591 | 0 | 22 |
| ln_nd1_copies_ul_serum_m5 | -0.365 | 0.1034 | 0.3591 | 0 | 22 |
| fgf21_s5_pgml | -0.365 | 0.1136 | 0.3674 | 0 | 21 |
| vf_c1_cs_tsa_scaled | 0.360 | 0.0914 | 0.3546 | 0 | 24 |
| cwi_cn_scaled | 0.359 | 0.0851 | 0.3546 | 0 | 25 |
| rbans_cd_scaled | 0.358 | 0.0933 | 0.3546 | 0 | 24 |
| ln_b2m_copies_ul_saliva_5 | -0.356 | 0.1612 | 0.4188 | 0 | 18 |
| cns_retinal_vssls | -0.355 | 0.0892 | 0.3546 | 0 | 25 |
| cpeptide_serum | -0.353 | 0.1508 | 0.4130 | 0 | 19 |
| fev1_predicted_average | 0.353 | 0.0989 | 0.3591 | 0 | 24 |
| neo_extra | 0.352 | 0.0918 | 0.3546 | 0 | 25 |
| nmdas_cca_6 | -0.349 | 0.0941 | 0.3546 | 0 | 25 |
| ctq_en | 0.349 | 0.0945 | 0.3546 | 0 | 25 |
| basoabsolute_edta1 | 0.347 | 0.1237 | 0.3836 | 0 | 22 |
| nmdas_ssi_1 | -0.346 | 0.0978 | 0.3591 | 0 | 25 |
| fev1fvc_average | 0.345 | 0.1068 | 0.3591 | 0 | 24 |
| dsm_5_disinhibition_imputed | 0.342 | 0.1016 | 0.3591 | 0 | 25 |
| ln_b2m_copies_ul_serum_m5 | -0.341 | 0.1307 | 0.3931 | 0 | 22 |
| ln_b2m_copies_ul_serum_20 | -0.339 | 0.1438 | 0.3974 | 0 | 21 |
| leq_positive_events_score | 0.338 | 0.1565 | 0.4188 | 0 | 20 |
| dsm_5_neg_aff_imputed | 0.336 | 0.1080 | 0.3591 | 0 | 25 |
| l_cat | 0.335 | 0.1093 | 0.3591 | 0 | 25 |
| ln_nd1_copies_ul_serum_60 | -0.335 | 0.1608 | 0.4188 | 0 | 20 |
| fgf21_s6_pgml | -0.332 | 0.1651 | 0.4218 | 0 | 20 |
| cns_weight | 0.330 | 0.3861 | 0.5869 | 0 | 10 |

| Variable | R | p | p_FDR | Sig_FDR | N |
|---|---|---|---|---|---|
| tics_total_score | 0.328 | 0.1171 | 0.3746 | 0 | 25 |
| nmdas_cf_1 | -0.327 | 0.1187 | 0.3759 | 0 | 25 |
| nmdas_complete | -0.324 | 0.1224 | 0.3836 | 0 | 25 |
| nmdas_cca_3 | -0.319 | 0.1290 | 0.3931 | 0 | 25 |
| ln_nd1_copies_ul_serum_5 | -0.318 | 0.1595 | 0.4188 | 0 | 22 |
| cns_heel_wlk_tst | 0.318 | 0.1295 | 0.3931 | 0 | 25 |
| ln_nd1_copies_ul_serum_10 | -0.318 | 0.1601 | 0.4188 | 0 | 22 |
| adj_wgt_perc | -0.313 | 0.1366 | 0.3931 | 0 | 25 |
| cns_reg_wlk_tst | 0.313 | 0.1366 | 0.3931 | 0 | 25 |
| cns_superficial | 0.310 | 0.1404 | 0.3931 | 0 | 25 |
| cns_deep | 0.310 | 0.1404 | 0.3931 | 0 | 25 |
| cns_score_sensation | 0.310 | 0.1404 | 0.3931 | 0 | 25 |
| cns_kj_reflex_l | 0.310 | 0.1407 | 0.3931 | 0 | 25 |
| neo_open | 0.310 | 0.1409 | 0.3931 | 0 | 25 |
| insulin_serum | -0.306 | 0.2020 | 0.4724 | 0 | 20 |
| hr_avg_p5_60s_2 | -0.301 | 0.1740 | 0.4373 | 0 | 23 |
| cns_score_cerebellarfunction | 0.298 | 0.1579 | 0.4188 | 0 | 25 |
| lactate_nmolul__t_5 | -0.297 | 0.2465 | 0.5098 | 0 | 18 |
| fvc_predicted_average | 0.297 | 0.1694 | 0.4292 | 0 | 24 |
| cns_kj_reflex_r | 0.294 | 0.1639 | 0.4218 | 0 | 25 |
| ln_b2m_copies_ul_saliva_120 | -0.294 | 0.3304 | 0.5662 | 0 | 14 |
| met_feo2_d2 | -0.288 | 0.4888 | 0.6604 | 0 | 9 |
| co2_serum | 0.285 | 0.2240 | 0.4814 | 0 | 21 |
| bun_serum | -0.284 | 0.2245 | 0.4814 | 0 | 21 |
| pcl_total | 0.283 | 0.1795 | 0.4473 | 0 | 25 |
| nab_snl_spd_perc | 0.283 | 0.1904 | 0.4564 | 0 | 24 |
| hip_waist_circum_ratio | -0.281 | 0.1834 | 0.4518 | 0 | 25 |
| vf_c1_cf_cor_scaled | 0.280 | 0.1843 | 0.4518 | 0 | 25 |
| hr_avg_p5_60s_1 | -0.280 | 0.1949 | 0.4630 | 0 | 24 |
| nmdas_ssi_6 | -0.279 | 0.1867 | 0.4541 | 0 | 25 |
| ln_b2m_copies_ul_serum_90 | -0.278 | 0.2634 | 0.5138 | 0 | 19 |
| cns_skin | -0.277 | 0.1907 | 0.4564 | 0 | 25 |
| ssq_s | -0.274 | 0.2174 | 0.4814 | 0 | 23 |
| fgf21_fasting_pgml | -0.273 | 0.2587 | 0.5138 | 0 | 20 |
| nmdas_cca_2 | -0.271 | 0.2007 | 0.4724 | 0 | 25 |
| lactate_nmolul__t_60 | -0.271 | 0.2934 | 0.5301 | 0 | 18 |
| ln_b2m_copies_ul_plasma_5 | 0.269 | 0.2653 | 0.5138 | 0 | 20 |

| Variable | R | p | p_FDR | Sig_FDR | N |
|---|---|---|---|---|---|
| ln_nd1_copies_ul_serum_120 | -0.269 | 0.2654 | 0.5138 | 0 | 20 |
| lactate_nmolul__t_10 | -0.268 | 0.2982 | 0.5301 | 0 | 18 |
| cwi_inhib_scaled | 0.267 | 0.2077 | 0.4814 | 0 | 25 |
| st_free_corsort_scaled | 0.266 | 0.2573 | 0.5138 | 0 | 21 |
| cns_gait_statn | 0.266 | 0.2098 | 0.4814 | 0 | 25 |
| hr_avg_p10_60s_1 | -0.264 | 0.2243 | 0.4814 | 0 | 24 |
| ln_b2m_copies_ul_serum_120 | -0.262 | 0.2790 | 0.5183 | 0 | 20 |
| pfs_phys | -0.261 | 0.2173 | 0.4814 | 0 | 25 |
| ipaq_intensity | 0.260 | 0.2207 | 0.4814 | 0 | 25 |
| cns_other | 0.259 | 0.2210 | 0.4814 | 0 | 25 |
| cns_score_otherfindings | 0.259 | 0.2210 | 0.4814 | 0 | 25 |
| nmdas_cf_7 | -0.258 | 0.2237 | 0.4814 | 0 | 25 |
| dcf_sit_stand_time | 0.257 | 0.2249 | 0.4814 | 0 | 25 |
| nmdas_cf_9 | -0.256 | 0.2280 | 0.4847 | 0 | 25 |
| ln_nd1_copies_ul_serum_20 | -0.255 | 0.2775 | 0.5183 | 0 | 21 |
| eosabsolute_edta1 | 0.252 | 0.2704 | 0.5138 | 0 | 22 |
| ln_nd1_copies_ul_serum_30 | -0.250 | 0.2875 | 0.5265 | 0 | 21 |
| nmdas_cf_2 | -0.249 | 0.2403 | 0.5073 | 0 | 25 |
| ln_b2m_copies_ul_plasma_m5 | 0.248 | 0.2910 | 0.5297 | 0 | 21 |
| ctq_md | -0.247 | 0.2440 | 0.5081 | 0 | 25 |
| karnofsky_100 | 0.242 | 0.6005 | 0.7528 | 0 | 8 |
| hr_avg_p10_300s | -0.242 | 0.2652 | 0.5138 | 0 | 24 |
| ln_nd1_copies_ul_plasma_5 | 0.239 | 0.2960 | 0.5301 | 0 | 22 |
| ln_b2m_copies_ul_saliva_10 | -0.237 | 0.3600 | 0.5794 | 0 | 18 |
| dsm_5_detachment_imputed | 0.236 | 0.2674 | 0.5138 | 0 | 25 |
| cns_wj_reflex_l | 0.236 | 0.2679 | 0.5138 | 0 | 25 |
| cns_wj_reflex_r | 0.236 | 0.2679 | 0.5138 | 0 | 25 |
| cns_proximalbulk_r | 0.235 | 0.2689 | 0.5138 | 0 | 25 |
| cns_proximalbulk_l | 0.235 | 0.2689 | 0.5138 | 0 | 25 |
| nmdas_cf_8 | -0.235 | 0.2689 | 0.5138 | 0 | 25 |
| ln_nd1_copies_ul_plasma_60 | 0.233 | 0.3529 | 0.5794 | 0 | 19 |
| crp_serum | -0.232 | 0.3240 | 0.5629 | 0 | 21 |
| cns_alt_supin_pron | 0.231 | 0.2768 | 0.5183 | 0 | 25 |
| nmdas_ssi_2 | -0.230 | 0.2796 | 0.5183 | 0 | 25 |
| ln_b2m_copies_ul_plasma_10 | 0.229 | 0.3448 | 0.5794 | 0 | 20 |
| ln_b2m_copies_ul_plasma_120 | 0.224 | 0.3725 | 0.5794 | 0 | 19 |
| hr_avg_p10_60s_2 | -0.223 | 0.3059 | 0.5406 | 0 | 24 |

| Variable | R | p | p_FDR | Sig_FDR | N |
|---|---|---|---|---|---|
| fgf21_s3_pgml | -0.223 | 0.3315 | 0.5662 | 0 | 22 |
| ln_b2m_copies_ul_plasma_20 | 0.222 | 0.3600 | 0.5794 | 0 | 20 |
| cns_score_myotaticreflexes | 0.222 | 0.2968 | 0.5301 | 0 | 25 |
| lactate_nmolul__t_120 | -0.219 | 0.3993 | 0.5966 | 0 | 18 |
| cns_crnl_nrv_iv | 0.209 | 0.3259 | 0.5629 | 0 | 25 |
| cns_crnl_nrv_vi | 0.209 | 0.3259 | 0.5629 | 0 | 25 |
| chloride_serum | 0.208 | 0.3793 | 0.5794 | 0 | 21 |
| ln_b2m_copies_ul_saliva_90 | -0.203 | 0.4334 | 0.6202 | 0 | 18 |
| cns_skeletal | 0.201 | 0.3451 | 0.5794 | 0 | 25 |
| ln_b2m_copies_ul_serum_60 | -0.200 | 0.4125 | 0.6058 | 0 | 20 |
| albumin_serum | 0.198 | 0.4023 | 0.5966 | 0 | 21 |
| lactate_nmolul__t_m5 | -0.198 | 0.4468 | 0.6202 | 0 | 18 |
| mfis_phy_sub | -0.197 | 0.3569 | 0.5794 | 0 | 25 |
| ctq_sa | 0.195 | 0.3606 | 0.5794 | 0 | 25 |
| pi_geneticdiagnosistype | -0.193 | 0.3664 | 0.5794 | 0 | 25 |
| fgf21_s2_pgml | -0.191 | 0.4064 | 0.5998 | 0 | 22 |
| cns_nasopharynx | 0.189 | 0.3775 | 0.5794 | 0 | 25 |
| cns_crnl_nrv_v | 0.189 | 0.3775 | 0.5794 | 0 | 25 |
| cns_crnl_nrv_viii | 0.189 | 0.3775 | 0.5794 | 0 | 25 |
| cns_toe_wlk_tst | 0.189 | 0.3775 | 0.5794 | 0 | 25 |
| cns_romberg_tst | 0.189 | 0.3775 | 0.5794 | 0 | 25 |
| cns_fng_ns_fng | 0.189 | 0.3775 | 0.5794 | 0 | 25 |
| cns_left_toe | 0.189 | 0.3775 | 0.5794 | 0 | 25 |
| cns_score_toesign | 0.189 | 0.3775 | 0.5794 | 0 | 25 |
| nmdas_ssi_3 | -0.189 | 0.3775 | 0.5794 | 0 | 25 |
| nmdas_cca_8 | -0.189 | 0.3775 | 0.5794 | 0 | 25 |
| general_medical_and_neurological_examination_complete | -0.188 | 0.3788 | 0.5794 | 0 | 25 |
| ipaq_total_met | 0.185 | 0.3985 | 0.5966 | 0 | 24 |
| mbi_exhaus | 0.184 | 0.4003 | 0.5966 | 0 | 24 |
| creatinine_serum | 0.184 | 0.4379 | 0.6202 | 0 | 21 |
| ln_b2m_copies_ul_plasma_60 | 0.183 | 0.4674 | 0.6400 | 0 | 19 |
| leq_total_events_score | 0.180 | 0.3987 | 0.5966 | 0 | 25 |
| fgf21_s1_pgml | -0.176 | 0.4454 | 0.6202 | 0 | 22 |
| ln_nd1_copies_ul_saliva_m5 | -0.175 | 0.4871 | 0.6604 | 0 | 19 |
| cns_aj_reflex_l | 0.174 | 0.4165 | 0.6081 | 0 | 25 |
| cns_ocular_fundi | -0.173 | 0.4180 | 0.6081 | 0 | 25 |
| compass_orth_int | 0.169 | 0.4288 | 0.6185 | 0 | 25 |

| Variable | R | p | p_FDR | Sig_FDR | N |
|---|---|---|---|---|---|
| ctq_pn | 0.169 | 0.4293 | 0.6185 | 0 | 25 |
| hr_avg_p5_300s | -0.169 | 0.4419 | 0.6202 | 0 | 24 |
| potassium_serum | 0.168 | 0.4778 | 0.6514 | 0 | 21 |
| cns_macula_regn | 0.167 | 0.4362 | 0.6202 | 0 | 25 |
| cns_tj_reflex_l | 0.164 | 0.4441 | 0.6202 | 0 | 25 |
| cns_tj_reflex_r | 0.164 | 0.4441 | 0.6202 | 0 | 25 |
| dhs_score | 0.161 | 0.4523 | 0.6250 | 0 | 25 |
| dcf_bodyfat_avg | 0.157 | 0.4651 | 0.6398 | 0 | 25 |
| respr_count_p5_60s_1 | -0.154 | 0.5422 | 0.7208 | 0 | 19 |
| sodium_serum | 0.153 | 0.5209 | 0.6975 | 0 | 21 |
| ln_b2m_copies_ul_saliva_30 | -0.148 | 0.5988 | 0.7528 | 0 | 16 |
| ln_nd1_copies_ul_saliva_10 | -0.145 | 0.5533 | 0.7249 | 0 | 20 |
| scor_abp1_psqi_global | 0.143 | 0.5058 | 0.6804 | 0 | 25 |
| mbi_ineff | 0.133 | 0.5454 | 0.7208 | 0 | 24 |
| dsm_5_cross_cutting_imputed | 0.130 | 0.5448 | 0.7208 | 0 | 25 |
| respr_count_p10_60s_2 | -0.129 | 0.5867 | 0.7528 | 0 | 21 |
| mfis_cog_sub | 0.127 | 0.5534 | 0.7249 | 0 | 25 |
| nmdas_cf_5 | -0.127 | 0.5556 | 0.7249 | 0 | 25 |
| ln_b2m_copies_ul_saliva_m5 | -0.125 | 0.6852 | 0.8156 | 0 | 14 |
| ln_nd1_copies_ul_plasma_30 | 0.124 | 0.6017 | 0.7528 | 0 | 21 |
| compass_pupillomotor | -0.124 | 0.5631 | 0.7316 | 0 | 25 |
| ln_b2m_copies_ul_saliva_60 | -0.123 | 0.6268 | 0.7684 | 0 | 19 |
| leq_negative_events_score | 0.122 | 0.5896 | 0.7528 | 0 | 23 |
| mfis_psy_sub | 0.121 | 0.5723 | 0.7403 | 0 | 25 |
| dsm_5_psychoticism | -0.118 | 0.5925 | 0.7528 | 0 | 24 |
| cns_eye_mvt | 0.116 | 0.5909 | 0.7528 | 0 | 25 |
| rbans_ll_scaled | 0.114 | 0.5974 | 0.7528 | 0 | 25 |
| respr_count_p10_60s_1 | -0.112 | 0.6477 | 0.7875 | 0 | 20 |
| ln_b2m_copies_ul_plasma_90 | -0.111 | 0.6728 | 0.8084 | 0 | 18 |
| cns_aj_reflex_r | 0.108 | 0.6170 | 0.7674 | 0 | 25 |
| ln_nd1_copies_ul_saliva_20 | 0.106 | 0.6648 | 0.8052 | 0 | 20 |
| neo_neuro | -0.106 | 0.6214 | 0.7674 | 0 | 25 |
| ssq_n | 0.106 | 0.6230 | 0.7674 | 0 | 25 |
| nmdas_cf_6 | -0.106 | 0.6235 | 0.7674 | 0 | 25 |
| ln_nd1_copies_ul_plasma_20 | 0.105 | 0.6700 | 0.8083 | 0 | 20 |
| met_ree_d2 | -0.101 | 0.8128 | 0.9125 | 0 | 9 |
| stai_y1_state_total | -0.100 | 0.6411 | 0.7827 | 0 | 25 |

| Variable | R | p | p_FDR | Sig_FDR | N |
|---|---|---|---|---|---|
| ln_nd1_copies_ul_serum_90 | -0.093 | 0.7126 | 0.8364 | 0 | 19 |
| srh_score | 0.087 | 0.6847 | 0.8156 | 0 | 25 |
| cns_peripheral_rtn | -0.087 | 0.6868 | 0.8156 | 0 | 25 |
| pfs_mental | -0.085 | 0.6939 | 0.8208 | 0 | 25 |
| ln_nd1_copies_ul_saliva_30 | -0.083 | 0.7216 | 0.8405 | 0 | 22 |
| cns_ears | 0.079 | 0.7125 | 0.8364 | 0 | 25 |
| nmdas_ssi_5 | -0.078 | 0.7179 | 0.8393 | 0 | 25 |
| met_vo2_d2 | -0.074 | 0.8618 | 0.9299 | 0 | 9 |
| ln_b2m_copies_ul_plasma_30 | 0.071 | 0.7793 | 0.8907 | 0 | 19 |
| cns_crnl_nrv_iii | 0.071 | 0.7424 | 0.8614 | 0 | 25 |
| cns_bj_reflex_l | 0.065 | 0.7622 | 0.8810 | 0 | 25 |
| compass_vasomotor | -0.064 | 0.7659 | 0.8819 | 0 | 25 |
| cns_truncal_coord | 0.063 | 0.7700 | 0.8833 | 0 | 25 |
| compass_gastrointest | -0.057 | 0.7911 | 0.9007 | 0 | 25 |
| mbi_cyn | -0.056 | 0.8001 | 0.9076 | 0 | 24 |
| nab_sdgb_perc | 0.054 | 0.8051 | 0.9099 | 0 | 24 |
| fibrinogen_blue | -0.052 | 0.8274 | 0.9219 | 0 | 21 |
| mcc_score | -0.051 | 0.8134 | 0.9125 | 0 | 25 |
| ln_nd1_copies_ul_saliva_60 | 0.049 | 0.8410 | 0.9238 | 0 | 20 |
| ln_nd1_copies_ul_plasma_90 | -0.049 | 0.8418 | 0.9238 | 0 | 20 |
| nab_shl_irg_perc | 0.047 | 0.8305 | 0.9219 | 0 | 24 |
| nmdas_ssi_8 | -0.046 | 0.8309 | 0.9219 | 0 | 25 |
| bdi_total | 0.045 | 0.8364 | 0.9238 | 0 | 25 |
| nmdas_ssi_9 | -0.042 | 0.8466 | 0.9257 | 0 | 25 |
| ln_nd1_copies_ul_saliva_90 | 0.040 | 0.8713 | 0.9360 | 0 | 20 |
| cns_crnl_nrv_ii | 0.039 | 0.8548 | 0.9296 | 0 | 25 |
| cns_score_generalmedical | 0.039 | 0.8562 | 0.9296 | 0 | 25 |
| neo_cons | -0.037 | 0.8626 | 0.9299 | 0 | 25 |
| ln_nd1_copies_ul_plasma_10 | -0.037 | 0.8763 | 0.9380 | 0 | 21 |
| cns_score_ocularfundi | -0.033 | 0.8799 | 0.9385 | 0 | 25 |
| cns_eyes | -0.031 | 0.8861 | 0.9419 | 0 | 25 |
| compass_total | -0.026 | 0.9043 | 0.9546 | 0 | 25 |
| pss_total | -0.026 | 0.9044 | 0.9546 | 0 | 25 |
| calcium_serum | -0.025 | 0.9169 | 0.9645 | 0 | 21 |
| ln_nd1_copies_ul_saliva_120 | -0.024 | 0.9224 | 0.9669 | 0 | 20 |
| scr_max_sc_5_min | -0.019 | 0.9360 | 0.9778 | 0 | 22 |
| scr_max_sc_10_min | -0.016 | 0.9488 | 0.9811 | 0 | 20 |

| Variable | R | p | p_FDR | Sig_FDR | N |
|---|---|---|---|---|---|
| nmdas_ssi_7 | 0.015 | 0.9439 | 0.9809 | 0 | 25 |
| nab_sdgf_perc | 0.015 | 0.9454 | 0.9809 | 0 | 24 |
| ln_nd1_copies_ul_plasma_120 | 0.014 | 0.9542 | 0.9816 | 0 | 20 |
| compass_bladded | -0.011 | 0.9580 | 0.9816 | 0 | 25 |
| cns_bj_reflex_r | 0.011 | 0.9590 | 0.9816 | 0 | 25 |
| stai_y1_trait_total | 0.010 | 0.9628 | 0.9822 | 0 | 25 |
| ln_nd1_copies_ul_plasma_m5 | 0.009 | 0.9708 | 0.9825 | 0 | 22 |
| ln_b2m_copies_ul_saliva_20 | -0.008 | 0.9756 | 0.9825 | 0 | 19 |
| ln_nd1_copies_ul_saliva_5 | -0.007 | 0.9760 | 0.9825 | 0 | 21 |
| cns_abdomen | -0.007 | 0.9735 | 0.9825 | 0 | 25 |
| hgba1c_edta1 | 0.004 | 0.9857 | 0.9889 | 0 | 22 |
| neo_agree | -0.002 | 0.9908 | 0.9908 | 0 | 25 |

Table S4. Sensitivity of the NMDAS–brain association to additional covariates, related to Figure 3. Baseline: NMDAS vs Brain r = −0.670, p = 0.0003 (controlling age only, n = 25 patients). Each row shows the adjusted partial correlation after additionally controlling for the listed confound. Bootstrap test (5,000 resamples) assesses whether the decrease in |r| is statistically significant. Sig_Decrease: TRUE = 95% bootstrap CI excludes zero.

| Covariate | ρ with outcome | p | Adjusted ρ | Adjusted p | n |
|---|---|---|---|---|---|
| cns_eyes (CNS Eyes exam) | 0.118 | 0.5832 | -0.769 | 0.0000 | 25 |
| cns_crnl_nrv_ii (Cranial nerve II) | 0.168 | 0.4335 | -0.660 | 0.0006 | 25 |
| cns_crnl_nrv_iii (Cranial nerve III) | 0.275 | 0.1932 | -0.725 | 0.0001 | 25 |
| cns_crnl_nrv_iv (Cranial nerve IV) | 0.323 | 0.1240 | -0.684 | 0.0003 | 25 |
| cns_crnl_nrv_vi (Cranial nerve VI) | 0.323 | 0.1240 | -0.684 | 0.0003 | 25 |
| pfs_phys (Physical fatigue) | -0.563 | 0.0042 | -0.441 | 0.0352 | 25 |
| pfs_mental (Mental fatigue) | -0.432 | 0.0348 | -0.576 | 0.0040 | 25 |
| cns_gait_statn (Gait and station) | 0.430 | 0.0359 | -0.573 | 0.0043 | 25 |
| nmdas_ssi_1 (Psychiatric symptoms) | -0.461 | 0.0233 | -0.554 | 0.0061 | 25 |
| Covariate | Δ\|ρ\| | CI lower | CI upper | Bootstrap p | Decrease |
| cns_eyes (CNS Eyes exam) | -0.099 | -0.324 | 0.089 | 1.000 | FALSE |
| cns_crnl_nrv_ii (Cranial nerve II) | 0.010 | -0.061 | 0.123 | 0.702 | FALSE |
| cns_crnl_nrv_iii (Cranial nerve III) | -0.054 | -0.319 | 0.195 | 1.000 | FALSE |
| cns_crnl_nrv_iv (Cranial nerve IV) | -0.014 | -0.280 | 0.248 | 1.000 | FALSE |
| cns_crnl_nrv_vi (Cranial nerve VI) | -0.014 | -0.279 | 0.242 | 1.000 | FALSE |
| pfs_phys (Physical fatigue) | 0.229 | -0.103 | 0.527 | 0.163 | FALSE |
| pfs_mental (Mental fatigue) | 0.094 | -0.068 | 0.384 | 0.240 | FALSE |
| cns_gait_statn (Gait and station) | 0.098 | -0.035 | 0.309 | 0.179 | FALSE |
| nmdas_ssi_1 (Psychiatric symptoms) | 0.117 | -0.052 | 0.315 | 0.154 | FALSE |

Adjusted correlations additionally control for the listed covariate and age. Bootstrap quantities and all reported precision are retained from the source table; 0.0000 denotes rounding, not an exact zero probability.

Table S5. Sensitivity of the NMDAS–behavior association to additional covariates, related to Figure 3. Baseline: NMDAS vs Behavior r = −0.460, p = 0.024 (controlling age only, n = 25 patients). Each row shows the adjusted partial correlation after additionally controlling for the listed confound. Bootstrap test (5,000 resamples) assesses whether the decrease in |r| is statistically significant. Sig_Decrease: TRUE = 95% bootstrap CI excludes zero.

| Covariate | ρ with outcome | p | Adjusted ρ | Adjusted p | n |
|---|---|---|---|---|---|
| cns_eyes (CNS Eyes exam) | -0.031 | 0.8861 | -0.613 | 0.0019 | 25 |
| cns_crnl_nrv_ii (Cranial nerve II) | 0.039 | 0.8548 | -0.460 | 0.0272 | 25 |
| cns_crnl_nrv_iii (Cranial nerve III) | 0.071 | 0.7424 | -0.610 | 0.0020 | 25 |
| cns_crnl_nrv_iv (Cranial nerve IV) | 0.209 | 0.3259 | -0.468 | 0.0243 | 25 |
| cns_crnl_nrv_vi (Cranial nerve VI) | 0.209 | 0.3259 | -0.468 | 0.0243 | 25 |
| pfs_phys (Physical fatigue) | -0.261 | 0.2173 | -0.450 | 0.0313 | 25 |
| pfs_mental (Mental fatigue) | -0.085 | 0.6939 | -0.492 | 0.0171 | 25 |
| cns_gait_statn (Gait and station) | 0.266 | 0.2098 | -0.389 | 0.0665 | 25 |
| nmdas_ssi_1 (Psychiatric symptoms) | -0.346 | 0.0978 | -0.335 | 0.1182 | 25 |
| Covariate | Δ\|ρ\| | CI lower | CI upper | Bootstrap p | Decrease |
| cns_eyes (CNS Eyes exam) | -0.154 | -0.411 | 0.129 | 1.000 | FALSE |
| cns_crnl_nrv_ii (Cranial nerve II) | -0.001 | -0.110 | 0.128 | 1.000 | FALSE |
| cns_crnl_nrv_iii (Cranial nerve III) | -0.151 | -0.421 | 0.214 | 1.000 | FALSE |
| cns_crnl_nrv_iv (Cranial nerve IV) | -0.008 | -0.295 | 0.383 | 1.000 | FALSE |
| cns_crnl_nrv_vi (Cranial nerve VI) | -0.008 | -0.299 | 0.386 | 1.000 | FALSE |
| pfs_phys (Physical fatigue) | 0.010 | -0.327 | 0.343 | 0.921 | FALSE |
| pfs_mental (Mental fatigue) | -0.033 | -0.262 | 0.200 | 1.000 | FALSE |
| cns_gait_statn (Gait and station) | 0.071 | -0.112 | 0.300 | 0.466 | FALSE |
| nmdas_ssi_1 (Psychiatric symptoms) | 0.125 | -0.112 | 0.379 | 0.305 | FALSE |

Adjusted correlations additionally control for the listed covariate and age. Bootstrap quantities and all reported precision are retained from the source table; 0.0000 denotes rounding, not an exact zero probability.
